# Effects of artificial light colour, intensity, structure and contrast on moth flight behaviour

**DOI:** 10.64898/2026.04.15.718644

**Authors:** Emmanuelle S. Briolat, James A. M. Galloway, Menno van Berkel, Jonathan Bennie, Kevin J. Gaston, Jolyon Troscianko

## Abstract

Nocturnal moths are severely affected by light pollution, most notoriously through fatal attraction to artificial lights, yet flight-to-light is not their only response. To investigate how artificial lights impact flight behaviour, we exposed over 1200 wild-caught moths of 62 species to LED lights with different characteristics, under varying background lighting conditions, and tracked over 500 flight paths in three dimensions. Flight-to-light behaviour and flight tortuosity both increased with light intensity, irrespective of spectrum, though tortuosity was affected by lower levels of white than amber light, suggesting white LEDs could impact moth trajectories from greater distances. Flight tortuosity was also higher upon exposure to a single light versus three producing equivalent illuminance. Conversely, higher background light levels led to reductions in both flight-to-light and tortuosity, but moths were also less likely to take flight in these conditions, suggesting that both point sources and diffuse background lighting disrupt moth movement. Finally, moths caught using light traps were less likely to fly and, if they did, more likely to fly towards light sources than those caught with butterfly nets. These findings suggest mitigation policies for light pollution should prioritize reducing light intensity, and point to new directions for future research.

## Introduction

Artificial light at night is a growing source of anthropogenic pollution, reshaping fundamental natural light cycles and the nocturnal light environment on a global scale, with consequences across the tree of life, at all levels from individual physiology to ecosystem function [1–3]. Impacts on nocturnal insects are particularly well-documented [4], including suppression of activity, feeding and reproduction [5–8], disruption of movement and migration [9], and ultimately contributions to population declines [10,11]. Flight-to-light, or positive phototaxis, is the best-known effect of exposure to artificial lights for nocturnal moths (Lepidoptera) and other insects [12], causing both direct mortality and other harms, such as increased predation risk [13] and distraction from essential activities [14]. Why moths are so attracted to lights has been an enduring puzzle, but recent research showed that a simple rule of dorsal orientation to light can explain the movements of individuals around lights at close range [15]. Yet in the field, insects display a variety of other responses to lights, such as freezing in place, flying upwards, downwards or erratically [16]. Experiments suggest only a small proportion of moths released near light sources will end up being attracted to them [17–19]. Instead, disruptions from artificial lights can take other forms, such as increased flight tortuosity, or “barrier” effects preventing insects from leaving lit areas, suggesting that measuring only the attractiveness of lights may underestimate their ecological impacts [18].

In the context of widespread insect declines [20], it is crucial to understand both how artificial lights disrupt insect movements, and how these harms might best be mitigated by changing different properties of lighting systems [21]. Research has largely focused on the effects of different colours, or spectra of light, on insect movements. This is of particular concern as traditional light types (e.g., gas discharge) are progressively replaced with light-emitting diodes (LEDs), that typically emit whiter light, rich in short wavelengths, which most species are especially sensitive to [22,23]. Amber lights are generally less attractive to insects [24], and hence perceived as a more ecologically-sensitive alternative, particularly phosphor-converted (PC) amber LEDs which still support some colour discrimination by humans [25]. Yet their impacts vary widely across taxa and behavioural contexts [25,26], and other potential mitigation strategies remain relatively underexplored [21] (but see [27–29]).

Here, we investigated how exposure to artificial lights with different characteristics affects where moths fly, and the shape of their flight paths, across a wide range of nocturnal species. After a period of acclimation to ambient light levels, wild-caught moths collected either with light traps or butterfly nets were individually released on the floor of an indoor flight arena. As they flew upwards, they passed through a beam of light created by a focal LED light, or lights, and flights were recorded with two cameras to enable reconstruction of three-dimensional tracks (Figure 1; Supplementary Video 1). Through two experiments, repeated in the summers of 2024 and 2025, we tested the impacts of varying light spectrum and intensity, ranging from skyglow levels to conditions below streetlights (Experiment 1), as well as different focal light structures (single LED at 30 lx, versus three at 10 lx each) and background light levels (Experiment 2; Figure 1).

**Figure 1:**
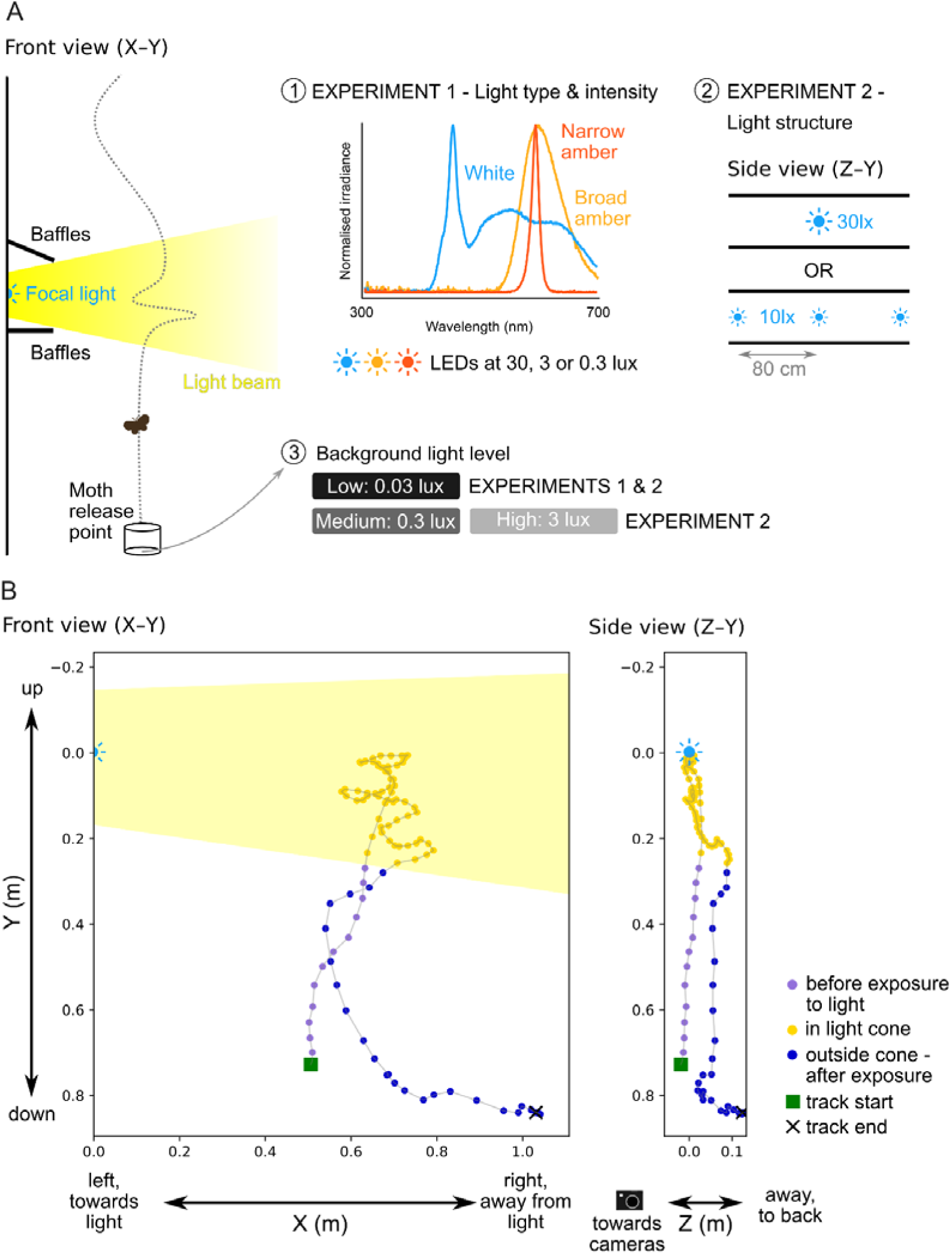
Tracking moth flights in response to light exposure. **(A)** Schematic of the experimental set-up, as viewed by the cameras (not to scale). Insets 1, 2 and 3 respectively provide details of light types, or spectra, and intensity treatments in experiment 1, light structures in experiment 2, and background light levels in both experiments. **(B)** Orthogonal projections of a reconstructed 3D flight in this system. Yellow shading in the front view represents the approximate position of the light beam. See also Supplementary Video S1.

## Methods

### Experimental design

This study was designed to test how a wide range of species of wild-caught moths would respond when exposed mid-flight to LED lights with different characteristics. Experiments with live moths were approved by the University of XXXX’s ethics committee (application ID: 1146374), and took place during two field seasons, from 30th July to 16th September 2024 and 2nd July to 27th September 2025. Moths from 62 species in 6 different families were collected in the grounds of the XXXX campus (XXXX) and nearby private gardens, using both light traps (with 125W or 80W mercury-vapor lamps, or a 15W actinic light) and nighttime collections with butterfly nets (Supplementary Table S1). Trapped moths were collected in the morning and housed individually in small plastic pots with air-holes, outdoors in a shaded location until the evening; hand-netted moths were placed in the same pots but were used in experiments within an hour of capture. Flight experiments took place indoors in a section of a dark room (130 x 280 x 208 high, in cm), screened off by opaque black curtains and black mesh, to contain the moths once they had flown. All moths were given at least 30 minutes to acclimate to the light conditions in each experiment before trials began, corresponding to estimates of the time taken for pigment migration in the moth eye for adaptation to ambient light conditions [30,31]. Moths were released individually, at floor level, by an experimenter sliding off the pot lid. Above them, single LEDs and baffles made of black card and ethylene-vinyl acetate (EVA) foam were used to produce a cone of light through which the moths passed as they flew upwards (Figure 1). Each trial lasted one minute, before moths that did fly were recaptured by the experimenters. Flight trials were carried out between 20:30 and midnight, in sessions lasting up to two hours. Moths were then released in the same areas they were captured in.

Flights were filmed from the side by an infrared (IR) stereo camera system. The system used a Raspberry Pi 5 board with two IR cameras attached to the main board, spaced 60cm apart, and mounted on an aluminium section pole. Custom code in Python and OpenCV was used to calibrate the two cameras using a series of images of a chess board. This built up a dataset used to create matrix transforms that account for lens distortion, rotation and translation, allowing any points visible in both cameras to have their 3D position calculated. Videos were recorded using a custom camera user interface to operate both cameras simultaneously, filming at 15 frames per second, with 1280x720 px resolution.

### Experiment 1

The first experiment was designed to investigate the behaviour of moths flying past LEDs of different spectra and intensity. We tested the effects of a standard white LED (correlated colour temperature (CCT) = 5986 K), a broadband phosphor-converted (PC) amber LED (CCT = 1743 K) and a narrowband amber LED (CCT = 1621 K) [8], as well as control conditions with no test lights. Each of the LEDs was calibrated in a dark room to produce an illuminance of either 30 lx, 3 lx or 0.3 lx at a distance of 50 cm, directly above the moth release point, using an open-source mini spectroradiometer with a sensitivity of 0.005 lx [32]. The light level experienced by moths at the release point was maintained at 0.03 - 0.05 lx across all treatments, as measured by the same spectroradiometer, matching conditions in similar flight experiments with hawkmoths [14]. In the treatments with an illuminance of 30 lx, this was achieved by bleed-through from the test lights; for the lower illuminance treatments, an additional white LED was turned on beyond the blackout curtains and pointed at the room’s silver ceiling, to create diffuse lighting of the same intensity.

A total of 806 moths of 55 known species in 6 families were tested in this experiment. In 2024, trials included only control conditions and all three LED types at the highest intensity; in 2025, all lower light levels were tested, alongside repeats of the highest intensity and control treatments (Supplementary Table S2 for sample sizes). Each moth was used in a single flight trial; within each year, blocks of each treatment were pseudo-randomized within and between experimental sessions, to try to balance the number of flights recorded per treatment. In 2024, trials in experiment 1 began with a single camera set-up (not allowing 3D tracking of the flight path), before switching to a stereo camera system; all trials in 2025 (and experiment 2) used the stereo camera system. Trials filmed with only a single camera were included in analyses of take-off probability and flight outcomes, but were then excluded from subsequent analyses of flight trajectories. Five trials were excluded from the dataset due to experimenter error, equipment failure or damage to the moth, yielding a total of N_1_ = 801 trials for analysis.

### Experiment 2

The second experiment used only a single light type, the white LED from experiment 1, to test the effects of both different structures of light, and varying contrasts between background light levels and the focal lights. In this set-up, the cone of light was produced either by a single LED, calibrated as above to 30 lx at a distance of 50 cm, or three single LEDs, positioned 80cm apart and each calibrated to produce an illuminance of 10 lx at the same distance (“single” and “multiple” lights respectively). Additional white LEDs pointing at the ceiling were also used to produce three levels of background lighting directly above the moth release point (approximately 0.03 lx, 0.3 lx and 3 lx), resulting in six different combinations of test lighting structure and background lighting levels. To achieve the highest ambient light level of 3 lx, two white LEDs were attached to the ceiling of the experimental chamber, screened by a thin polytetrafluoroethylene (PTFE) diffuser.

A total of 462 moths of 32 species in 5 families were tested in this experiment, with 46 – 164 trials per treatment combination. In 2024, background light levels of 0.03 and 0.3 lx were tested; in 2025, the 3 lx background treatment was added, alongside repeats of the 0.03 lx treatment (see Supplementary Table S2 for sample sizes). For this experiment, each moth was used in two trials, with different focal light treatments under the same background light level. Each experimental session featured a single background light level, pseudo-randomized to balance flight numbers in each treatment, and the order of testing the single or multiple lights was randomized between sessions. However, the number of trials run with single and multiple focal lights are not balanced for the medium and high background light conditions, due to additional treatments that were later excluded from the main analyses. First, in 2024, 47 moths tested with the multiple focal light treatment under medium background lighting were tested in an additional condition, with a single central light producing 10 lx illuminance. These trials were compared to the same moths’ trials with the multiple 10 lx light treatment, and to trials from the same year, under the same background lighting, with a single 30 lx light, to disentangle the impacts of altering the overall intensity of lighting, by changing either the number or intensity of lights; no significant differences were found between these treatments, whether in terms of the probability of successful flight, flight-to-light and erratic spiralling behaviour, or flight tortuosity (see Supplementary Figure S1). This suggested that a difference in overall focal light intensity from 10 to 30 lx was insufficient to alter moth behaviour in our set-up, so the single 10 lx treatment was not repeated in 2025, and trials in this treatment were excluded from the main dataset. Second, in 2025, 24 moths in the single light with high background lighting condition were paired with trials under high background lighting, but with no focal light; these trials were excluded from the main dataset, but were compared to control flights under low background light levels in the same year, to further investigate the effect of background lighting only (Supplementary Figure S2). From a total of 853 relevant trials in experiment 2, 21 were excluded as for experiment 1, yielding a final number of N_2_ = 832 trials for analysis.

### Video analysis

Flight tracks were analysed using custom-built Python scripts. To make the flying moths easier to see, each video was first processed to convert motion information into false colour images, following methods developed for the BehaveAI motion detection and classification framework [33]. The position of the moth was then recorded by clicking the front edge of the motion track in every frame, from the moment each moth took off until it either reached the focal light, landed, or exited the filming area. If a moth clearly landed or exited before crossing the path of the focal light beam, the rest of the minute of footage per trial was watched, in case the moth took off again or returned into view. If so, either the first flight segment in which it crossed the path of the light, or the longest flight segment, was selected for tracking. Reference videos taken before every experimental session, or whenever the set-up was moved, allowed us to record the position of the focal lights (or central focal light when multiple lights were used), as well as the position of the light beam, for each video clip.

Each moth position along the tracked flights could therefore be labelled as falling in or out of the light beam, so tracks in which moths never crossed into the light could be objectively identified. Tracks with fewer than five points, or which did not cross the light beam, were excluded from analyses of flight outcomes and tortuosity; for flights filmed in 3D with the stereo camera system, these criteria applied to tracks from both cameras. To map tracks in 3D, moth trajectories were tracked independently for each camera, trimmed to the same start and end points, then combined to produce new three-dimensional coordinates, based on the calibration information of the camera system. Tracks were then smoothed with a Savitzky-Golay filter [34,35], with a window of 5 and a second-order polynomial.

Outcomes of the tracked flights were scored manually by experimenters, semi-blind to treatment (the presence or absence of a focal light could not be concealed, although its spectrum and the presence of background light could not be determined). Tracks were considered to end at the focal light if a moth either landed on the light apparatus (including baffles funnelling the light beam) or its track disappeared into the glare of the light. Other possible flight outcomes recorded included moths returning to the floor, landing elsewhere (such as the curtains and walls delimiting the flight arena), and exiting the frame. Erratic spiralling behaviour was defined as detrimental spiralling motion that substantially interrupted the moth’s flight path, specifically as either spiralling downwards, or performing three or more consecutive spirals in any other direction; this was scored over the full length of each trial. Once a moth track disappeared into the glare of the focal light, further movement around the LED, including spiralling around the light, was not tracked or considered for scoring as erratic spiralling, as movement near the light was artificially constrained by the presence of the baffles.

Quantitative measures of flight tortuosity were taken as the mean and standard deviation of turning angles, calculated from the 3D coordinates of the tracked flights. To test how flight behaviour changed within trials, after moths were exposed to the focal light, these metrics were also calculated separately for track segments before and after light exposure. Moths whose flight paths intersected with the light beam but were flying directly away from it were unlikely to have seen the focal light or to have experienced any effect from it; to more accurately account for these scenarios, exposure to light was scored according to a combination of criteria. For each tracked moth, the post-exposure flight segment began from the first position in which its coordinates fell within the light beam, and it was deemed able to see the light, as defined by a bearing angle to the light of -135° to 135° (where an angle of 0° represents flight directly towards the light). Only tracks with both pre- and post-exposure segments longer than five points were included in analyses of the effects of light exposure on flight tortuosity.

### Statistical analysis

All statistical analyses were carried out in R version 4.3.2 [36]. For experiment 1, data from both years were combined for all analyses, as preliminary tests restricted to treatments repeated in both 2024 and 2025 found no effect of year on any variable of interest. The probability of moths taking off, flying to light or flying upwards by the end of their tracked flights, performing spirals or erratic spiralling behaviour, were initially analysed across all treatments using binomial mixed effects models built with the package glmmTMB [37], with treatment and capture method as fixed effects, and moth species and family as random effects. When a significant effect of treatment was found, secondary analyses were carried out without control flights, breaking down treatment into interacting variables representing light intensity and spectrum, to tease apart the effects of these different properties of the focal light. When modelling the effect of treatment on the probability of flight-to-light, the treatment level with the broadband amber focal LED at 0.3 lx was dropped from the analysis, as no flight-to-light was observed in that condition. Quantitative measures of flight tortuosity, taken as the mean and standard deviation of turning angles across whole flights, were analysed using equivalent linear mixed effects models with focal light treatment and capture method as fixed effects, and moth species and family as random effects; the dependent variables were square root transformed when necessary to fit model assumptions. Similarly to flight outcomes, when a significant effect of treatment was found, further analyses were run for the lit treatments only, separating out the interacting effects of light spectrum and intensity. When analysing tortuosity metrics across flight segments before and after moths were exposed to the focal light, treatment was allowed to interact with a binary variable representing exposure to light, and an additional random effect for trial ID was added. Models for the standard deviation of turning angle before and after light exposure did not converge when all random effects were included, so phylogeny was only accounted for at the family level; model comparisons using Akaike’s Information Criterion (AIC [38]) were used to compare models including species or family. All model assumptions were checked using the *simulateResiduals* function in the package DHARMa [39], and quantitative metrics of tortuosity were square-root or log-transformed as necessary to fit assumptions. In all cases, stepwise model simplification was carried out using likelihood ratio tests, and planned comparisons with Tukey’s post-hoc tests, using the packages multcomp [40] and emmeans [41].

Data from experiment 2 were analysed as above, with an additional fixed effect for trial number, and random effect for moth ID, to account for repeated flights by the same individuals. As there was no unlit control in this experiment, treatment was always included as interacting variables for background light level and focal light structure. Preliminary analyses restricted to treatments repeated in both years (with low background light level) found a higher overall probability of observing spirals in flight and flight-to-light, as well as higher mean turning angles, in 2025 compared to 2024, possibly due to differences in the construction of the baffles funnelling the light beam. These metrics were thus analysed separately for each year. Models of mean turning angle in 2024 and standard deviation of turning angle across both years, over complete flight tracks, could not converge with all random effects included for family, species and moth ID; models with different combinations of random effects were compared using AIC [38] to identify the best model structure. When modelling tortuosity metrics in flight segments before and after exposure to light, moth ID was replaced by a random effect for trial ID. For mean turning angles before and after exposure in 2024, models did not converge with all random effects; trial ID was prioritised so as to account for repeated measures of each flight path, but results did not change if the best-fitting model was used instead (based on AIC, with species and family but no trial ID). Supplementary analyses with additional treatments excluded from the main dataset were carried out using the same protocols (see Supplementary Figures S1, S2).

## Results

### Probability of moths taking flight under different lighting conditions

Overall, moths took flight within one minute of their introduction to the experimental arena in 49% (394/801) and 35% (295/834) of trials in experiments 1 and 2 respectively. Varying background lighting levels affected flight probability: in experiment 2, moths were significantly less likely to take off under the highest background light level tested ( ≈ 3 lx), with odds of a moth taking off over two thirds lower than under both low and medium background lighting conditions (generalized linear mixed effects model (GLMM), N = 834, X^2^_2_ = 25.23, p < 0.001; Tukey contrasts, p_high-medium_ < 0.001, p_high-low_ <0.001, p_low-medium_ = 0.998, ; GLMM, estimate = -1.251, odds ratio (OR)= 0.286, and estimate = -1.233, OR = 0.291, for high compared to low and medium levels respectively; Figure 2B,C). This effect was supported by additional analyses of trials with no focal light but different background light intensities (see Supplementary Figure S2). Moths in experiment 2 were also used in repeated trials, typically 30 mins to an hour apart, and were half as likely to fly in their second trial compared to their first (GLMM, Trial number: X^2^_1_ = 15.152, p < 0.001; estimate for trial 2 = -0.699, OR = 0.497; Figure 2D). Finally, in both experiments, moths collected with light traps were only half as likely to fly as those caught by hand with butterfly nets (GLMM, N = 801, X^2^_1_ = 16.128, p < 0.001, estimate for traps =-0.700, OR = 0.497 in experiment 1, and X^2^_1_ = 4.732, p = 0.0296, estimate for traps = -0.550, OR = 0.577 in experiment 2; Figure 2E,F; Supplementary Figure S3). There was no effect of any focal light variable on flight probability, whether light treatments in experiment 1 (GLMM, X^2^_9_ = 14.974, p = 0.0916; Figure 2A), or structure (single or multiple) in experiment 2 (GLMM, Light structure * Background light level, X^2^_2_ = 5.660, p = 0.059; Light structure: X^2^_1_ = 0.122, p = 0.727), suggesting that moths were effectively shielded from perceiving the focal light(s) at the point of release.

**Figure 2:**
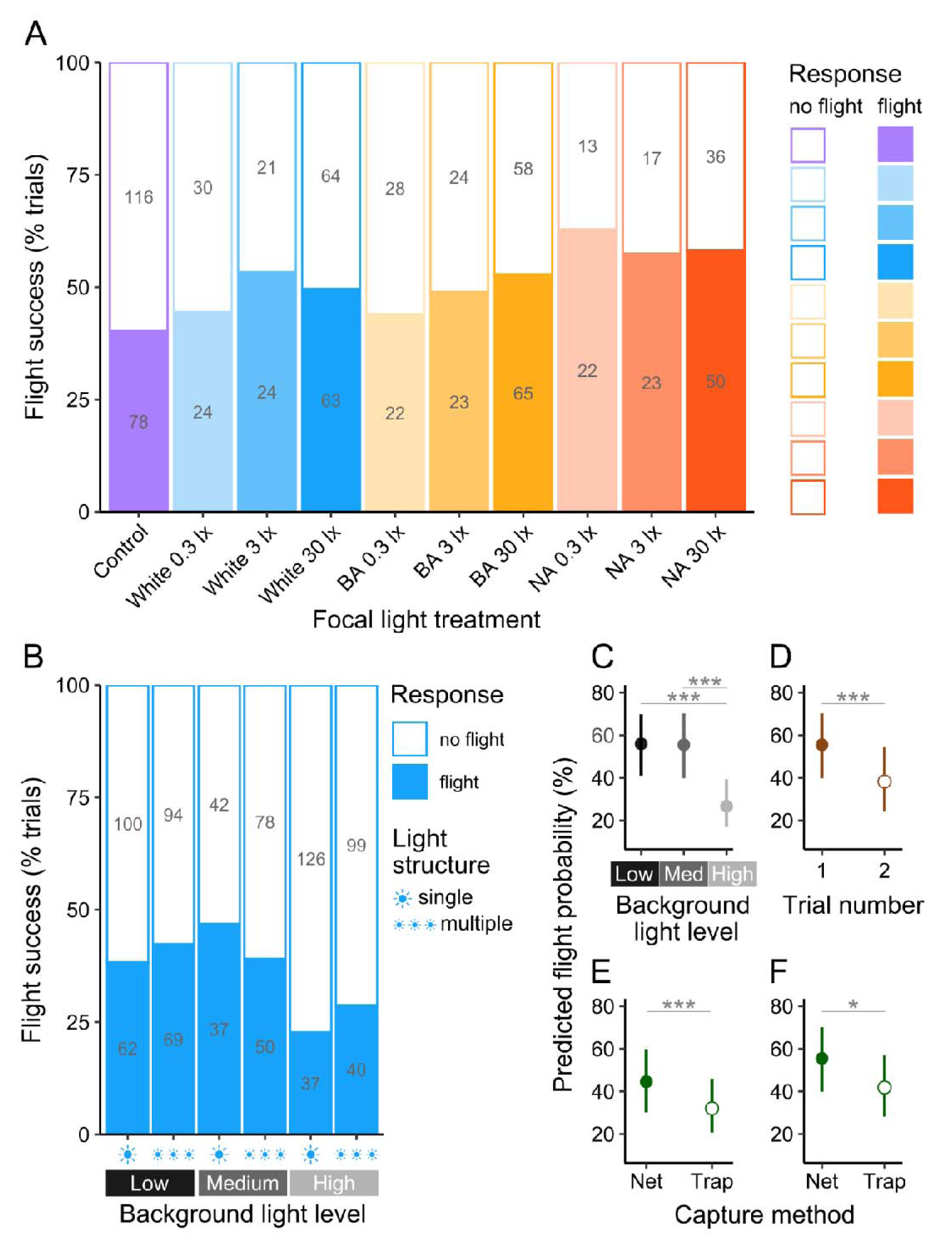
Flight probability. **(A)** Percentage of trials with take-offs per focal light intensity and spectrum, in experiment 1. **(B)** Percentage of trials with take-offs per combination of focal light structure and background lighting in experiment 2. In A & B, values in grey indicate raw numbers per group. **(C-D)** Predicted flight probability in experiment 2 according to background light level **(C)** and trial number **(D)**. **(E-F)** Predicted flight probability for moths collected with light traps or nets, in experiments 1 **(E)** and 2 **(F)**. In **(D-F)**, dots represent predicted probabilities for fixed effects only, and whiskers 95% confidence intervals. Asterisks show significant differences: * = p < 0.05, ** = p < 0.01, ***: p < 0.001.

### Flight-to-light behaviour

Flight paths with fewer than 5 points, or where the moth did not enter the light beam, were excluded, resulting in 302 flights for further analysis in experiment 1 (18-57 per treatment) and 264 (32-58 per treatment) in experiment 2 (Supplementary Table S2). Flight paths typically ended when moths flew up and out of frame (in 60% and 52% of trials in experiments 1 and 2 respectively), or to the light apparatus (24% and 31% resp.). A small proportion flew down, to the floor or out of frame (7% and 8% resp.), while the remainder flew out of the flight arena sideways or towards the camera, or landed on the sides of the arena (Figure 3A-E).

**Figure 3:**
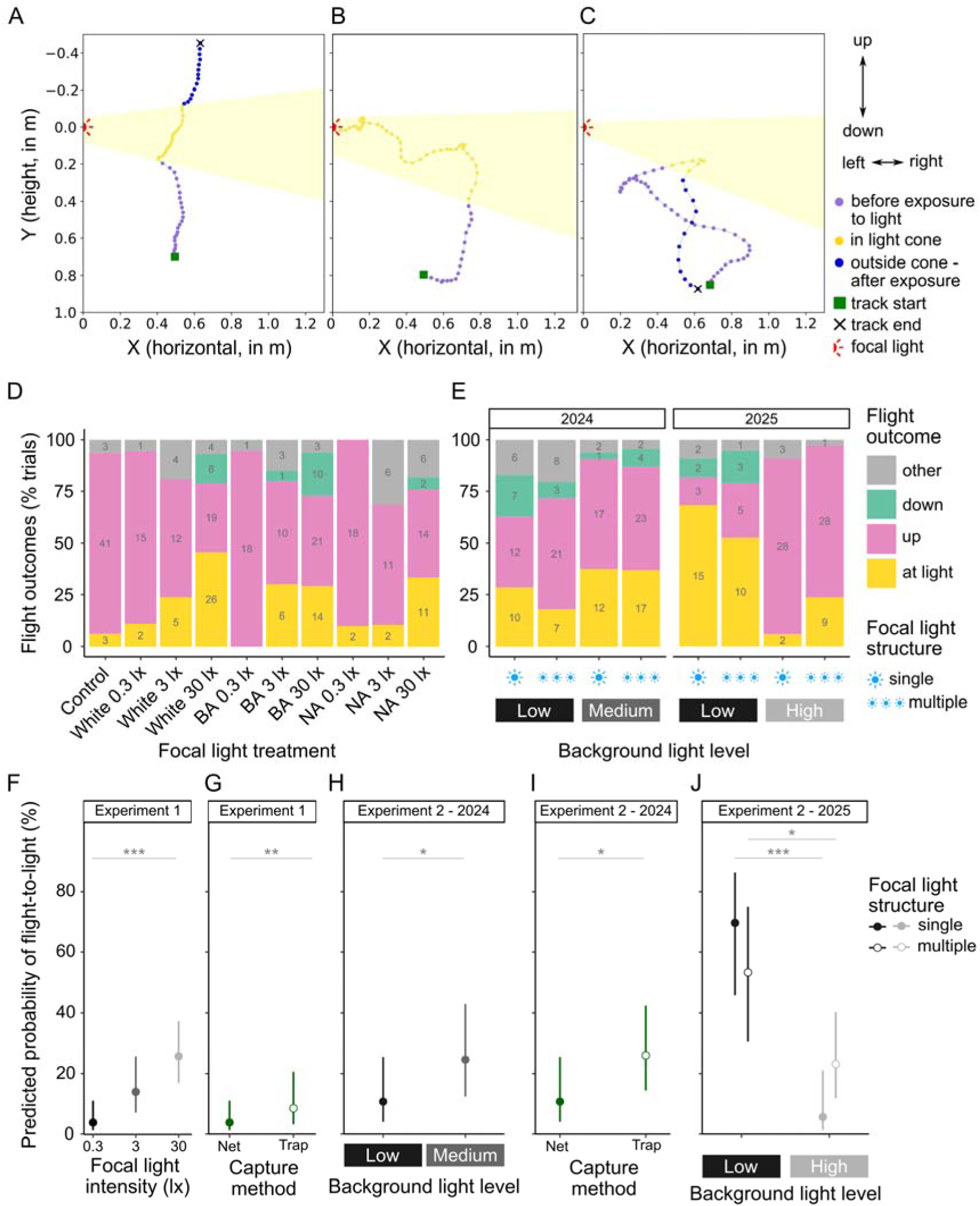
Flight outcomes. (A-C) Orthogonal projections in XY (camera view) for example tracks in which moths flew up **(A)**, to light **(B)** and down **(C)**. Shading indicates approximate position of the light beam. **(D-E)** Outcomes of tracked flights in experiments 1 **(D)** and 2 **(E)**; values in grey represent raw numbers of moths. **(F-J)** Probability of flight-to-light, as predicted by the best statistical models for experiment 1 **(F-G)** and 2 **(H-J)**. Dots represent predicted probabilities for fixed effects only, and whiskers 95% confidence intervals. Asterisks show significance levels: * = p < 0.05, ** = p < 0.01, ***: p < 0.001.

Across both experiments, flight outcomes were shaped by the intensity of both the focal light and background lighting, as well as collection method. In experiment 1, the probability of flight-to-light differed between treatments (GLMM, N = 283, X^2^_9_ = 31.508, p < 0.001), and comparisons between lit conditions revealed that flight-to-light increased with focal light intensity (GLMM, N = 255, X^2^_2_ = 24.286, p < 0.001; Figure 3F), but not focal light type , or spectrum (GLMM, N = 255, X^2^_2_ = 3.705, p =0.157). Moths exposed to lights at 30 lx were more likely to fly to the light apparatus than those experiencing the lowest light intensity of 0.3 lx, while other pairwise comparisons did not reach significance (Tukey’s, p_30 lx–0.3lx_ < 0.001, p_30 lx–3 lx_ = 0.0898, p_3 lx–0.3 lx_ = 0.0521). In addition, light-trapped moths were over twice as likely to end their flight at the light than net-caught individuals (GLMM, X^2^_1_ = 6.945, p = 0.00841; Estimate for light traps vs. nets = 0.851; OR = 2.342; Figure 3G). Conversely, the probability of moths flying upwards was also independent of light type but decreased with higher focal light intensities, though the effect of capture method did not reach significance (Supplementary Table S3).

In experiment 2, flight-to-light behaviour in repeated treatments was more frequent in 2025 than 2024, so data from each year were analysed separately (GLMM, N = 115, Year: X^2^_1_ = 14.091, p < 0.001; see methods). In both cases, background light levels were key to determining flight outcomes, but increasing background lighting had different effects. In 2024, flight-to-light was twice as likely to be observed under the medium background lighting condition than under low light, with no effect of focal light structure (single or multiple lights; GLMM, N = 152, Light structure * Background light level: X^2^_1_ = 0.521, p = 0.471; Light structure: X^2^_1_ = 0.587, p = 0.444; Background light level: X^2^_1_ = 6.209, p = 0.0127; Estimate for medium light level = 0.999, OR = 2.716, Figure 3H). In 2025, there was a significant interaction between background lighting and focal light structure (GLMM, N = 112, Light structure * Background light level: X^2^_1_ = 5.248, p = 0.0220, Figure 3J), but flight-to-light was always more likely under low background light levels than under high light (Tukey’s, Estimate for low compared to high background lighting, with a single focal light = 3.65, p < 0.001;

Estimate with multiple lights = 1.34, p = 0.0338). As in experiment 1, moths were far more likely to fly to light if they had been collected with light traps in 2024 (GLMM, Capture method: X^2^_1_ = 6.364, p = 0.0116, estimate for traps = 1.073, OR = 2.924, Figure 3I), but this effect was not observed in 2025 (GLMM, X^2^_1_ = 1.359, p = 0.244). However, main effects of background light level and capture method were supported by analysis of the probability of moths flying upwards: across both years, more moths flew upwards with increasing background light intensity, and light-trapped moths were half as likely to fly upwards than moths caught with nets (Supplementary Table S3). Trial number had no effect on flight-to-light behaviour in either year (GLMM, Trial number X^2^_1_ = 1.296, p = 0.295; X^2^_1_ = 0.367, p = 0.545 resp.), or on the probability of upwards flight (Supplementary Table S3).

### Flight tortuosity

Moths exposed to artificial lights are known to perform erratic flights [16] or spiral towards the light source [15]. Here, spirals were recorded by observers in 47% to 51% of trials with flights that met the analysis criteria, across experiments 1 and 2. Harmful erratic spiralling behaviour – defined as spiralling downwards, or performing three or more consecutive spirals in any direction (Figure 4A-B) – was seen in 17% and 22% of flights respectively. The probability of this erratic spiralling behaviour occurring was greater under the high focal light intensity in experiment 1, while light type, or spectrum, had no effect (GLMM, N = 255, Intensity * Spectrum: X^2^_4_ = 0.895, p = 0.925; Spectrum: X^2^_2_ = 0.605, p = 0.739; Intensity: X^2^_2_ = 25.387, p < 0.001; Tukey’s: p_30 lx–0.3lx_ = 0.00272, p_30 lx–3 lx_ = 0.00165, p_3 lx–0.3 lx_ = 0.958 ; Figure 4C). In experiment 2, this behaviour was more common when moths were exposed to a single rather than multiple lights, and under lower backgrounds light levels, although pairwise comparisons between background light levels did not reach significance (GLMM, N = 264, Light structure * Background light level: X^2^_2_ = 4.066, p = 0.131; Background light level: X^2^_2_ = 6.198, p = 0.0451; Light structure: X^2^_2_ = 5.737, p = 0.0166; Tukey’s: p_high-low_ = 0.0585, p_high-medium_ = 0.113, p_medium-low_ = 0.987; Figure 4E). Capture method and trial number had no effect (GLMMs; experiment 1, Capture method: X^2^_1_ = 2.117, p = 0.146; experiment 2: Capture method: X^2^_1_ = 1.278, p = 0.258; Trial number: X^2^_1_ = 0.129, p = 0.720). Similar trends were found for the probability of observing any spirals at all, confirming the trends for focal light intensity and structure, and background light levels (Supplementary Table S3).

**Figure 4:**
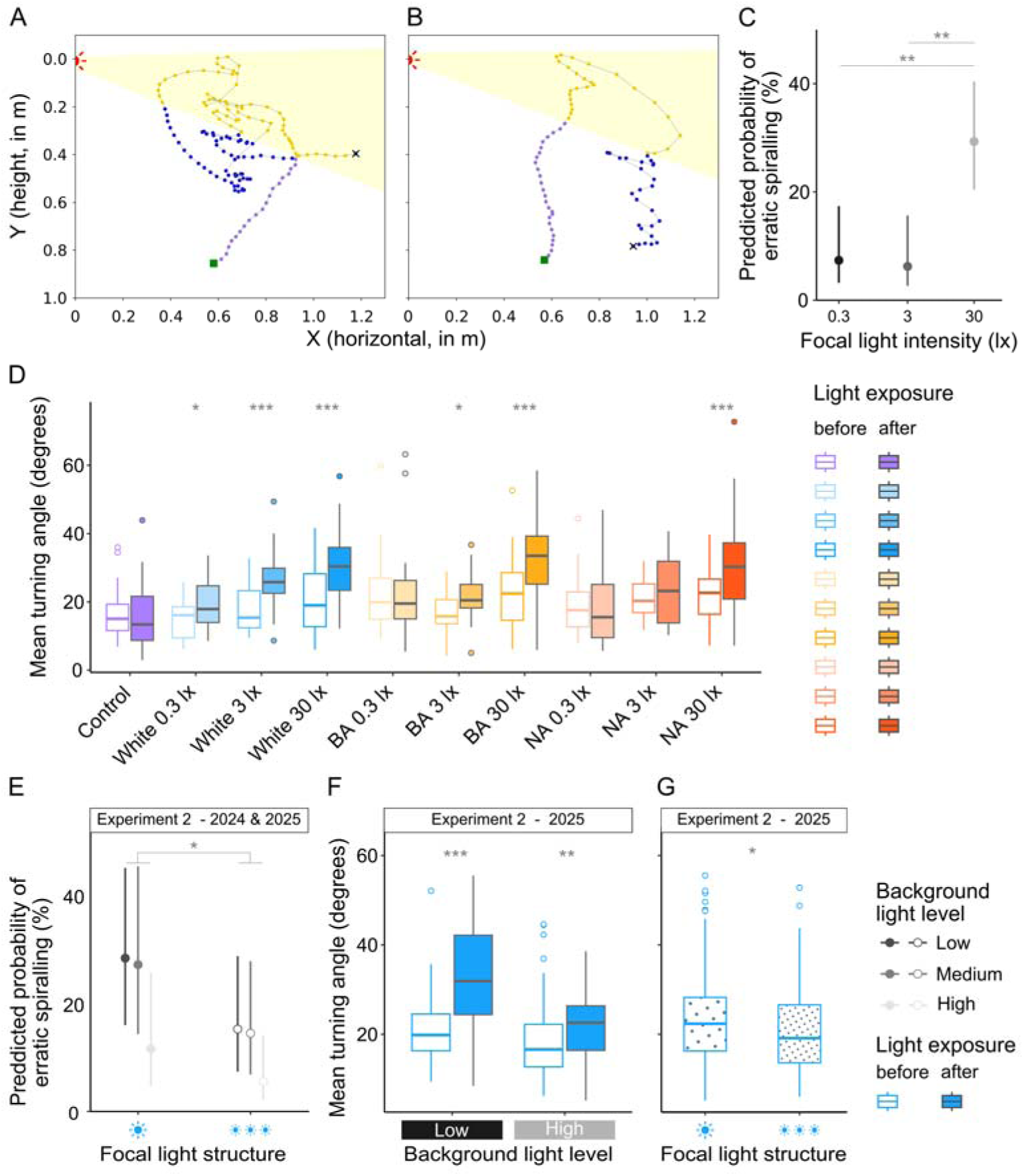
Flight tortuosity. (A-B) Orthogonal projections in XY (camera view) for example tracks in which moths displayed erratic spiralling behaviour, spiralling in place **(A)** or down to the ground **(B)**. Shading indicates approximate position of the light beam. **(C)** Predicted probability of erratic spiralling behaviour in experiment 1, according to the best statistical model. **(D)** Mean turning angles before and after exposure to the focal light beam, for every treatment in experiment 1. **(E)** Predicted probability of erratic spiralling behaviour in experiment 2, according to the best statistical model. **(F-G)** Mean turning angles before and after exposure to the light beam in experiment 2, per background light level **(F)** and focal light structure **(G)**. In **C** & **E**, dots represent predicted probabilities for fixed effects only, and whiskers 95% confidence intervals. In **D, F** & **G**, boxplots represent the median and interquartile ranges, and whiskers 95% confidence intervals. Asterisks show significance levels: * = p < 0.05, ** = p < 0.01, ***: p < 0.001.

To investigate how artificial light properties affect flight paths more quantitatively, we measured the tortuosity of tracked flights as the mean and standard deviation of turning angles in three dimensions. Calculated across the whole flight paths, both these metrics indicated greater tortuosity under the highest focal light intensity in experiment 1, with no significant differences between light spectra or capture methods, matching results for spiralling behaviour (Supplementary Table S4). Yet splitting flight paths into segments before and after exposure to the focal light showed that light spectrum did modulate the impact of light exposure on flight tortuosity, in line with expectations from moth photoreceptor sensitivities. Mean turning angle significantly increased once moths had crossed into the light beam under all intensities of white light, but only under the two higher intensities for broadband amber light, and the highest light intensity of 30 lx for the narrowband amber LED (linear mixed effects model (LMM), N = 510, Focal light treatment * Light exposure: X^2^_9_ = 38.123, p < 0.001; Capture method: X^2^_1_ = 0.715, p = 0.398; Figure 4D). Changes in the standard deviation of turning angle after exposure to the light beam also depended on light treatment, with significant increases only associated with white LEDs and the highest level of broadband amber LEDs (LMM, N = 510, Focal light treatment * Light exposure: X^2^_9_ = 19.918, p = 0.0184; Capture method: X^2^_1_ = 1.554, p = 0.213; Tukey’s pairwise contrasts for effect of light exposure for each treatment: Estimate_White 30 lx_ = 0.414, p_White 30 lx_ = 0.0334; Estimate_White 3 lx_ = 0.968, p_White 3 lx_ < 0.001; Estimate_White 0.3 lx_ = 0.713, p_White 0.3 lx_ = 0.0238; Estimate_Broadband amber 30 lx_ = 0.539, p_Broadband amber 30 lx_ = 0.0058; all others p > 0.05). There was no difference in either metric between equivalent flight segments in the control treatment.

In experiment 2, measures of flight tortuosity consistently indicated straighter paths under the highest background light level, and to a lesser extent when moths were exposed to a single rather than multiple lights. Mean turning angles across whole flights were higher overall in 2025 than 2024 (LMM, N = 115, Year: X^2^_1_ = 7.416, p = 0.00647), so results were analysed separately. There were no significant effects of any treatment variables in 2024, when only low and medium background light levels were tested, even when analysing flight paths split into segments before and after exposure to the focal light beam (Supplementary Tables S4, S5). In 2025, the mean turning angles across whole

flights were significantly smaller under high than low background lighting, but effects of light structure were not significant (LMM, N = 112, Background light level: X^2^_1_ = 8.875, p = 0.00289, Supplementary Table S4). Differences between focal light structures became discernible when comparing flight segments before and after exposure to the focal lights: crossing into the light beam always led to an increase in mean turning angles, but this effect was smaller under high background light levels, and mean turning angles were smaller overall in trials with multiple focal lights (LMM, N = 218, Background light level* Light exposure: X^2^_1_ = 11.232, p < 0.001; Light structure: X^2^_1_ = 5.892, p = 0.0152; Tukey’s for effect of exposure under high and low background lighting respectively: Estimate = 3.65, p = 0.0077 and Estimate = 11.22, p < 0.001). Results for the standard deviation of turning angle, analysed across both years, matched those for mean turning angle in 2025, suggesting that turning angles were less variable under higher background lighting and in trials with multiple focal lights (Supplementary Tables S4, S5).

## Discussion

With these experiments, we establish a simple set-up to track moth flight paths in 3D, and show that multiple characteristics of LED lights contribute to disrupting flight behaviour in a wide range of nocturnal moth species. The intensity of a light source and its contrast to background light conditions were particularly important. Within a range of realistic intensities, exposure to higher intensity lights was associated with a higher probability of flight-to-light and more tortuous flights, irrespective of light spectrum. By contrast, higher background light levels reduced these harmful impacts: under the highest level of background light tested, just one order of magnitude lower than the focal 30 lx light, moths were less likely to be attracted to the light, and flight tortuosity increased less upon exposure to the light beam, than when this same focal light was tested in dark conditions similar to natural nighttime light levels. Yet these high background light levels also dramatically reduced the probability of moths taking flight at all, adding to recent evidence of activity suppression under artificial lights [7,8]. This suggests that both bright lights in dark places and high ambient light levels impede moth movement, but in different ways. Meanwhile, exposure to a single bright light versus three separate lights combining to produce the same illuminance did not affect flight-to-light behaviour, but was associated with a higher probability of erratic spiralling, and more tortuous flights.

Notably, there was little impact of different light spectra on moth flight behaviour in our experiments, with no difference in the probability of flight-to-light between white and amber LEDs of equivalent illuminance. While moths and other nocturnal insects are generally more attracted to short wavelengths of light [17,24], evidence of differences in attraction between LED types is more equivocal [42–44,12], including between white and PC amber LEDs [45]. Tracking moth flight paths in 3D did reveal more subtle effects of light spectrum, suggesting that white LEDs may affect flight tortuosity at lower intensities (and hence greater distances) than broadband amber LEDs, and even more so compared to narrowband amber LEDs, as expected from moth photoreceptor sensitivities [46]. This pattern also matches the effects of these same light types on activity in nocturnal moths [8], and supports increasing evidence that amber lighting, while potentially less disruptive than white LEDs, will still have severe impacts on insect behaviour at realistic light intensities [47]. Other experiments with nocturnal hawkmoths have found conflicting results, with variation in flight behaviour associated with differences between LED spectra rather than intensity [14], but all intensities used there were far higher (150 - 590 lx) than the ones tested in this study. As red-shifted light may also be detrimental to other species [25], prioritising mitigation strategies that reduce the overall intensity and reach of artificial lights is likely to be most beneficial for biodiversity [21].

Importantly for future work on moth responses to artificial lights, we found significant differences between the behaviour of moths collected using light traps or by hand with nets. These findings support recent studies that highlighted differences between estimates of abundance and diversity based on surveys with light traps versus other methods [48,49], and variation in activity levels between net- and trap-caught individuals [8]. Samples from light traps may be affected by environmental conditions and competition from other light sources [50], or be biased towards species or individuals that are inherently more attracted to light, which is problematic for experiments subsequently testing responses to light. Light-trapped moths also bear the costs of a night spent in the trap with no access to food, and potential damage associated with crashing into a light, heat and dehydration. Catching insects on the wing at night, with butterfly nets, presents a less harmful alternative that potentially better represents the community of nocturnal moths active in the environment [51]. In our experiments, light-trapped moths were not only half as likely to take off than net-caught moths, but were also more likely to end their flights at the light; this suggests that light-trapped moths genuinely differed in their response to light, rather than simply being in worse condition. Similarly to other work collecting moths with both techniques, our light-trapped samples included more Noctuidae than the net-caught samples [51], but family had little impact on flight-to-light. In addition, comparisons between species caught using both methods suggest that sampling biases are unlikely to explain the behavioural differences we observed. While further work is needed to unpick the role of species composition, sex, age, condition, and individual variation in shaping moth behaviour towards lights, differences related to collection methods should be explicitly considered [50].

Our experiments also highlight new opportunities for studying how artificial lights affect flying insects. Tracking nocturnal moths in flight is a difficult task, due to their small size and the dim lighting conditions in which they are active, and relatively few studies have mapped flight paths in three dimensions. Notable exceptions include pioneering research on the predator-prey relationships of bats and moths [52] and some work on pheromone tracking [53,54] while 3D tracking has only recently been used to investigate moth behaviour around artificial lights [15]. Until then, most studies of moth flight paths around lights either have relied on qualitative scoring of behaviour [16,55]or were restricted to larger moths, particularly hawkmoths (Sphingidae) and *Noctua pronuba*, to facilitate tagging [18] or detection by cameras [19], even when tracking only in two dimensions [18]. Yet technological advances, particularly in machine learning, have stimulated work on 3D tracking of unmarked insects in free flight [56], in low light and with species as small as mosquitoes [57] and micromoths [53]. Fabian et al. [15] used multiple approaches to provide compelling evidence that a simple dorsal orientation response, with moths keeping the light source at their backs, can explain many of the behaviours seen very close to lights. This involved recording 3D flight trajectories from both large moths carrying reflective markers in a laboratory setting, and from free-flying wild insects in the field, filmed with an infrared stereo camera system. Here, we tracked the flight paths of a wide range of unmarked moth species in three dimensions with an even simpler stereo set-up, without high-speed cameras, specific flight tunnel set-ups or proprietary software. By converting motion signals into colour patterns [58], we were able to enhance the visibility of small flying targets, allowing us to film moths from many different families, including species with wingspans below 2cm. While filming at higher spatial and temporal resolution with high-speed cameras reveals more detailed information about moth posture [15], recording their flight trajectories alone is enough to assess their responses to different light strategies. The advent of low-cost, accessible and flexible systems for tracking moths in 3D will greatly facilitate future research on insect responses to artificial lights.

Our results confirm that artificial lights do not only attract nocturnal insects, but also trigger a much wider range of behaviours [16,18]. Further work investigating moth flight paths could address outstanding gaps in understanding, including why some moths do not perform the typical dorsal response, and the mechanisms underpinning attraction to lights from greater distances [15]. The potential consequences of behaviours other than flight-to-light should also be explored, including recovery times for moths that fly down to the ground or freeze in place, and energetic costs of erratic flights. Disturbances to moth flight paths from lights might also interact with anti-predator behaviour; moths are known to respond differently to bat echolocation calls under LED lights [55], and the flight paths of moths exposed to lights might otherwise affect their vulnerability to predators. Combined with effects of lights on physiology and behaviours outside of flying and movement [4,12], it is clear that the attractiveness of lights is a useful but incomplete measure of the harms they may cause to nocturnal insects, let alone other species. Testing how adjusting different properties of lighting systems affects the various responses of moths to lights will be crucial to identifying the most effective mitigation strategies. Our results provide further support for prioritizing a reduction in light intensity over changes to light spectrum, and shielding lights to preserve dark refuges and reduce skyglow [15,29]. Other potential solutions would benefit from further experiments at a larger scale, including intermittent lights with timers or motion activators [59], and different configurations of lights; for example, properly assessing the benefits of deploying more numerous but dimmer lamps would require more tests, with field-realistic intensities and spacings of streetlights. Ultimately, a combination of methodological advances allowing detailed observations of moth responses to varying light properties, field trials with real streetlights [18,27,29] and simulations of lighting strategies at the landscape scale [60], will build a fuller picture of how nocturnal insects move around artificial lights, helping to guide practical recommendations for lighting policy.

## Acknowledgments

We thank Andrew Szopa-Comley for assistance with fieldwork and reviewing analysis code, and Helen Briolat, Maisy Inston, Charlie Rayner and William English for help with moth collection. AI-assisted technologies (ChatGPT, GPT-5.3 model) were used to help with the writing of Python scripts for video analyses.

## Author contributions

Conceptualization: JT, KJG, JB

Methodology: JT, ESB, JAMG, MVB

Investigation: ESB, JAMG, MVB

Formal Analysis: ESB, MVB

Visualization: ESB

Writing—original draft: ESB,

Writing—review & editing: JT, KJG, JB, ESB, JAMG, MVB

## References

1. Kyba CCM et al. 2017 Artificially lit surface of Earth at night increasing in radiance and extent. Science Advances 3, e1701528. (doi:10.1126/sciadv.1701528)

2. Kyba CCM, Altıntaş YÖ, Walker CE, Newhouse M. 2023 Citizen scientists report global rapid reductions in the visibility of stars from 2011 to 2022. Science 379, 265–268. (doi:10.1126/science.abq7781)

3. Gaston KJ, Ackermann S, Bennie J, Cox DTC, Phillips BB, Sánchez de Miguel A, Sanders D. 2021 Pervasiveness of biological impacts of artificial light at night. Integrative and Comparative Biology 61, 1098–1110. (doi:10.1093/icb/icab145)

4. Owens ACS, Cochard P, Durrant J, Farnworth B, Perkin EK, Seymoure B. 2020 Light pollution is a driver of insect declines. Biological Conservation 241, 108259. (doi:10.1016/j.biocon.2019.108259)

5. van Geffen KG, van Eck E, de Boer RA, van Grunsven RHA, Salis L, Berendse F, Veenendaal EM. 2015 Artificial light at night inhibits mating in a Geometrid moth. Insect Conservation and Diversity 8, 282–287. (doi:10.1111/icad.12116)

6. van Langevelde F, van Grunsven RHA, Veenendaal EM, Fijen TPM. 2017 Artificial night lighting inhibits feeding in moths. Biology Letters 13, 20160874. (doi:10.1098/rsbl.2016.0874)

7. Meah RJ et al. 2026 Light pollution creates multiple threats to the movement ecology of nocturnal arthropod taxa. Current Biology 36, 541–548.e4. (doi:10.1016/j.cub.2025.11.055)

8. Briolat ES, Galloway JAM, Cornelius E, Wright CJ, Bennie J, Gaston KJ, Troscianko J. 2026 Severe and widespread reductions in night-time activity of nocturnal moths under modern artificial lighting spectra. Proc Biol Sci 293, 20252704. (doi:10.1098/rspb.2025.2704)

9. Burt CS, Kelly JF, Trankina GE, Silva CL, Khalighifar A, Jenkins-Smith HC, Fox AS, Fristrup KM, Horton KG. 2023 The effects of light pollution on migratory animal behavior. Trends in Ecology & Evolution 38, 355–368. (doi:10.1016/j.tree.2022.12.006)

10. Boyes DH, Evans DM, Fox R, Parsons MS, Pocock MJO. 2021 Street lighting has detrimental impacts on local insect populations. Science Advances 7, eabi8322. (doi:10.1126/sciadv.abi8322)

11. van Grunsven RHA, van Deijk JR, Donners M, Berendse F, Visser ME, Veenendaal E, Spoelstra K. 2020 Experimental light at night has a negative long-term impact on macro-moth populations. Current Biology 30, R694–R695. (doi:10.1016/j.cub.2020.04.083)

12. Boyes DH, Evans DM, Fox R, Parsons MS, Pocock MJO. 2021 Is light pollution driving moth population declines? A review of causal mechanisms across the life cycle. Insect Conservation and Diversity 14, 167–187. (doi:10.1111/icad.12447)

13. Frank KD. 2006 Effects of artificial night lighting on moths. In Ecological Consequences of Artificial Night Lighting (eds C Rich, T Longcore), pp. 184–206. Washington, DC: Island Press.

14. Storms M, Degen T, Degen J. 2025 Female moths call in vain: Streetlights diminish the promise of mating. Ecological Entomology 50, 729–740. (doi:10.1111/een.13441)

15. Fabian ST, Sondhi Y, Allen PE, Theobald JC, Lin H-T. 2024 Why flying insects gather at artificial light. Nature Communications 15.

16. Fabusova M, Gaston KJ, Troscianko J. 2024 Pulsed artificial light at night alters moth flight behaviour. Biology Letters 20, 20240403. (doi:10.1098/rsbl.2024.0403)

17. Brehm G, Niermann J, Jaimes-Nino LM, Enseling D, Jüstel T, Axmacher J, Warrant E, Fiedler K. 2021 Moths are strongly attracted to ultraviolet and blue radiation. Insect Conservation and Diversity 14. (doi:10.1111/icad.12476)

18. Degen J et al. 2024 Shedding light with harmonic radar: Unveiling the hidden impacts of streetlights on moth flight behavior. Proceedings of the National Academy of Sciences 121, e2401215121. (doi:10.1073/pnas.2401215121)

19. Gaydecki P. 2019 Automated moth flight analysis in the vicinity of artificial light. Bulletin of Entomological Research 109, 127–140. (doi:10.1017/S0007485318000378)

20. Wagner DL. 2020 Insect declines in the Anthropocene. Annual Review of Entomology 65, 457–480. (doi:10.1146/annurev-ento-011019-025151)

21. Owens AC, Pocock MJ, Seymoure BM. 2024 Current evidence in support of insect-friendly lighting practices. Current Opinion in Insect Science 66, 101276. (doi:10.1016/j.cois.2024.101276)

22. Longcore T, Rodríguez A, Witherington B, Penniman JF, Herf L, Herf M. 2018 Rapid assessment of lamp spectrum to quantify ecological effects of light at night. Journal of Experimental Zoology Part A: Ecological and Integrative Physiology 329, 511–521. (doi:10.1002/jez.2184)

23. Gaston KJ, Sánchez de Miguel A. 2022 Environmental impacts of artificial light at night. Annual Review of Environment and Resources 47, 373–398. (doi:10.1146/annurev-environ-112420-014438)

24. Deichmann JL, Ampudia Gatty C, Andía Navarro JM, Alonso A, Linares-Palomino R, Longcore T. 2021 Reducing the blue spectrum of artificial light at night minimises insect attraction in a tropical lowland forest. Insect Conservation and Diversity 14, 247–259. (doi:10.1111/icad.12479)

25. Owens ACS, Dressler CT, Lewis SM. 2022 Costs and benefits of “insect friendly” artificial lights are taxon specific. Oecologia 199, 487–497. (doi:10.1007/s00442-022-05189-6)

26. Gaston KJ, Morrell S, Bennie J. 2025 Reddening the nighttime environment: Use of PC-amber LED lighting. Ecological Solutions and Evidence 6, e70144. (doi:10.1002/2688-8319.70144)

27. Bolliger J, Haller J, Wermelinger B, Blum S, Obrist MK. 2022 Contrasting effects of street light shapes and LED color temperatures on nocturnal insects and bats. Basic and Applied Ecology 64, 1–12. (doi:10.1016/j.baae.2022.07.002)

28. van Koppenhagen N, Haller J, Kappeler J, Gossner MM, Bolliger J. 2024 LED streetlight characteristics alter the functional composition of ground-dwelling invertebrates. Environmental Pollution 355, 124209. (doi:10.1016/j.envpol.2024.124209)

29. Dietenberger M, Jechow A, Kalinkat G, Schroer S, Saathoff B, Hölker F. 2024 Reducing the fatal attraction of nocturnal insects using tailored and shielded road lights. Commun Biol 7, 671. (doi:10.1038/s42003-024-06304-4)

30. Post CT, Goldsmith TH. 1965 Pigment migration and light-adaptation in the eye of the moth, galleria mellonella. The Biological Bulletin 128, 473–487. (doi:10.2307/1539906)

31. Berry S. 2022 The Use of Optical Coherence Tomography to Demonstrate Dark and Light Adaptation in a Live Moth. Environ Entomol 51, 643–648. (doi:10.1093/ee/nvac044)

32. Troscianko J. 2023 OSpRad: An open-source, low-cost, high-sensitivity spectroradiometer. Journal of Experimental Biology 226, jeb245416. (doi:10.1242/jeb.245416)

33. Troscianko J, O’Shea-Wheller TA, Galloway JAM, Gaston KJ. 2026 BehaveAI enables rapid detection and classification of objects and behavior from motion. PLOS Biology 24, e3003632. (doi:10.1371/journal.pbio.3003632)

34. Savitzky Abraham, Golay MJE. 1964 Smoothing and Differentiation of Data by Simplified Least Squares Procedures. Anal. Chem. 36, 1627–1639. (doi:10.1021/ac60214a047)

35. McLean DJ, Skowron Volponi MA. 2018 trajr: An R package for characterisation of animal trajectories. Ethology 124, 440–448. (doi:10.1111/eth.12739)

36. R Core Team. 2023 R: A language and environment for statistical computing.

37. Broooks ME, Kristensen K, van Benthem KJ, Magnusson A, Berg CW, Nielsen A, Skaug HJ, Maechler M, Bolker BM. 2017 glmmTMB Balances Speed and Flexibility Among Packages for Zero-inflated Generalized Linear Mixed Modeling. The R Journal 9, 378–400. (doi:10.32614/RJ-2017-066)

38. Akaike H. 1973 Information theory and an extension of the maximum likelihood principle. In (eds BN Petrov, F Csáki), pp. 267–281. Budapest, Hungary: Akadémia Kiadó.

39. Hartig F. 2024 DHARMa: Residual diagnostics for hierarchical (multi-Level / mixed) regression models. R package version 0.4.7.

40. Hothorn T, Bretz F, Westfall P. 2008 Simultaneous inference in general parametric models. Biometrical Journal 50, 346–363.

41. Lenth RV. 2025 emmeans: Estimated marginal means, aka least-squares means. R package version 1.10.6-090003.

42. Longcore T, Aldern HL, Eggers JF, Flores S, Franco L, Hirshfield-Yamanishi E, Petrinec LN, Yan WA, Barroso AM. 2015 Tuning the white light spectrum of light emitting diode lamps to reduce attraction of nocturnal arthropods. Phil. Trans. R. Soc. B 370, 20140125. (doi:10.1098/rstb.2014.0125)

43. Pawson SM, Bader MK-F. 2014 LED lighting increases the ecological impact of light pollution irrespective of color temperature. Ecological Applications 24, 1561–1568. (doi:10.1890/14-0468.1)

44. Wakefield A, Broyles M, Stone EL, Harris S, Jones G. 2018 Quantifying the attractiveness of broad-spectrum street lights to aerial nocturnal insects. Journal of Applied Ecology 55, 714–722. (doi:10.1111/1365-2664.13004)

45. Gaston KJ, Morrell S, Bennie J. 2025 Reddening the nighttime environment: Use of PC-amber LED lighting. Ecological Solutions and Evidence 6, e70144. (doi:10.1002/2688-8319.70144)

46. Longcore T. 2023 A compendium of photopigment peak sensitivities and visual spectral response curves of terrestrial wildlife to guide design of outdoor nighttime lighting. Basic and Applied Ecology 73, 40–50. (doi:10.1016/j.baae.2023.09.002)

47. Czarnecka M, Grubisic M, Pilotto F, Jechow A, Hölker F. 2025 Colours of the Night: Spectrum-Specific Impacts of Light Pollution on Biota. Global Change Biology 31, e70569. (doi:10.1111/gcb.70569)

48. Battles I, Burkness E, Crossley MS, Edwards CB, Holmstrom K, Hutchison W, Ingerson-Mahar J, Owens D, Owens ACS. 2024 Moths are less attracted to light traps than they used to be. J Insect Conserv 28, 1007–1018. (doi:10.1007/s10841-024-00588-x)

49. Grenis K, Nufio C, Wimp GM, Murphy SM. 2023 Does artificial light at night alter moth community composition? Philosophical Transactions of the Royal Society B: Biological Sciences 378, 20220365. (doi:10.1098/rstb.2022.0365)

50. Chan W-P et al. 2025 Historical behavioural data disentangle evolutionary and environmental drivers of recent declines in insect attraction to light. Proc Biol Sci 292, 20252319. (doi:10.1098/rspb.2025.2319)

51. Macgregor CJ, Evans DM, Fox R, Pocock MJO. 2017 The dark side of street lighting: Impacts on moths and evidence for the disruption of nocturnal pollen transport. Global Change Biology 23, 697–707. (doi:10.1111/gcb.13371)

52. Corcoran AJ, Conner WE. 2016 How moths escape bats: predicting outcomes of predator–prey interactions. J Exp Biol 219, 2704–2715. (doi:10.1242/jeb.137638)

53. El-Sayed AM, Gödde J, Arn H. 2000 A Computer-Controlled Video System for Real-Time Recording of Insect Flight in Three Dimensions. Journal of Insect Behavior 13, 881–900. (doi:10.1023/A:1007866602219)

54. Rutkowski AJ, Quinn RD, Willis MA. 2009 Three-dimensional characterization of the wind-borne pheromone tracking behavior of male hawkmoths, Manduca sexta. J Comp Physiol A 195, 39–54. (doi:10.1007/s00359-008-0380-9)

55. Wakefield A, Stone EL, Jones G, Harris S. 2015 Light-emitting diode street lights reduce last-ditch evasive manoeuvres by moths to bat echolocation calls. R Soc Open Sci. 2, 150291. (doi:10.1098/rsos.150291)

56. Sun C, Gaydecki P. 2021 A Visual Tracking System for Honey Bee (Hymenoptera: Apidae) 3D Flight Trajectory Reconstruction and Analysis. J Insect Sci 21. (doi:10.1093/jisesa/ieab023)

57. Spitzen J, Spoor CW, Kranenbarg S, Beeuwkes J, Grieco F, Noldus LPJJ, van leeuwen JL, Takken W. 2008 Track3D: Visualization and flight track analysis of Anopheles gambiae s.s. mosquitoes. Proceedings of Measuring Behavior 2008 - 6th International Conference on Methods and Techniques in Behavioral Research

58. Troscianko J, O’Shea-Wheller TA, Galloway J, Gaston KJ. 2025 BehaveAI: a framework for rapidly detecting and classifying objects and behaviour from motion. , 2025.11.04.686536. (doi:10.1101/2025.11.04.686536)

59. Owens AC, Pocock MJ, Seymoure BM. 2024 Current evidence in support of insect-friendly lighting practices. Current Opinion in Insect Science 66, 101276. (doi:10.1016/j.cois.2024.101276)

60. Morrell S, Hatchell J, Wordingham F, Bennie J, Inston MJ, Shannon JP, Rayner CW, Gaston KJ. 2026 Mitigating the environmental and ecological impacts of evolving city-scale streetlighting installations. J R Soc Interface 23, 20250453. (doi:10.1098/rsif.2025.0453)

