## Supplementary material for "Effects of artificial light colour, intensity, structure and contrast on moth flight behaviour"

### <u>Supplementary Materials for:</u> Effects of artificial light colour, intensity, structure and contrast on moth flight behaviour

##### Supplementary Figures S1 – S3; Supplementary Tables S1 – S5

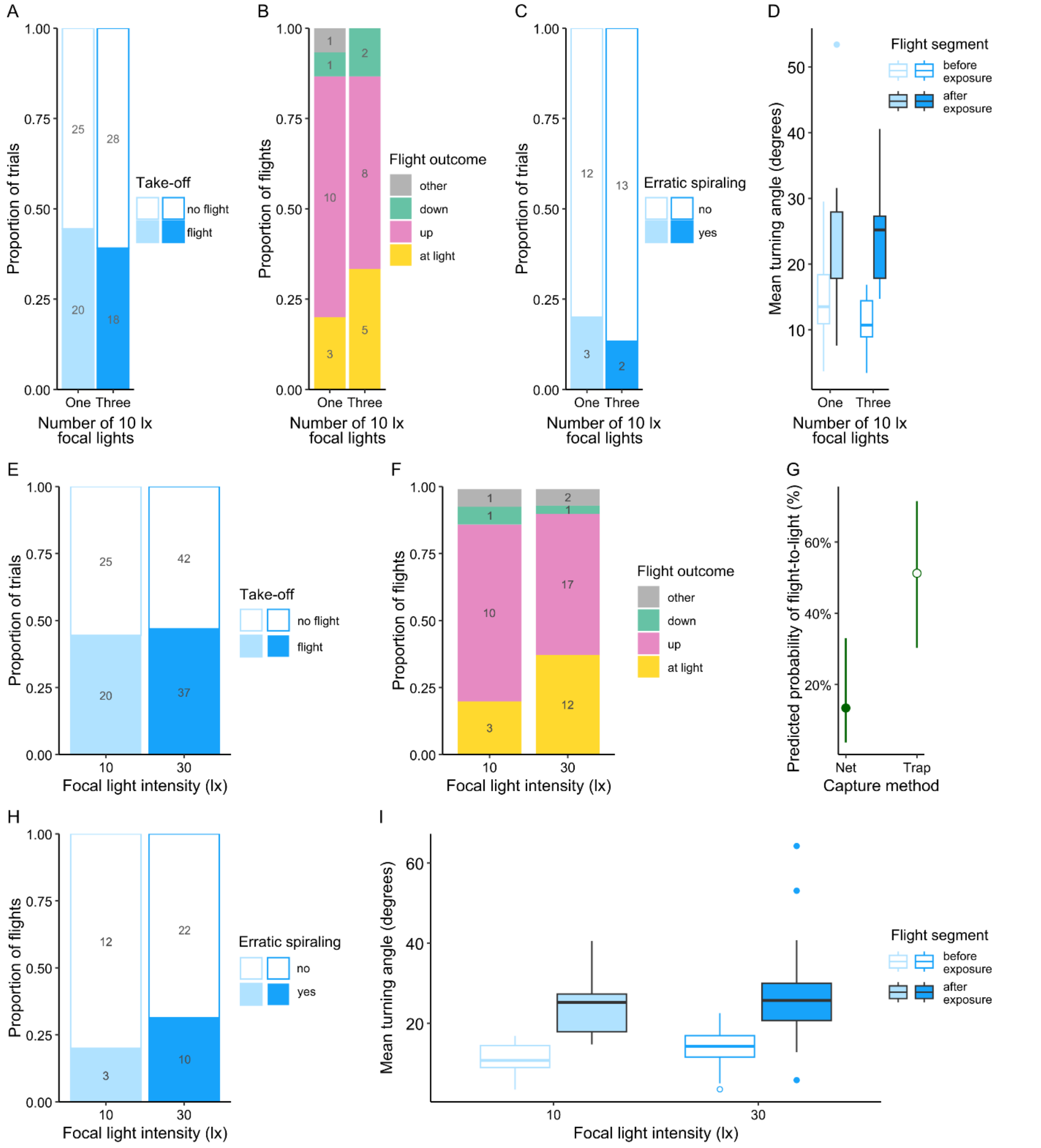


<u>Supplementary Figure S1</u>. **Effects of exposure to a single 10 lx focal light.** There was no difference in any aspect of moth behaviour between trials exposed to a single 10 lx and trials with either multiple 10 lx lights **(A-D)** or a single 30 lx light **(E-I)**. **(A)** Proportion of trials in which moths took off between single and multiple 10 lx treatments. **(B)** Proportion of trials with different outcomes. **(C)** Proportion of trials with erratic spirals. **(D)** Mean turning angle before and after exposure to the focal light beam. **(E)** Proportion of trials in which moths took off between 10 lx and 30 lx treatments. **(F)** Proportion of trials with different outcomes. **(G)** Predicted probability of flight-to-light in trials with 10 or 30 lx focal lights, based on capture method. Trap-caught moths were more

likely to fly to light than net-caught moths (GLMM, N = 47, capture method: $X^2_1 = 6.924$, p = 0.00850); dots represent predicted probabilities for fixed effects only, and whiskers 95% confidence intervals. **(H)** Proportion of trials with erratic spirals, when exposed to 10 or 30 lx focal lights. **(I)** Mean turning angle before and after exposure to the focal light beam, with 10 or 30 lx lights. In **D** & **I**, boxplots represent the median and interquartile ranges, and whiskers 95% confidence intervals.

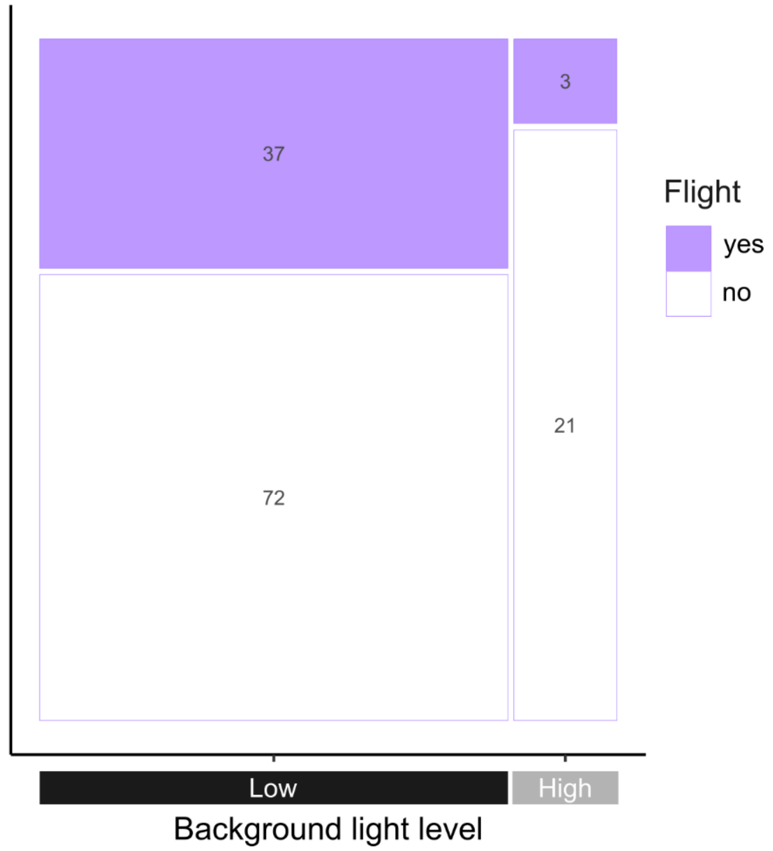


**Supplementary Figure S2. Frequency of successful flights for typical control conditions in experiment 1 in 2025, with low background lighting levels (N = 109), compared to additional trials testing control conditions with high background light levels (matching the conditions in experiment 2, approximately 3lx, N = 24).** A binomial mixed effects model found a significantly higher probability of moths taking off under low than high background light levels (GLMM, N = 133, $X^2_1 = 5.222$, p = 0.0223; estimate for high light level = -1.354, odds ratio = 0.258). In the mosaic plot, column width and numbers in grey represent the number of trials in each group.

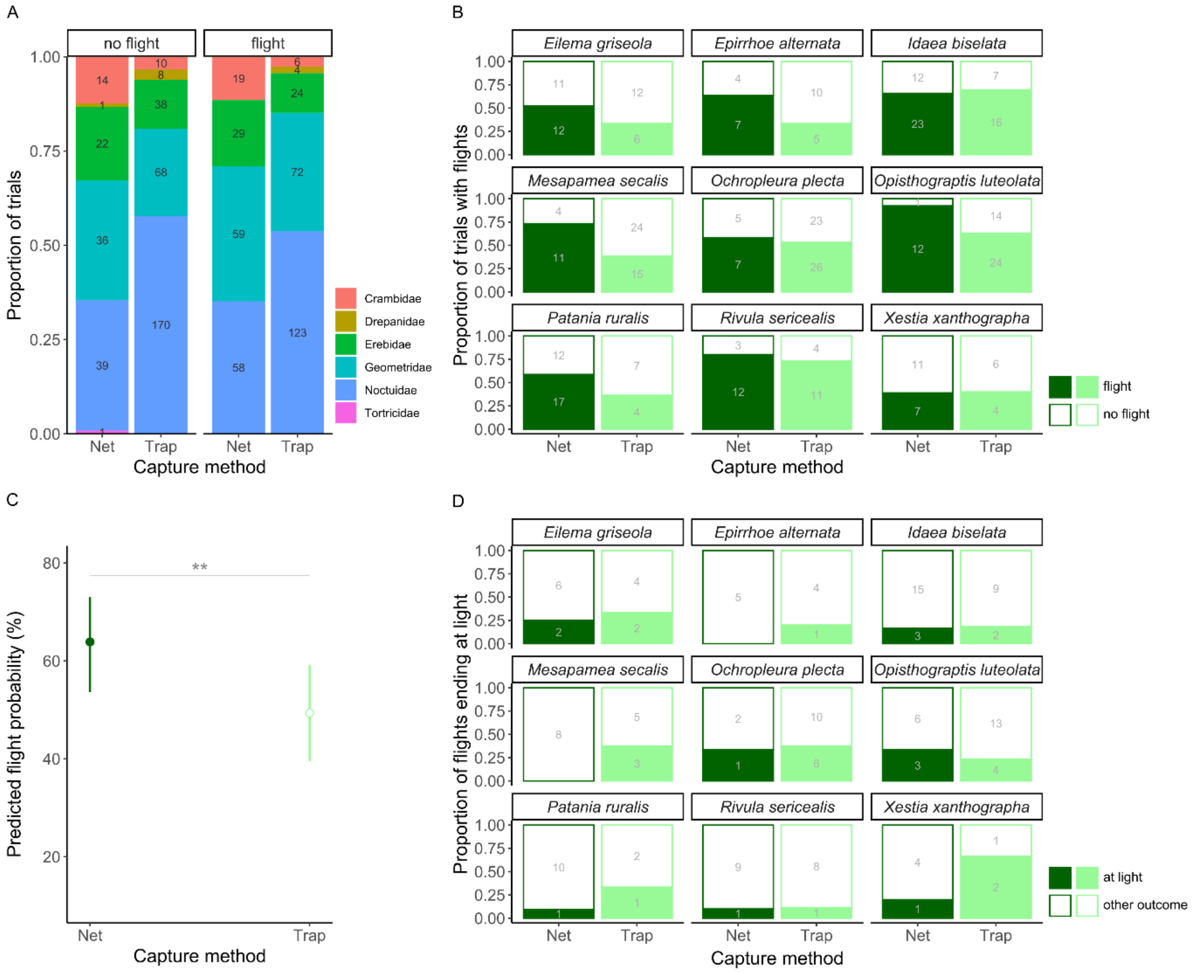


**Supplementary Figure S3. Effects of species and capture method in experiment 1.**
**(A)** Proportions of moth families tested in experiment 1; these were similar between capture methods, and between trials in which moths took off or not. **(B)** Proportion of trials in which moths took off, for nine species with at least 10 individuals collected using both methods. Within species, moths were either more likely to fly if caught with nets than with traps, or there was no difference. **(C)** Predicted flight probability for moths of those nine species, based on capture method. A binomial linear mixed effects model with species and family-level random effects found a significant effect of capture method, with a similar effect size as for the main dataset (GLMM, N = 389, $X^2_1 = 7.137$, p = 0.00755, estimate for traps = -0.597, odds ratio = 0.550). Focal light treatment was not included in the model, as it had no effect on flight probability in the main dataset. Dots represent predicted probabilities for fixed effects only, based on this model, and whiskers 95% confidence intervals. **(D)** Proportion of flights ending at the light apparatus, for the same nine species. Differences between capture method follow the same trend as for flight probability **(B)**, but the effect of capture method was not significant (binomial linear mixed effects model with species and family-level random effects, capture method and focal light intensity as fixed effects; N = 141, capture method: $X^2_1 = 2.955$, p = 0.0856).

**Supplementary Table S1.** **Moth species, with the number of individuals used per species for each experiment.**

| Family | Species | Experiment 1 | Experiment 1 supplementary trials | Experiment 2 | Total |
|---|---|---|---|---|---|
| **Crambidae** | *Anania hortulata* | 9 | - | - | 9 |
| | *Patania ruralis* | 41 | - | 22 | 63 |
| **Drepanidae** | *Thyatira batis* | 13 | - | 5 | 18 |
| **Erebidae** | *Eilema depressa* | 1 | - | - | 1 |
| | *Eilema griseola* | 41 | - | - | 41 |
| | *Euplagia quadripunctaria* | 7 | - | - | 7 |
| | *Herminia grisealis* | 2 | - | - | 2 |
| | *Hypena obsitalis* | - | - | 1 | 1 |
| | *Hypena proboscidalis* | 19 | 1 | 64 | 84 |
| | *Miltochrista miniata* | 7 | - | - | 7 |
| | *Phragmatobia fuliginosa* | 3 | - | - | 3 |
| | *Scoliopteryx libatrix* | 4 | - | 1 | 5 |
| **Geometridae** | *Abraxas grossulariata* | 7 | - | - | 7 |
| | *Cabera pusaria* | 5 | - | - | 5 |
| | *Camptogramma bilineata* | 5 | 1 | 2 | 8 |
| | *Chloroclysta siterata* | 1 | - | - | 1 |
| | *Colostygia pectinataria* | - | - | 9 | 9 |
| | *Cosmorhoe ocellata* | - | - | 1 | 1 |
| | *Crocallis elinguaria* | 1 | - | - | 1 |
| | *Dysstroma truncata* | 9 | - | 15 | 24 |
| | *Ecliptopera silaceata* | 34 | - | 5 | 39 |
| | *Epione repandaria* | 1 | - | - | 1 |
| | *Epirrhoe alternata* | 26 | - | 14 | 40 |
| | *Hemithea aestivaria* | 1 | - | - | 1 |
| | *Hydriomena furcata* | 2 | - | - | 2 |
| | *Idaea aversata* | 3 | - | - | 3 |
| | *Idaea biselata* | 60 | - | 14 | 74 |
| | *Opisthograptis luteolata* | 51 | 2 | 37 | 90 |
| | *Peribatodes rhomboidaria* | 8 | - | 2 | 10 |
| | *Scopula imitaria* | - | - | 1 | 1 |
| | *Scopula marginepunctata* | 2 | - | - | 2 |
| | *Selenia dentaria* | 7 | - | - | 7 |
| | *Xanthorhoe designata* | 2 | - | 4 | 6 |
| | *Xanthorhoe fluctuata* | 12 | - | 4 | 16 |
| **Noctuidae** | *Abrostola sp.* | 23 | - | 4 | 27 |
| | *Acronicta rumicis* | 7 | - | - | 7 |
| | *Agrotis ipsilon* | 3 | - | 5 | 8 |

| Family | Species | Experiment 1 | Experiment 1 supplementary trials | Experiment 2 | Total |
|---|---|---|---|---|---|
| **Noctuidae** | *Agrotis segetum* | - | 1 | 1 | 2 |
| | *Aporophyla nigra* | 1 | - | - | 1 |
| | *Autographa gamma* | 28 | - | 17 | 45 |
| | *Caradrina clavipalpis* | 1 | - | - | 1 |
| | *Cosmia trapezina* | 3 | - | - | 3 |
| | *Craniophora ligustri* | 2 | - | - | 2 |
| | *Helicoverpa armigera* | - | - | 1 | 1 |
| | *Hoplodrina sp.* | 2 | - | - | 2 |
| | *Luperina testacea* | 2 | - | 1 | 3 |
| | *Mesapamea secalis* | 54 | - | 8 | 62 |
| | *Mesoligia furuncula* | 1 | - | - | 1 |
| | *Mythimna unipuncta* | - | - | 1 | 1 |
| | *Noctua comes* | 1 | - | - | 1 |
| | *Noctua janthe* | 25 | - | 3 | 28 |
| | *Noctua pronuba* | 100 | 11 | 69 | 180 |
| | *Ochropleura plecta* | 61 | - | 16 | 77 |
| | *Phlogophora meticulosa* | 20 | 5 | 57 | 82 |
| | *Polymixis xanthomista* | 2 | - | - | 2 |
| | *Rivula sericealis* | 31 | - | 7 | 38 |
| | *Tiliacea citrago* | 1 | - | - | 1 |
| | *Xanthia togata* | 3 | - | - | 3 |
| | *Xestia c-nigrum* | 21 | - | 29 | 50 |
| | *Xestia triangulum* | 1 | - | - | 1 |
| | *Xestia xanthographa* | 28 | 3 | 42 | 73 |
| **Tortricidae** | *Agapeta hamana* | 1 | - | - | 1 |
| **TOTAL** | | 806 | 24 | 462 | 1292 |

**Supplementary Table S2. Sample sizes for experiments 1 and 2, per year and treatment group.** In experiment 1, each trial represents a unique individual moth. In experiment 2, numbers in the table refer to trials, not individual moths; each moth takes part in two trials, with different focal light structures but the same background light level. Numbers for the single and multiple (three) focal light treatments do not match for the medium and high background light level, due to additional test groups outside the main experiment: 47 moths from the medium background light group were used for trials with a single 10 lx focal light rather than a single 30 lx focal light, and 24 moths from the high background light group were used for control trials with a high background light but no focal light (see Supplementary Figure S1). Of the 806 trials in experiment 1 and 853 trials in experiment 2, respectively 5 and 19 were excluded due to experimenter error or damage to the moth, and moths took flight within one minute in 394 and 295 trials respectively. Further reductions in the number of flight paths are due to exclusion of flight paths with fewer than five points, or where moths did not cross the light beam.

| | Treatment | Trials | | | Flight paths for analysis | | | |
|---|---|---|---|---|---|---|---|---|
| | | **2024** | **2025** | **Total** | **2024 – 2D** | **2024 – 3D** | **2025 – all 3D** | **Total** |
| **Experiment 1** | Control | 85 | 109 | 194 | 16 | 7 | 24 | 47 |
| | White 30 lx | 86 | 41 | 127 | 12 | 30 | 15 | 57 |
| | White 3 lx | - | 45 | 45 | - | - | 21 | 21 |
| | White 0.3 lx | - | 55 | 55 | - | - | 18 | 18 |
| | Broadband amber 30 lx | 92 | 33 | 125 | 4 | 30 | 14 | 48 |
| | Broadband amber 3 lx | - | 47 | 47 | - | - | 20 | 20 |
| | Broadband amber 0.3 lx | - | 50 | 50 | - | - | 19 | 19 |
| | Narrowband amber 30lx | 53 | 35 | 88 | - | 18 | 15 | 33 |
| | Narrowband amber 3 lx | - | 40 | 40 | - | - | 19 | 19 |
| | Narrowband amber 0.3 lx | - | 35 | 35 | - | - | 20 | 20 |
| | | 316 | 490 | **806** | 32 | 85 | 185 | **302** |
| **Experiment 2** | Low background light – single focal light | 122 | 46 | 168 | - | 35 | 22 | 57 |
| | Low background light – multiple focal lights | 122 | 46 | 168 | - | 39 | 19 | 58 |
| | Medium background light – single focal light | 83 | - | 83 | - | 32 | - | 32 |
| | Medium background light – multiple focal lights | 130 | - | 130 | - | 46 | - | 46 |
| | High background light – single focal light | - | 164 | 164 | - | - | 33 | 33 |
| | High background light – multiple focal lights | - | 140 | 140 | - | - | 38 | 38 |
| | | 457 | 396 | **853** | | 152 | 112 | **264** |

**<u>Supplementary Table S3.</u> Outputs for models of the probability of upwards flight and spiralling behaviour (any spiral observed), related to Figures 3 & 4.** Estimates (Est.) and standard error (SE) not provided for factors with more than 2 levels. Factor reference levels are trial one, net collection, single focal light and low background light level for trial number, capture method, focal light structure and background light level respectively. Tukey's post-hoc tests were run when significant effects were found for factors with more than two levels. Significant effects highlighted in italics.

| | Model | Factor | Estimate | SE | $X^2$ | df | p | Tukey's post-hoc tests |
|---|---|---|---|---|---|---|---|---|
| **Experiment 1** | Upwards flight | Spectrum * Intensity | - | - | 2.221 | 4 | 0.695 | - |
| | | Spectrum | - | - | 1.329 | 2 | 0.515 | - |
| | | Capture method (trap) | -0.496 | 0.283 | 3.110 | 1 | 0.0778 | - |
| | | *Intensity* | - | - | *46.102* | *2* | *< 0.001* | *30 lx – 0.3 lx:*<br>*Est. = -2.582, p < 0.001*<br>30 lx – 3 lx:<br>Est. = -0.643, p = 0.0955<br>*3 lx – 0.3 lx:*<br>*Est. = -1.939, p < 0.001* |
| | Spiral in flight | Spectrum * Intensity | - | - | 4.700 | 4 | 0.320 | - |
| | | Capture method (trap) | -0.105 | 0.299 | 0.123 | 1 | 0.726 | - |
| | | Spectrum | - | - | 1.523 | 2 | 0.467 | - |
| | | *Intensity* | - | - | *32.349* | *2* | *< 0.001* | *30 lx – 0.3 lx:*<br>*Est. = 1.913, p < 0.001*<br>*30 lx – 3 lx:*<br>*Est. = 1.065, p = 0.00494*<br>3 lx – 0.3 lx:<br>Est. = 0.849, p = 0.111 |
| **Experiment 2** | Upwards flight | Focal light structure * Background light level | - | - | 4.151 | 2 | 0.126 | - |
| | | Trial number (two) | -0.265 | 0.297 | 0.813 | 1 | 0.367 | - |
| | | Light structure (multiple) | 0.222 | 0.291 | 0.587 | 1 | 0.443 | - |
| | | *Capture method (trap)* | *-0.711* | *0.328* | *5.016* | *1* | *0.0251* | - |
| | | *Background light level* | - | - | *23.593* | *2* | *< 0.001* | *High – Low:*<br>*Est. = 1.803, p < 0.001*<br>*High – Medium:*<br>*Est. = 1.359, p = 0.00488*<br>Medium – Low:<br>Est. = 0.443, p = 0.404 |

| | Model | Factor | Estimate | SE | $X^2$ | df | p | Tukey's post-hoc tests |
|---|---|---|---|---|---|---|---|---|
| **Experiment 2** | Spiral in flight, 2024 | Focal light structure * Background light level (multiple, medium) | -0.240 | 0.770 | 0.0968 | 1 | 0.756 | - |
| | | Capture method (trap) | 0.451 | 0.420 | 1.183 | 1 | 0.277 | - |
| | | Trial number (two) | -0.374 | 0.359 | 1.093 | 1 | 0.296 | - |
| | | Background light level (medium) | -0.0427 | 0.370 | 0.0133 | 1 | 0.908 | - |
| | | *Light structure (multiple)* | *-0.904* | *0.362* | *6.535* | *1* | *0.0106* | - |
| | Spiral in flight, 2025 | Focal light structure * Background light level (multiple, medium) | -1.256 | 1.005 | 1.624 | 1 | 0.203 | - |
| | | Capture method (trap) | -0.483 | 0.520 | 0.889 | 1 | 0.346 | - |
| | | Trial number (two) | -0.678 | 0.464 | 2.226 | 1 | 0.136 | - |
| | | Light structure (multiple) | -0.244 | 0.493 | 0.326 | 1 | 0.568 | - |
| | | *Background light level (high)* | *-1.502* | *0.491* | *10.032* | *1* | *0.00154* | - |

**<u>Supplementary Table S4.</u> Model outputs for analyses of flight tortuosity, calculated across whole flight paths.** For experiment 1, mean turning angle was square-root transformed to meet model assumptions, and both moth species and family were included as random effects. For experiment 2, mean turning angle is analysed separately for 2024 and 2025. For 2024, mean turning angle is not transformed, and only moth species and family, but not individual ID, are included as random effects; for 2025, mean turning angle is log transformed, but all random effects are included. Standard deviation of turning angle in experiment 2 is analysed across both years, with only moth species and family as random effects. Estimates (Est.) and standard error (SE) not provided for factors with more than 2 levels. Factor reference levels are trial one, net collection, single focal light and low background light level for trial number, capture method, focal light structure and background light level respectively. Significant effects are highlighted in italics. BA= Broadband amber, NA = Narrowband amber.

| | Model | Factor | Estimate | SE | $X^2$ | df | p | Tukey's post-hoc tests |
|---|---|---|---|---|---|---|---|---|
| **Experiment 1** | Mean turning angle, effect of treatment | Treatment * Capture method | - | - | 7.096 | 9 | 0.627 | - |
| | | Capture method (trap) | 0.107 | 0.120 | 0.801 | 1 | 0.371 | - |
| | | *Treatment* | - | - | *47.446* | *9* | *< 0.001* | *White 30 lx – Control: Est. = 0.861, p < 0.01*<br>*BA 30 lx – Control: Est. = 0.943, p < 0.01*<br>*NA 30 lx – Control: Est. = 0.828, p = 0.0111*<br>All other comparisons with control: p > 0.05 |
| | Mean turning angle, effect of focal light spectrum & intensity | Spectrum * Intensity | - | - | 8.900 | 4 | 0.0637 | - |
| | | Spectrum | - | - | 0.684 | 2 | 0.710 | - |
| | | Capture method (trap) | 0.106 | 0.121 | 0.731 | 1 | 0.393 | - |
| | | *Intensity* | - | - | *26.832* | *2* | *< 0.001* | *30 lx – 0.3 lx: Est. = 0.712, p < 0.001*<br>*30 lx – 3 lx: Est. = 0.398, p = 0.010*<br>3 lx – 0.3 lx: Est. = 0.314, p = 0.122 |
| | SD of turning angle, testing effect of treatment | Treatment * Capture method | - | - | 6.307 | 9 | 0.709 | - |
| | | Capture method (trap) | 1.752 | 1.161 | 2.302 | 1 | 0.129 | - |
| | | *Treatment* | - | - | *39.466* | *9* | *< 0.001* | *White 30 lx – Control: Est. = 8.123, p < 0.01*<br>*BA 30 lx – Control: Est. = 8.057, p < 0.01*<br>*NA 30 lx – Control: Est. = 7.619, p = 0.0247*<br>All other comparisons with control: p > 0.05 |

| | Model | Factor | Estimate | SE | $X^2$ | df | p | Tukey's post-hoc tests |
|---|---|---|---|---|---|---|---|---|
| **Experiment 1** | SD of turning angle, testing effect of focal light spectrum & intensity | Spectrum | - | - | 0.446 | 2 | 0.800 | - |
| | | Capture method (trap) | 1.515 | 1.186 | 1.573 | 1 | 0.210 | - |
| | | *Intensity* | *-* | *-* | *21.278* | *2* | *< 0.001* | *30 lx – 0.3 lx: Est. = 6.238, p < 0.001*<br>*30 lx – 3 lx: Est. = 4.022, p = 0.0106*<br>*3 lx – 0.3 lx: Est. = 2.216, p = 0.360* |
| **Experiment 2** | Mean of turning angle, in 2024 | Light structure * Background light level (multiple, medium) | 0.920 | 2.516 | 0.169 | 1 | 0.681 | - |
| | | Trial number (two) | -0.402 | 1.431 | 0.111 | 1 | 0.740 | - |
| | | Background light level (medium) | -0.801 | 1.223 | 0.406 | 1 | 0.524 | - |
| | | Capture method (trap) | 2.309 | 1.413 | 2.704 | 1 | 0.100 | - |
| | | Light structure (multiple) | -2.166 | 1.173 | 3.411 | 1 | 0.0648 | - |
| | Mean of turning angle, in 2025 | Light structure * Background light level (multiple, high) | -0.0565 | 0.118 | 0.245 | 1 | 0.620 | - |
| | | Trial number (two) | -0.0169 | 0.0579 | 0.103 | 1 | 0.749 | - |
| | | Capture method (trap) | 0.0611 | 0.0669 | 0.774 | 1 | 0.379 | - |
| | | Light structure (multiple) | -0.101 | 0.0949 | 3.267 | 1 | 0.0707 | - |
| | | *Background light level (high)* | *-0.201* | *0.0662* | *8.875* | *1* | *0.00289* | - |

| | Model | Factor | Estimate | SE | $X^2$ | df | p | Tukey's post-hoc tests |
|---|---|---|---|---|---|---|---|---|
| **Experiment 2** | SD of turning angle | Light structure * Background light level | - | - | 2.174 | 2 | 0.337 | - |
| | | Trial number (two) | 0.897 | 1.1536 | 0.764 | 1 | 0.382 | - |
| | | Capture method (trap) | 0.931 | 1.154 | 0.599 | 1 | 0.439 | - |
| | | *Light structure (multiple)* | *-3.121* | *1.030* | *9.065* | *1* | *0.00261* | - |
| | | *Background light level* | - | - | *10.078* | *2* | *0.00648* | *High – Low: Est. = -4.155, p = 0.00399*<br>High – Medium: Est. = -2.223, p = 0.253<br>Medium – Low: Est. = -1.932, p = 0.277 |

**<u>Supplementary Table S5.</u> Model outputs for analyses of flight tortuosity before and after exposure to light, in experiment 2.** Estimates (Est.) and standard error (SE) not provided for factors with more than 2 levels. The model for mean turning angle in 2024 includes only flight ID as a random effect; all other models include moth species, family and flight ID. Standard deviation (SD) of turning angle is square-root transformed to meet model assumptions. Factor reference levels are trial one, net collection, single focal light and low background light level for trial number, capture method, focal light structure and background light level respectively. For the SD of turning angle, Tukey's post-hoc tests suggest a significant effect of exposure to light for all background light levels, but smaller under the highest one: $\text{Estimate}_{\text{Low}} = 1.535$, $p_{\text{Low}} < 0.001$, $\text{Estimate}_{\text{Medium}} = 1.602$, $p_{\text{Medium}} < 0.001$, $\text{Estimate}_{\text{High}} = 0.508$, $p_{\text{High}} = 0.0046$. Significant effects are highlighted in italics.

| Model | Factor | Estimate | SE | $X^2$ | df | p |
|---|---|---|---|---|---|---|
| Mean turning angle, in 2024 | Background light level * Light structure * Exposure | -3.389 | 4.182 | 0.675 | 1 | 0.411 |
| | Light structure * Exposure | -0.112 | 2.088 | 0.003 | 1 | 0.957 |
| | Light structure * Background light level | 1.486 | 2.234 | 0.463 | 1 | 0.496 |
| | Background light level * Exposure | -3.382 | 2.069 | 2.690 | 1 | 0.101 |
| | Trial number (two) | -0.208 | 1.073 | 0.0386 | 1 | 0.844 |
| | Light structure (multiple) | -0.560 | 1.054 | 0.289 | 1 | 0.591 |
| | Capture method (trap) | 0.841 | 1.077 | 0.620 | 1 | 0.431 |
| | Background light level (medium) | -1.879 | 1.0402 | 3.259 | 1 | 0.0711 |
| | *Exposure to light* | *13.489* | *1.040* | *130.54* | *1* | *< 0.001* |
| Mean turning angle, in 2025 | Background light level * Light structure * Exposure | 6.924 | 4.440 | 2.496 | 1 | 0.114 |
| | Trial number (two) | -0.582 | 1.308 | 0.232 | 1 | 0.630 |
| | Light structure * Background light level | -0.289 | 2.631 | 0.0116 | 1 | 0.914 |
| | Capture method (trap) | 1.114 | 1.444 | 0.544 | 1 | 0.461 |
| | Light structure * Exposure | 1.079 | 2.152 | 0.258 | 1 | 0.612 |
| | *Light structure (multiple)* | *-3.083* | *1.250* | *5.892* | *1* | *0.0152* |
| | *Background light level * Exposure* | *-7.563* | *2.219* | *11.232* | *1* | *< 0.001* |
| SD of turning angle, all data | Background light level * Light structure * Exposure | - | - | 2.110 | 2 | 0.348 |
| | Trial number (two) | -0.0350 | 0.0996 | 0.133 | 1 | 0.716 |
| | Light structure * Background light level | - | - | 0.855 | 2 | 0.652 |
| | Capture method (trap) | 0.0475 | 0.106 | 0.157 | 1 | 0.692 |
| | Light structure * Exposure | 0.128 | 0.190 | 0.461 | 1 | 0.497 |
| | *Light structure (multiple)* | *-0.343* | *0.0959* | *12.473* | *1* | *< 0.001* |
| | *Background light level * Exposure* | - | - | *24.816* | *2* | *< 0.001* |
