## Supplementary data and code for "Effects of artificial light colour, intensity, structure and contrast on moth flight behaviour": Analysis_Script.R

```
#Analysis script for: "Effects of artificial light colour, intensity, structure and contrast on moth flight behaviour"

#18/03/2026

#load libraries
library(tidyverse)
library(ggplot2)
library(stringr)
library(lubridate)
library(patchwork)
library(ggmosaic)
library(ggpattern)
library(sjPlot)
library(lme4)
library(DHARMa)
library(glmmTMB)
library(emmeans)
library(multcomp)

#EXPERIMENT 1 - SINGLE LIGHTS----

#Experiment 1 - trial information----

INFO_SINGLE<-read.csv("Experiment1_MothInfo.csv", header=TRUE)
str(INFO_SINGLE)#832 trials

#separate treatment into focal light spectrum and intensity
for (i in 1:length(INFO_SINGLE$Treatment)){
  INFO_SINGLE$Spectrum[i]<-str_split(INFO_SINGLE$Treatment, "_")[[i]][1]
  INFO_SINGLE$Intensity[i]<-str_split(INFO_SINGLE$Treatment, "_")[[i]][2]
}
#set variables to factors where needed
factors <- c("Unique_Moth_ID","MothID", "Treatment","LatinName","Family",
             "CaptureMethod","Location","TrialNumber","Year","Spectrum", "Intensity", "Type")

#recode variables as needed
INFO_SINGLE<-INFO_SINGLE%>%
  mutate(Spectrum=recode(Spectrum,
                               "NA"="Narrow Amber","BA"="Broad Amber","BW" ="White"))%>%
  mutate(across(all_of(factors), as.factor))%>%
  mutate(Spectrum = factor(Spectrum,
                                levels = c("Control","White","Broad Amber","Narrow Amber")))%>%
  mutate(Intensity = factor(Intensity,
                           levels = c("0.3lx","3lx","30lx")))%>%
  mutate(Treatment = factor(Treatment,
                            levels = c("Control","Control_3lx","BW_0.3lx","BW_3lx","BW_30lx",
                                       "BA_0.3lx","BA_3lx","BA_30lx",
                                       "NA_0.3lx","NA_3lx","NA_30lx")))

str(INFO_SINGLE)
levels(INFO_SINGLE$Treatment)

#remove additional "control 3 lx" treatment from main dataset
#and create supplementary dataset with only controls from 2025
SUPP_SINGLE<-INFO_SINGLE%>%
  filter(Spectrum=="Control")%>%
  filter(Year=="2025")%>%
  droplevels()
str(SUPP_SINGLE)

#exclude 3lx controls from main dataset
INFO_SINGLE<-INFO_SINGLE%>%
  filter(Treatment!="Control_3lx")%>%
  droplevels()
dim(INFO_SINGLE)#808 trials now - there were 24 trials in 3lx control group

#remove trials with unknown species (N=2)
INFO_SINGLE<-INFO_SINGLE%>%
  filter(Family!="Unknown")%>%
  droplevels()
dim(INFO_SINGLE)#806 trials for analysis
head((INFO_SINGLE))

#count number of species & families in experiment 1
Expt1Species<-unique(INFO_SINGLE$LatinName)
Expt1Family<-unique(INFO_SINGLE$Family)
length(Expt1Species)
length(Expt1Family) #55 species of 6 families tested

#sample sizes per treatment and year - see supplementary material
INFO_SINGLE%>%
  group_by(Treatment, Year)%>%
  summarise(n=n())
INFO_SINGLE%>%
  group_by(Treatment)%>%
  summarise(n=n())

#Experiment 1 - track data----
SINGLETRACKS<-read.csv("Experiment1_AnalysedTracks.csv", header=TRUE)
head(SINGLETRACKS)
str(SINGLETRACKS)
unique(SINGLETRACKS$Unique_Moth_ID)#309 moths with tracked flight paths

#Clean data

#1) remove tracks from unknown species
SINGLETRACKS<-SINGLETRACKS%>%
  filter(Family!="Unknown")%>%
  droplevels()
unique(SINGLETRACKS$Unique_Moth_ID)#308 - 1 track from an unknown species removed

#2) remove tracks that do not cross focal light path

#identify flights that do not contain points in light beam for both cameras
NOLIGHT<-SINGLETRACKS%>%
  group_by(Unique_Moth_ID)%>%
  summarise(PointsInLight=sum(After_Light_Both))%>%
  filter(PointsInLight==0)
length(unique(NOLIGHT$Unique_Moth_ID))#36 flights to remove

#remove those unique flights from dataset
SINGLETRACKS_LIGHT<-SINGLETRACKS%>%
  filter(!(Unique_Moth_ID %in% NOLIGHT$Unique_Moth_ID))
unique(SINGLETRACKS_LIGHT$Unique_Moth_ID)#272 flights remain

#3) check that tracks start and end according to rules
  #ie. check that each track is a single uninterrupted flight, where the track does not continue after the moth lands somewhere

#to do this, we first identify any tracks with 3 or more points in the same position, in camera A or B
#then check the footage manually to verify whether the track is correct, or requires trimming

SINGLETRACKS_LIGHT_CHECK_A_B <- SINGLETRACKS_LIGHT %>%
  group_by(Unique_Moth_ID) %>% #group by moth track
  arrange(Frame) %>% #arrange frames in order (this should already be the case)
  mutate(Euclidean_distance_A = sqrt((A_X-lag(A_X))^2+ (A_Y-lag(A_Y))^2)) %>% #calculate euclidean distance between consecutive points in the track from camera A
  mutate(Stop_A = ifelse(Euclidean_distance_A==0, 1, 0)) %>% #identify points where moth does not move
  mutate(Counter_A = Stop_A + lag(Stop_A,1)) %>% #look for two instances of 0 movement (this implies 3 consecutive points in same position)
  mutate(Euclidean_distance_B = sqrt((B_X-lag(B_X))^2+ (B_Y-lag(B_Y))^2)) %>% #repeat for track from camera B
  mutate(Stop_B = ifelse(Euclidean_distance_B==0, 1, 0)) %>%
  mutate(Counter_B = Stop_B + lag(Stop_B,1)) %>%
  filter(any(Counter_A>=2|Counter_B>=2)) %>% #restrict to cases where conditions are met in tracks from A or B
  ungroup()
unique(SINGLETRACKS_LIGHT_CHECK_A_B$Unique_Moth_ID) #list moths with potential issues
  #21 moths to check
  #of these, 8 were deemed to not need trimming (moths did not land)
  #for the others, apply trimming based on frame numbers for stop counts, and manual checks of the footage
  #also checked that scored outcomes match new end points of tracks

SINGLETRACKS_LIGHT<-SINGLETRACKS_LIGHT%>%
  mutate(trim = case_when( #create new variable, trim, with trim = 1 if frame should be removed, else trim=0
    Unique_Moth_ID == "M20_2025" & Frame>42 ~ 1,
    Unique_Moth_ID == "M60_2025" & Frame>78 ~ 1,
    Unique_Moth_ID == "M19_2025" & Frame>81 ~ 1,
    Unique_Moth_ID == "M316_2025" & Frame>78 ~ 1,
    Unique_Moth_ID == "M51_2025" & Frame>122 ~ 1,
    Unique_Moth_ID == "M381_2025" & Frame>257 ~ 1,
    Unique_Moth_ID == "M287_2025" & Frame<190 ~ 1,
    Unique_Moth_ID == "M677_2025" & Frame<249 ~ 1,
    Unique_Moth_ID == "M290_2025" & Frame>305 ~ 1,
    Unique_Moth_ID == "M660_2025" & Frame<445 ~ 1,
    Unique_Moth_ID == "M660_2025" & Frame>475 ~ 1,
    Unique_Moth_ID == "M105_2025" & Frame<633 ~ 1,
    Unique_Moth_ID == "M172_2024" & Frame>171 ~ 1,
    Unique_Moth_ID == "M203_2024" & Frame>323 ~ 1,
    .default = 0
  ))%>%
  filter(trim==0)

#check they all still cross light path, and are long enough for analysis
NOLIGHT2<-SINGLETRACKS_LIGHT%>%
  group_by(Unique_Moth_ID)%>%
  summarise(PointsInLight=sum(After_Light_Both))%>%
  filter(PointsInLight==0)
length(unique(NOLIGHT2$Unique_Moth_ID))#0, so all tracks still cross light path
SINGLETRACKS_LIGHT%>%
  group_by(Unique_Moth_ID)%>%
  summarise(n=n())%>%
  filter(n<5)#all over 5 points long

#4) identify any tracks with overly large displacements, likely indicating tracking errors

#calculate displacement (in 3D) between points, per moth track
TEMPTRACKS<-list()

for (i in 1: length(unique(SINGLETRACKS_LIGHT$Unique_Moth_ID))){
  
  mothid<-unique(SINGLETRACKS_LIGHT$Unique_Moth_ID)[i]
  
  TEMPTRACKS[[i]]<-SINGLETRACKS_LIGHT%>%
    filter(Unique_Moth_ID ==mothid)%>%
    arrange(Frame)
  TEMPTRACKS[[i]]$Displacement<-rep(0, length(TEMPTRACKS[[i]]$X))
  
  for (j in 2:length(TEMPTRACKS[[i]]$X)){
    TEMPTRACKS[[i]]$Displacement[j]<-sqrt((TEMPTRACKS[[i]]$X_smooth[j]-TEMPTRACKS[[i]]$X_smooth[j-1])^2+
                                            (TEMPTRACKS[[i]]$Y_smooth[j]-TEMPTRACKS[[i]]$Y_smooth[j-1])^2+
                                            (TEMPTRACKS[[i]]$Z_smooth[j]-TEMPTRACKS[[i]]$Z_smooth[j-1])^2)
  }
}
#recombine into single dataframe
SINGLETRACKS_LIGHT<-do.call("rbind", TEMPTRACKS)

#identify tracks with odd displacements
hist(SINGLETRACKS_LIGHT$Displacement, breaks=100)
summary(SINGLETRACKS_LIGHT$Displacement)
LARGE<-SINGLETRACKS_LIGHT%>%
  filter(Displacement> 0.5)#11 flights - this matches what visual inspection of tracks suggests
unique(LARGE$Unique_Moth_ID)
  #keep these in dataset for the moment, for analysis of flight outcomes etc.
  #but the error-prone segments should be removed for tortuosity analysis

#data cleaning done

#Recode variables as for info dataset above
for (i in 1:length(SINGLETRACKS_LIGHT$Treatment)){
  SINGLETRACKS_LIGHT$Spectrum[i]<-str_split(SINGLETRACKS_LIGHT$Treatment, "_")[[i]][1]
  SINGLETRACKS_LIGHT$Intensity[i]<-str_split(SINGLETRACKS_LIGHT$Treatment, "_")[[i]][2]
}
factors <- c("Unique_Moth_ID","Moth_ID", "Treatment","LatinName","Family",
             "CaptureMethod","Location","Year","Spectrum", "Intensity")
SINGLETRACKS_LIGHT<-SINGLETRACKS_LIGHT%>%
  mutate(across(all_of(factors), as.factor))%>%
  mutate(Treatment = factor(Treatment,
                            levels = c("Control","BW_0.3lx","BW_3lx","BW_30lx",
                                       "BA_0.3lx","BA_3lx","BA_30lx",
                                       "NA_0.3lx","NA_3lx","NA_30lx","Control_3lx")))
str(SINGLETRACKS_LIGHT)

#Remove Control 3lx light treatment
SUPP_SINGLETRACKS<-SINGLETRACKS_LIGHT%>%
  filter(Treatment=="Control_3lx")%>%
  droplevels()
str(SUPP_SINGLETRACKS)
length(unique(SUPP_SINGLETRACKS$Unique_Moth_ID))#only 2 flights meet criteria - too few for analysis

SINGLETRACKS_LIGHT<-SINGLETRACKS_LIGHT%>%
  filter(Treatment!="Control_3lx")%>%
  droplevels()
str(SINGLETRACKS_LIGHT)

#Key numbers

#number of flights:
length(unique(SINGLETRACKS_LIGHT$Unique_Moth_ID))#270 flights to analyse 
SINGLETRACKS_LIGHT%>%
  group_by(Treatment, Unique_Moth_ID)%>%
  summarise(n=n())%>%
  group_by(Treatment)%>%
  summarise(n=n())#18 to 45 flights per treatment have 3D tracks

#distribution of families between treatments:
FAMILIES<-SINGLETRACKS_LIGHT%>%
  group_by(Family,Treatment, Unique_Moth_ID)%>%
  summarise(n=n())%>%
  group_by(Family, Treatment)%>%
  summarise(n=n())
ggplot(FAMILIES, aes(fill=Family, y=n, x=Treatment, label=n)) + 
  geom_bar(position="fill", stat="identity")+
  theme_bw()+ylab("Proportion of flights")+
  geom_text(size = 3, position = position_fill(vjust = 0.5))
  #family composition is similar across treatment groups, although more noctuids in control

#number of different species tracked:
SINGLETRACKSPECIES<-unique(SINGLETRACKS_LIGHT$LatinName)#34 species
SINGLETRACKSPECIES
unique(SINGLETRACKS_LIGHT$Family)#5 families

#Experiment 1 - results ----

#1) Flight probability----

#Key numbers
INFO_SINGLE%>%
  group_by(TakeOff)%>%
  summarise(n=n()) #806 flight attempts; of these, 5 excluded, 407 did not fly within 1 minute, 394 took off

#Plot proportion of successful take-offs

#summarise by treatment
SUMMARY_INFO_SINGLE<-INFO_SINGLE%>%
  filter(!is.na(TakeOff))%>%#exclude NAs
  group_by(Treatment,TakeOff)%>%
  summarise(n=n())%>%
  ungroup()%>%
  mutate(TakeOff = factor(TakeOff,
                          levels = c("0","1")))%>%
  mutate(TreatmentTO = paste(Treatment,TakeOff))#composite variable for plotting

#create colour lists for plotting
SUMMARY_INFO_SINGLE$TreatmentTO<-factor(SUMMARY_INFO_SINGLE$TreatmentTO,
                                        levels = c("Control 0",  "Control 1",
                                                   "BW_0.3lx 0", "BW_0.3lx 1",
                                                   "BW_3lx 0" ,  "BW_3lx 1",
                                                   "BW_30lx 0",  "BW_30lx 1",
                                                   "BA_0.3lx 0", "BA_0.3lx 1",
                                                   "BA_3lx 0","BA_3lx 1",
                                                   "BA_30lx 0" , "BA_30lx 1",
                                                   "NA_0.3lx 0","NA_0.3lx 1",
                                                   "NA_3lx 0" ,  "NA_3lx 1",
                                                   "NA_30lx 0" , "NA_30lx 1"))
levels(SUMMARY_INFO_SINGLE$TreatmentTO)
expt1colours<-c("#ab7fff","#afddfa","#64c1fa","#19a4fa","#ffe4b2",
                "#ffc966","#ffae1a","#ffc7b2","#ff8f66","#ff571a")
expt1colours2<-c("white","#ab7fff","white","#afddfa","white","#64c1fa","white","#19a4fa","white","#ffe4b2",
                 "white","#ffc966","white","#ffae1a","white","#ffc7b2","white","#ff8f66","white","#ff571a")
plot_TO_expt1<-ggplot(SUMMARY_INFO_SINGLE, aes(fill=TreatmentTO, col=Treatment, y=n, x=Treatment, label=n)) + 
  geom_bar(position="fill", stat="identity")+
  ylab("Flight success (% trials)")+xlab("Focal light treatment")+
  scale_y_continuous(expand=c(0,0),breaks=c(0, 0.25, 0.5, 0.75, 1), labels= c(0,25,50,75,100))+
  scale_x_discrete(labels = c("Control", "White 0.3 lx",
                              "White 3 lx", "White 30 lx", "BA 0.3 lx",
                              "BA 3 lx","BA 30 lx","NA 0.3 lx",
                              "NA 3 lx","NA 30 lx"))+
  scale_fill_manual(values =expt1colours2)+
  scale_color_manual(values=c(expt1colours))+
  geom_text(size = 3, position = position_fill(vjust = 0.5), col="grey40")+
  theme(legend.text = element_text(size=10),
        legend.title = element_blank())+
  theme_classic()+
  guides(fill="none")+
  guides(fill="none")+
  ggtitle("A")+
  theme(axis.title.x = element_text(size = 12),
        axis.text.x = element_text(angle = 45, hjust = 1, size=10),
        axis.title.y = element_text(size = 12),
        axis.text.y=element_text(size=10),
        axis.title=element_text(size=16))
plot_TO_expt1

#model probability of take-off

#a) Is there a difference between years (for repeated treatments)?

#create dataset with only treatments repeated in 2024 and 2025
YEARTEST<-INFO_SINGLE%>%
  filter(Treatment =="Control" |Intensity=="30lx")%>% 
  droplevels()%>%
  mutate(TakeOff = factor(TakeOff,levels = c("0","1")))
str(YEARTEST)

#plot
SUMMARY_YEARTEST<-YEARTEST%>%
  filter(!is.na(TakeOff))%>%
  group_by(Treatment,Year, TakeOff)%>%
  summarise(n=n())%>%
  ungroup()
ggplot(SUMMARY_YEARTEST, aes(fill=TakeOff, y=n, x=Year, label=n)) + 
  geom_bar(position="fill", stat="identity")+
  facet_wrap(.~Treatment)+
  scale_fill_manual(values = c("grey70","mediumpurple1"))+
  theme_classic()+ylab("Proportion of flights")+
  geom_text(size = 3, position = position_fill(vjust = 0.5))

#model
year_mod1<-glmmTMB(TakeOff ~ Treatment*Year+CaptureMethod+
                     (1|LatinName)+(1|Family),
                   family=binomial, data=YEARTEST)
summary(year_mod1)
year_mod1_resid <- simulateResiduals(year_mod1)
plot(year_mod1_resid)#good

year_mod1b<-glmmTMB(TakeOff ~ Treatment+CaptureMethod+Year+
                      (1|LatinName)+(1|Family),
                    family=binomial, data=YEARTEST)
anova(year_mod1, year_mod1b)#can remove interaction
summary(year_mod1b)

year_mod1c<-glmmTMB(TakeOff ~ Treatment+CaptureMethod+
                      (1|LatinName)+(1|Family),
                    family=binomial, data=YEARTEST)
anova(year_mod1c, year_mod1b)#can remove year
summary(year_mod1c)

#b) across both years, effect of treatment?

#model with treatment only
prob_mod1<-glmmTMB(TakeOff ~ Treatment+CaptureMethod+
                     (1|LatinName)+(1|Family),
                   family=binomial, data=INFO_SINGLE)
summary(prob_mod1)
prob_mod1_resid <- simulateResiduals(prob_mod1)
plot(prob_mod1_resid)#good

prob_mod1b<-glmmTMB(TakeOff ~ Treatment+
                      (1|LatinName)+(1|Family),
                    family=binomial, data=INFO_SINGLE)
anova(prob_mod1, prob_mod1b)#can't remove capture method

prob_mod1c<-glmmTMB(TakeOff ~ CaptureMethod+(1|LatinName)+(1|Family),
                    family=binomial, data=INFO_SINGLE)
anova(prob_mod1c, prob_mod1)#can remove treatment
summary(prob_mod1c)

prob_mod1d<-glmmTMB(TakeOff ~ 1+(1|LatinName)+(1|Family),
                    family=binomial, data=INFO_SINGLE)
anova(prob_mod1c, prob_mod1d)#significant effect of capture method

#odds ratios
exp(-0.7)#odds ratio 0.497, odds of taking off are reduced by half when moths are captured by traps

#plot binomial model results

plot_TO_expt1_capture<-plot_model(prob_mod1c, type = "pred", 
                                  terms = c("CaptureMethod"),
                                  title = "E",
                                  colors = "darkgreen")+
  theme_classic()+
  scale_y_continuous(limits=c(0.1,0.8),breaks = c(0.2, 0.4, 0.6,0.8), labels = c(20,40,60,80))+  ylab("Predicted flight probability (%)")+xlab("Capture method")+
  theme(axis.title.x = element_text(size = 12),
        axis.title = element_text(size=14),
        axis.text.x = element_text(size = 10),
        axis.title.y = element_text(size = 12),
        axis.text.y=element_text(size=10))
plot_TO_expt1_capture

#check random effects - family has very small effect, but species worth looking into
#random effects
re<-ranef(prob_mod1c)
print(re, simplify=FALSE)

str(re$cond$LatinName)
re_species<-re$cond$LatinName
re_species$Species<-row.names(re_species)
colnames(re_species)[1]<-"Intercept"
re_species %>%
  mutate(Species = fct_reorder(Species, Intercept)) %>%
  ggplot( aes(x=Intercept, y=Species)) +
  geom_point()+ theme_classic()+
  geom_vline(xintercept=0, col="red")

#2) Outcomes - flight to light----

#outcomes of flights with 3D tracks
  #use INFO_SINGLE and restrict to only analysed tracks in SINGLETRACKS_LIGHT
OUTCOMES_SINGLE_3D<-INFO_SINGLE%>%
  filter(Unique_Moth_ID %in% SINGLETRACKS_LIGHT$Unique_Moth_ID)
dim(OUTCOMES_SINGLE_3D)#270 flights
str(OUTCOMES_SINGLE_3D)

#include 2D flights - how many fly past the light?
ANALYSIS_2DONLY<-INFO_SINGLE%>%
  filter(Type=="2D")%>%
  filter(TakeOff==1)
dim(ANALYSIS_2DONLY)#57 successful flights filmed in 2D
OUTCOMES_2DONLY<-INFO_SINGLE%>%
  filter(Type=="2D")%>%
  filter(!is.na(TrackedFlight_Outcome))
dim(OUTCOMES_2DONLY)#32 flights where moth passes light

#combine 2D and 3D flights that meet criteria for outcome analysis
OUTCOMES_EXPT1<-rbind(OUTCOMES_2DONLY,OUTCOMES_SINGLE_3D)
dim(OUTCOMES_EXPT1)#302

#key numbers:
OUTCOMES_EXPT1%>%
  group_by(Treatment)%>%
  summarise(n=n())#18 to 57 per treatment in experiment 1

#break down by year, 2D/3D and treatment - see supplementary material
OUTCOMES_EXPT1%>%
  group_by(Treatment, Type, Year)%>%
  summarise(n=n())

#proportion of flights ending at light:
OUTCOMES_EXPT1%>%
  group_by(FlighttoLight_tracked)%>%
  summarise(n=n())%>%
  ungroup()
71/302#24% flights ended at the light

#overall proportion of flights ending up:
OUTCOMES_EXPT1%>%
  group_by(Up_tracked)%>%
  summarise(n=n())%>%
  ungroup()
179/302#60% of tracked fligths end with the moth flying upwards

#overall proportion of flights ending down:
OUTCOMES_EXPT1%>%
  group_by(Down_tracked)%>%
  summarise(n=n())%>%
  ungroup()
21/(302)#7% 

#overall proportion of flights with erratic spiraling:
OUTCOMES_EXPT1%>%
  group_by(Erratic_spiralling)%>%
  summarise(n=n())%>%
  ungroup()
52/(302)#17% flights with spirals

#plot outcomes - see figure 2D

OUTCOMES_EXPT1<-OUTCOMES_EXPT1%>%
  mutate(Treatment = factor(Treatment,
                            levels = c("Control","BW_0.3lx","BW_3lx","BW_30lx",
                                       "BA_0.3lx","BA_3lx","BA_30lx",
                                       "NA_0.3lx","NA_3lx","NA_30lx")))%>%
  mutate(Outcome_simpler = factor(Outcome_simpler,
                          levels = c("at light", "up","down", "other")))%>%
  droplevels()

OUTCOME_SUMMARY<-OUTCOMES_EXPT1%>%
  group_by(Treatment, Outcome_simpler)%>%
  summarise(n=n())%>%
  mutate(Outcome_simpler = factor(Outcome_simpler,
                            levels = c("other","down","up","at light")))%>%
   ungroup()

outcomes_single<-ggplot(OUTCOME_SUMMARY, aes(fill=Outcome_simpler, y=n, x=Treatment, label=n)) + 
  geom_bar(position="fill", stat="identity")+
  theme_bw()+
  geom_text(size = 3, position = position_fill(vjust = 0.5))+
  labs(fill="Flight outcome")+
  ylab("Flight outcomes (% trials)")+
  scale_y_continuous(breaks=c(0,0.25, 0.5, 0.75, 1), labels = c(0,25,50,75,100))+
  ggtitle("D")+
  scale_fill_manual(values = c("#B3B3B3","#66C2A5","#E78AC3","#FFD92F"))+
  geom_text(size = 3, position = position_fill(vjust = 0.5), col="grey50")+  
  theme(legend.text = element_text(size=10))+
  theme_classic()+
  theme(axis.title.x = element_text(size = 12),
        axis.title = element_text(size=16),
        axis.text.x = element_text(angle = 45, hjust = 1, size=10),
        axis.title.y = element_text(size = 12),
        axis.text.y=element_text(size=10))
  #+guides(fill="none")
outcomes_single

#a) test for difference between years

OUTCOMES_YEAR<-OUTCOMES_EXPT1%>%
  filter(Treatment=="Control"|Treatment=="BW_30lx"|Treatment=="BA_30lx"|Treatment=="NA_30lx")%>%
  droplevels()
  #model including family level cannot be computed once treatment*year has been removed,
  #so switch to model without family, which has low explaining power (variance estimate is 10^-19)
year_mod2<-glmmTMB(FlighttoLight_tracked ~ Treatment*Year+CaptureMethod+
                     (1|LatinName),
                   family=binomial, data=OUTCOMES_YEAR)
summary(year_mod2)
year_mod2_resid <- simulateResiduals(year_mod2)
plot(year_mod2_resid)#ok
year_mod2b<-glmmTMB(FlighttoLight_tracked ~ Treatment+Year+CaptureMethod+
                      (1|LatinName),
                    family=binomial, data=OUTCOMES_YEAR)

anova(year_mod2, year_mod2b)#remove interaction
summary(year_mod2b)
year_mod2c<-glmmTMB(FlighttoLight_tracked ~ Treatment+CaptureMethod+
                      (1|LatinName),
                    family=binomial, data=OUTCOMES_YEAR)

anova(year_mod2c, year_mod2b)#remove year
summary(year_mod2c)

#b) model effect of treatment across both years

with(OUTCOMES_EXPT1,
     table(FlighttoLight_tracked, Treatment))
#BA0.3lx has no flight to light, which leads to erroneous estimates in the model fit (very large values, with large standard error)
#so remove this treatment from the analysis
#also do not test for treatment*capture interaction, as two few numbers in many combinations, again leading to very large estimates and se

OUTCOMES_EXPT1_FTL<-OUTCOMES_EXPT1%>%
  filter(Treatment!="BA_0.3lx")%>%
  droplevels()

ftl_mod1<-glmmTMB(FlighttoLight_tracked ~ Treatment+CaptureMethod+
                    (1|LatinName)+(1|Family),
                  family=binomial, data=OUTCOMES_EXPT1_FTL)
summary(ftl_mod1)
ftl_mod1_resid <- simulateResiduals(ftl_mod1)
plot(ftl_mod1_resid)#good
ftl_mod1b<-glmmTMB(FlighttoLight_tracked ~ CaptureMethod+
                    (1|LatinName)+(1|Family),
                  family=binomial, data=OUTCOMES_EXPT1_FTL)
anova(ftl_mod1, ftl_mod1b)#effect of treatment
ftl_mod1c<-glmmTMB(FlighttoLight_tracked ~ Treatment+
                     (1|LatinName)+(1|Family),
                   family=binomial, data=OUTCOMES_EXPT1_FTL)
anova(ftl_mod1, ftl_mod1c)#effect of capture method
summary(ftl_mod1)
summary(glht(ftl_mod1, linfct = mcp(Treatment = "Tukey"))) 
#only white 30 lx differs from the control

#c) model effects of focal light spectrum and intensity
  #exclude control treatment, but not BA0.3lx

EXCLCONTROL_EXPT1<-OUTCOMES_EXPT1%>%
  filter(Treatment!="Control")%>%
  droplevels()
str(EXCLCONTROL_EXPT1)

#cannot test interaction of spectrum and intensity, leads to very large estimates for broad amber
ftl_mod2<-glmmTMB(FlighttoLight_tracked ~ Spectrum+Intensity+CaptureMethod+
                    (1|LatinName)+(1|Family),
                  family=binomial, data=EXCLCONTROL_EXPT1)
summary(ftl_mod2)
ftl_mod2_resid <- simulateResiduals(ftl_mod2)
plot(ftl_mod2_resid)#good

ftl_mod2b<-glmmTMB(FlighttoLight_tracked ~ Intensity+CaptureMethod+
                    (1|LatinName)+(1|Family),
                  family=binomial, data=EXCLCONTROL_EXPT1)
anova(ftl_mod2, ftl_mod2b)#remove spectrum
summary(ftl_mod2b)

ftl_mod2c<-glmmTMB(FlighttoLight_tracked ~ CaptureMethod+
                     (1|LatinName)+(1|Family),
                   family=binomial, data=EXCLCONTROL_EXPT1)
anova(ftl_mod2c, ftl_mod2b)#significant effect of intensity

ftl_mod2d<-glmmTMB(FlighttoLight_tracked ~ Intensity+
                     (1|LatinName)+(1|Family),
                   family=binomial, data=EXCLCONTROL_EXPT1)
anova(ftl_mod2b, ftl_mod2d)#significant effect of capture method

summary(ftl_mod2b)
#looks like significant effects of intensity and capture method
exp(0.851)#odds of flight to light 2.342 times as high for trapped moths
summary(glht(ftl_mod2b, linfct = mcp(Intensity = "Tukey")))
#for intensity, lowest intensity significantly differs from highest, 
#other pairwise comparisons close to significance 

#plot effects
plot_ftl_expt1_intensity<-plot_model(ftl_mod2b, type = "pred", 
                                   terms = c("Intensity"),
                                   colors = c("black","grey40","grey80"),
                                   title="F",
                                   axis.labels = c("0.3 lx", "3 lx", "30 lx"))+
  theme_classic()+
  scale_y_continuous(limits=c(0,0.9),breaks = c(0,0.2, 0.4, 0.6,0.8), 
                     labels = c(0,20,40,60,80),expand=c(0,0))+
  xlab("Focal light intensity")+ylab("Predicted probability of flight-to-light (%)")+
  theme(axis.title.x = element_text(size = 12),
        axis.title = element_text(size=16),
        axis.text.x = element_text(size = 10),
        axis.title.y = element_text(size = 12),
        axis.text.y=element_text(size=10))
plot_ftl_expt1_intensity

plot_ftl_expt1_capture<-plot_model(ftl_mod2b, type = "pred", 
                                   terms = c("CaptureMethod"),
                                   colors = "darkgreen",
                                   title="G")+
  theme_classic()+
  scale_y_continuous(limits=c(0,0.9),breaks = c(0,0.2, 0.4, 0.6,0.8), 
                     labels = c(0,20,40,60,80), expand =c(0,0))+
  xlab("Capture method")+ylab("Predicted probability of flight-to-light (%)")+
  theme(axis.title.x = element_text(size = 12),
        axis.title = element_text(size=16),
        axis.text.x = element_text(size = 10),
        axis.title.y = element_text(size = 12),
        axis.text.y=element_text(size=10))
plot_ftl_expt1_capture

#3) Outcomes- up----

#a)test year difference
year_mod3<-glmmTMB(Up_tracked ~ Treatment*Year+CaptureMethod+
                     (1|LatinName)+(1|Family),
                   family=binomial, data=OUTCOMES_YEAR)
summary(year_mod3)
year_mod3_resid <- simulateResiduals(year_mod3)
plot(year_mod3_resid)#good
year_mod3b<-glmmTMB(Up_tracked ~ Treatment+Year+CaptureMethod+
                      (1|LatinName)+(1|Family),
                    family=binomial, data=OUTCOMES_YEAR)

anova(year_mod3, year_mod3b)#remove interaction
year_mod3c<-glmmTMB(Up_tracked ~ Treatment+CaptureMethod+
                      (1|LatinName)+(1|Family),
                    family=binomial, data=OUTCOMES_YEAR)

anova(year_mod2c, year_mod2b)#remove year

#b) model with treatment, across both years
up_mod1<-glmmTMB(Up_tracked ~ Treatment+CaptureMethod+
                    (1|LatinName)+(1|Family),
                  family=binomial, data=OUTCOMES_EXPT1)
summary(up_mod1)
up_mod1_resid <- simulateResiduals(up_mod1)
plot(up_mod1_resid)#good

up_mod1b<-glmmTMB(Up_tracked ~ CaptureMethod+
                   (1|LatinName)+(1|Family),
                 family=binomial, data=OUTCOMES_EXPT1)
anova(up_mod1, up_mod1b)#significant effect of treatment
summary(glht(up_mod1, linfct = mcp(Treatment = "Tukey"))) 
#all 30 lx light treatments differ from the control

#test effect of spectrum and intensity

up_mod2<-glmmTMB(Up_tracked ~ Spectrum*Intensity+CaptureMethod+
                    (1|LatinName)+(1|Family),
                  family=binomial, data=EXCLCONTROL_EXPT1)
summary(up_mod2)
up_mod2_resid <- simulateResiduals(up_mod2)
plot(up_mod2_resid)#good

up_mod2b<-glmmTMB(Up_tracked ~ Spectrum+Intensity+CaptureMethod+
                     (1|LatinName)+(1|Family),
                   family=binomial, data=EXCLCONTROL_EXPT1)
anova(up_mod2, up_mod2b)#remove interaction
summary(up_mod2b)

up_mod2c<-glmmTMB(Up_tracked ~ Intensity+CaptureMethod+
                     (1|LatinName)+(1|Family),
                   family=binomial, data=EXCLCONTROL_EXPT1)
anova(up_mod2c, up_mod2b)#remove spectrum
summary(up_mod2c)

up_mod2d<-glmmTMB(Up_tracked ~ CaptureMethod+
                    (1|LatinName)+(1|Family),
                  family=binomial, data=EXCLCONTROL_EXPT1)
anova(up_mod2c, up_mod2d)#significant effect of intensity

up_mod2e<-glmmTMB(Up_tracked ~ Intensity+
                    (1|LatinName)+(1|Family),
                  family=binomial, data=EXCLCONTROL_EXPT1)
anova(up_mod2c, up_mod2e)#capture method not quite significant 

up_mod2f<-glmmTMB(Up_tracked ~ 1+
                    (1|LatinName)+(1|Family),
                  family=binomial, data=EXCLCONTROL_EXPT1)
anova(up_mod2e, up_mod2f)#significant effect of light intensity

summary(glht(up_mod2e, linfct = mcp(Intensity = "Tukey")))
#30lx - 0.3lx and 3lx - 0.3lx comparisons are significant

plot_model(up_mod2e, type = "pred", terms = c("Intensity"))+
  theme_classic()+
  xlab("Focal light intensity")+ylab("Predicted probability of upwards flight")+
  ggtitle ("")+
  theme(axis.title.x = element_text(size = 12),
        axis.text.x = element_text(size = 10),
        axis.title.y = element_text(size = 12),
        axis.text.y=element_text(size=10))


#4) Erratic flight----

SPIRAL_SUMMARY<-OUTCOMES_EXPT1%>%
  group_by(Treatment, Erratic_spiralling)%>%
  summarise(n=n())%>%
  mutate(Erratic_spiralling = as.factor(Erratic_spiralling))%>%
  ungroup()

ggplot(SPIRAL_SUMMARY, aes(fill=Erratic_spiralling, y=n, x=Treatment, label=n)) + 
  geom_bar(position="fill", stat="identity")+
  theme_bw()+
  geom_text(size = 3, position = position_fill(vjust = 0.5))+
  labs(fill="Erratic spiralling")+
  ylab("Erratic spiralling (% trials)")+
  scale_y_continuous(breaks=c(0,0.25, 0.5, 0.75, 1), labels = c(0,25,50,75,100))+
  geom_text(size = 3, position = position_fill(vjust = 0.5), col="grey50")+  theme(legend.text = element_text(size=10))+
  theme_classic()+
  theme(axis.title.x = element_text(size = 12),
        axis.title = element_text(size=16),
        axis.text.x = element_text(angle = 45, hjust = 1, size=10),
        axis.title.y = element_text(size = 12),
        axis.text.y=element_text(size=10))

#no erratic spiraling in the control, so this makes it difficult to analyse
#skip straight to analysis with dataset excluding the control, testing effects of intensity and spectrum

#a)test year difference
EXCLCONTROL_EXPT1_YEAR<-EXCLCONTROL_EXPT1%>%
  filter(Intensity =="30lx")
head(EXCLCONTROL_EXPT1_YEAR)

year_mod4<-glmmTMB(Erratic_spiralling ~ Spectrum*Year+CaptureMethod+
                     (1|LatinName)+(1|Family),
                   family=binomial, data=EXCLCONTROL_EXPT1_YEAR) #only one intensity level tested both years, so here can only test spectrum * year
year_mod4_resid <- simulateResiduals(year_mod4)
plot(year_mod4_resid)#good

year_mod4b<-glmmTMB(Erratic_spiralling ~ Spectrum+Year+CaptureMethod+
                      (1|LatinName)+(1|Family),
                    family=binomial, data=EXCLCONTROL_EXPT1_YEAR)
anova(year_mod4, year_mod4b)#remove interaction

year_mod4c<-glmmTMB(Erratic_spiralling ~ Spectrum+CaptureMethod+
                      (1|LatinName)+(1|Family),
                    family=binomial, data=EXCLCONTROL_EXPT1_YEAR)
anova(year_mod4c, year_mod4b)#remove year


#b) model with intensity and spectrum, across both years

sp_mod1<-glmmTMB(Erratic_spiralling ~ Spectrum*Intensity+CaptureMethod+
                   (1|LatinName)+(1|Family),
                 family=binomial, data=EXCLCONTROL_EXPT1)
summary(sp_mod1)
sp_mod1_resid <- simulateResiduals(sp_mod1)
plot(sp_mod1_resid)#good

sp_mod1b<-glmmTMB(Erratic_spiralling ~ Spectrum+Intensity+CaptureMethod+
                    (1|LatinName)+(1|Family),
                  family=binomial, data=EXCLCONTROL_EXPT1)
anova(sp_mod1, sp_mod1b)#remove interaction
summary(sp_mod1b)

sp_mod1c<-glmmTMB(Erratic_spiralling ~ Intensity+CaptureMethod+
                    (1|LatinName)+(1|Family),
                  family=binomial, data=EXCLCONTROL_EXPT1)
anova(sp_mod1c, sp_mod1b)#remove spectrum
summary(sp_mod1c)
sp_mod1d<-glmmTMB(Erratic_spiralling ~ Intensity+
                    (1|LatinName)+(1|Family),
                  family=binomial, data=EXCLCONTROL_EXPT1)
anova(sp_mod1d, sp_mod1c)#remove capture method
summary(sp_mod1d)
sp_mod1e<-glmmTMB(Erratic_spiralling ~ 1+
                    (1|LatinName)+(1|Family),
                  family=binomial, data=EXCLCONTROL_EXPT1)
anova(sp_mod1d, sp_mod1e)#significant effect of intensity only
summary(glht(sp_mod1d, linfct = mcp(Intensity = "Tukey")))
#30 lx different from both 0.3 and 3

#plot model
plot_sp_expt1_intensity<-plot_model(sp_mod1d, type = "pred", 
                                    terms = c("Intensity"),
                                    colors = c("grey30","grey50","grey90"),
                                    title="C")+
  theme_classic()+
  scale_y_continuous(limits=c(0,0.5),breaks = c(0,0.2, 0.4), 
                     labels = c(0,20,40), expand =c(0,0))+
  xlab("Focal light intensity (lx)")+ylab("Predicted probability of erratic spiralling (%)")+
  theme(axis.title.x = element_text(size = 12),
        axis.title=element_text(size=16),
        axis.text.x = element_text(size = 10),
        axis.title.y = element_text(size = 12),
        axis.text.y=element_text(size=10))
plot_sp_expt1_intensity

#what about any form of spiraling?

#make variable for record of any spiral observed, and update datasets with it
OUTCOMES_EXPT1<-OUTCOMES_EXPT1%>%
  mutate(AnySpiral = case_when(
    Spiral_up ==1|Spiral_down==1|Spiral_other==1 ~1,
    .default=0))

OUTCOMES_YEAR<-OUTCOMES_EXPT1%>%
  filter(Treatment=="Control"|Treatment=="BW_30lx"|Treatment=="BA_30lx"|Treatment=="NA_30lx")%>%
  droplevels()  
EXCLCONTROL_EXPT1<-OUTCOMES_EXPT1%>%
  filter(Treatment!="Control")%>%
  droplevels()

ANYSPIRAL_SUMMARY<-OUTCOMES_EXPT1%>%
  group_by(Treatment, AnySpiral)%>%
  summarise(n=n())%>%
  mutate(AnySpiral= as.factor(AnySpiral))%>%
  ungroup()

OUTCOMES_EXPT1%>%
  group_by(AnySpiral)%>%
  summarise(n=n())%>%
  ungroup()
139/302# 47 % trials featured a spiral

ggplot(ANYSPIRAL_SUMMARY, aes(fill=AnySpiral, y=n, x=Treatment, label=n)) + 
  geom_bar(position="fill", stat="identity")+
  theme_bw()+
  geom_text(size = 3, position = position_fill(vjust = 0.5))+
  labs(fill="Spiralling")+
  ylab("Spiral observed (% trials)")+
  scale_y_continuous(breaks=c(0,0.25, 0.5, 0.75, 1), labels = c(0,25,50,75,100))+
  geom_text(size = 3, position = position_fill(vjust = 0.5), col="grey50")+  theme(legend.text = element_text(size=10))+
  theme_classic()+
  theme(axis.title.x = element_text(size = 12),
        axis.title = element_text(size=16),
        axis.text.x = element_text(angle = 45, hjust = 1, size=10),
        axis.title.y = element_text(size = 12),
        axis.text.y=element_text(size=10))

#this variable can be analysed with all data, including the control

#a) test difference between years
year_mod5<-glmmTMB(AnySpiral ~ Treatment*Year+CaptureMethod+
                     (1|LatinName)+(1|Family),
                   family=binomial, data=OUTCOMES_YEAR) 
year_mod5_resid <- simulateResiduals(year_mod5)
plot(year_mod5_resid)#good
summary(year_mod5)

year_mod5b<-glmmTMB(AnySpiral ~ Treatment+Year+CaptureMethod+
                     (1|LatinName)+(1|Family),
                   family=binomial, data=OUTCOMES_YEAR) 
anova(year_mod5, year_mod5b)#can just remove interaction
summary(year_mod5b)
year_mod5c<-glmmTMB(AnySpiral ~ Treatment+CaptureMethod+
                      (1|LatinName)+(1|Family),
                    family=binomial, data=OUTCOMES_YEAR)
anova(year_mod5c, year_mod5b) #can remove year

#b) model across years
sp_mod2<-glmmTMB(AnySpiral ~ Treatment+CaptureMethod+
                     (1|LatinName)+(1|Family),
                   family=binomial, data=OUTCOMES_EXPT1)
sp_mod2_resid <- simulateResiduals(sp_mod2)
plot(sp_mod2_resid)#good
summary(sp_mod2)

sp_mod2b<-glmmTMB(AnySpiral ~ Treatment+
                   (1|LatinName)+(1|Family),
                 family=binomial, data=OUTCOMES_EXPT1)
anova(sp_mod2, sp_mod2b)
summary(sp_mod2b)#sig effect of treatment only
summary(glht(sp_mod2b, linfct = mcp(Treatment = "Tukey")))
  #significant contrasts are all between 30 lx treatments and controls or 0.3 lx treatments

#c) model with intensity and spectrum, across both years

sp_mod3<-glmmTMB(AnySpiral ~ Spectrum*Intensity+CaptureMethod+
                   (1|LatinName)+(1|Family),
                 family=binomial, data=EXCLCONTROL_EXPT1)
summary(sp_mod3)
sp_mod3_resid <- simulateResiduals(sp_mod3)
plot(sp_mod3_resid)#good

sp_mod3b<-glmmTMB(AnySpiral ~ Spectrum+Intensity+CaptureMethod+
                    (1|LatinName)+(1|Family),
                  family=binomial, data=EXCLCONTROL_EXPT1)
anova(sp_mod3, sp_mod3b)#remove interaction
summary(sp_mod3b)

sp_mod3c<-glmmTMB(AnySpiral ~ Intensity+Spectrum+
                    (1|LatinName)+(1|Family),
                  family=binomial, data=EXCLCONTROL_EXPT1)
anova(sp_mod3c, sp_mod3b)#remove capture method
summary(sp_mod3c)
sp_mod3d<-glmmTMB(AnySpiral ~ Intensity+
                    (1|LatinName)+(1|Family),
                  family=binomial, data=EXCLCONTROL_EXPT1)
anova(sp_mod3d, sp_mod3c)#remove spectrum
summary(sp_mod3d)
sp_mod3e<-glmmTMB(AnySpiral ~ 1+
                    (1|LatinName)+(1|Family),
                  family=binomial, data=EXCLCONTROL_EXPT1)
anova(sp_mod3d, sp_mod3e)#significant effect of intensity only
summary(glht(sp_mod3d, linfct = mcp(Intensity = "Tukey")))
#30 lx significantly different from both 0.3 and 3 lx treatments

plot_model(sp_mod3d, type = "pred", 
                                    terms = c("Intensity"),
                                    colors="gs",
                                    title="")+
  theme_classic()+
  xlab("Light intensity (lx)")+ylab("Predicted probability of observing spirals (%)")+
  theme(axis.title.x = element_text(size = 12),
        axis.title=element_text(size=16),
        axis.text.x = element_text(size = 10),
        axis.title.y = element_text(size = 12),
        axis.text.y=element_text(size=10))

#7)Trajectory analyses----

#functions----

# Function to compute turning angles, vector lengths, and bearing to light in 3D
calculate_turning_angles_and_lengths <- function(df) {
  
  coords <- as.matrix(df[, c("X_smooth", "Y_smooth", "Z_smooth")])
  
  # Step 1: Compute direction vectors (differences between consecutive points)
  vectors <- diff(coords)
  
  # Step 2: Function to compute 3D angle between two vectors
  angle_between <- function(v1, v2) {
    dot_prod <- sum(v1 * v2)
    norm_v1 <- sqrt(sum(v1^2))
    norm_v2 <- sqrt(sum(v2^2))
    cos_theta <- dot_prod / (norm_v1 * norm_v2)
    cos_theta <- max(min(cos_theta, 1), -1)  # Clamp to avoid rounding errors
    return(acos(cos_theta))  # Angle in radians
  }
  
  # Step 3: Compute turning angles between consecutive flight vectors
  angles <- numeric(nrow(vectors) - 1)
  for (i in 1:length(angles)) {
    angles[i] <- angle_between(vectors[i, ], vectors[i + 1, ])
  }
  
  # Step 4: Calculate vector lengths 
  vector_lengths <- sqrt(rowSums(vectors^2))
  
  # Step 5: Compute distance between first and last points
  first_point <- coords[1, ]
  last_point <- coords[nrow(coords), ]
  distance_first_last <- sqrt(sum((last_point - first_point)^2))
  
  # Step 6: Compute distances to the focal light (or central focal light, if multiple)
  focallight <- c(df$LightX3D[1], df$LightY3D[1], df$LightZ3D[1])
  distance_to_light <- sqrt(rowSums((coords - matrix(focallight, nrow(coords), 3, byrow = TRUE))^2))
  
  # Step 7: Compute bearing to light for each vector
  # For each segment, compare the flight vector to the vector pointing to the light from that segment's start point
  bearing_to_light <- numeric(nrow(vectors))
  for (i in 1:nrow(vectors)) {
    flight_vec <- vectors[i, ]
    to_light_vec <- focallight - coords[i, ]  # vector from current point to light
    bearing_to_light[i] <- angle_between(flight_vec, to_light_vec)
  }
  
  # Step 8: Add results to dataframe
  df$turning_angle_rad <- c(NA, NA, angles)
  df$turning_angle_deg <- c(NA, NA, angles * 180 / pi)
  df$vector_length <- c(NA, vector_lengths)
  df$distance_first_last <- distance_first_last
  df$distance_to_light <- distance_to_light
  df$bearing_to_light_rad <- c(NA, bearing_to_light)
  df$bearing_to_light_deg <- c(NA, bearing_to_light * 180 / pi)
  
  return(df)
}


# Function to calculate metrics
calculate_tortuosity_metrics <- function(df) {
  # Calculate total path length
  total_path_length <- sum(df$vector_length, na.rm = TRUE)
  
  # Calculate number of data points
  total_path_freq <- nrow(df)
  total_path_duration<-total_path_freq/15 # duration of flight in s,with videos at 15 fps
  
  # Calculate speed - meters per second
  speed_ms <- total_path_length / total_path_duration
  
  # Calculate mean cosine of turning angles
  mean_cos_angle <- mean(cos(df$turning_angle_rad), na.rm = TRUE)
  
  # Calculate mean of turning angles
  mean_angle <- mean(df$turning_angle_deg, na.rm = TRUE)
  
  # Calculate sd of turning angles
  sd_angle <- sd(df$turning_angle_deg, na.rm = TRUE)
  
  # Calculate sinuosity
  sinuosity <- 1/((2 * total_path_length) / (1 - mean_cos_angle))
  
  # AB length of track
  distance_first_last <- mean(df$distance_first_last)
  
  # Mean distance to light
  mean_distance_to_light <- mean(df$distance_to_light)
  
  #Return a new dataframe with the metrics
  
  #take first row of each df (by unique moth or flight id)
  #and remove uninformative rows
  newdf<-df[1,]%>%
    dplyr::select(!c(A_X,A_Y,A_Centre_light_X,A_Centre_light_Y,
                    B_X,B_Y, B_Centre_light_X,B_Centre_light_Y,
                    Moth_ID, trim,Displacement,turning_angle_rad,turning_angle_deg,
                    vector_length,distance_to_light,bearing_to_light_rad, bearing_to_light_deg))
  #add metrics calculated above
  newdf$total_path_length[1] = total_path_length
  newdf$total_path_freq[1] = total_path_freq
  newdf$speed_ms_3D[1] = speed_ms
  newdf$distance_first_last[1] = distance_first_last
  newdf$mean_angle_3D [1]= mean_angle
  newdf$sd_angle_3D[1] = sd_angle
  newdf$mean_cos_angle[1] = mean_cos_angle
  newdf$sinuosity[1] = sinuosity
  newdf$mean_distance_to_light[1] = mean_distance_to_light
  newdf$track_duration[1] =total_path_duration

  return(newdf)
}

#calculate tortuosity and other flight metrics----

#for 3D flights that meet analysis criteria only
#apply turning angle function to each unique flight (N=270) 
FlightIDs<-unique(SINGLETRACKS_LIGHT$Unique_Moth_ID)
FLIGHTS<-list()

for (i in 1:length(FlightIDs)){
  TEMPDF<-SINGLETRACKS_LIGHT%>%
    filter(Unique_Moth_ID ==FlightIDs[i])
  str(TEMPDF)
  FLIGHTS[[i]]<-calculate_turning_angles_and_lengths(TEMPDF)#create new list with data with turning angles for each flight
}

#before calculating summary metrics for each track, remove points identified as having large displacements in 3D,
#indicating error in 3D tracking
FLIGHTS<-lapply(FLIGHTS, function(x) filter(x, Displacement < 0.5))

#combine into larger dataframe
SINGLETRACKS_ANGLES <- bind_rows(FLIGHTS)

#check if NAs are correct
length(which(is.na(SINGLETRACKS_ANGLES$bearing_to_light_deg)))# 270 NAs -one per flight, in first row

#apply tortuosity function to each flight
SINGLE_TORTUOSITY_LIST <- lapply(FLIGHTS, calculate_tortuosity_metrics)
length(SINGLE_TORTUOSITY_LIST)
head(SINGLE_TORTUOSITY_LIST[[1]])

#combine tortuosity results
SINGLETRACKS_METRICS <- do.call(rbind,SINGLE_TORTUOSITY_LIST)
str(SINGLETRACKS_METRICS)

#plot results
col_names <- c("Unique_Moth_ID","Treatment","LatinName","Family","CaptureMethod","Location")
SINGLETRACKS_METRICS <- SINGLETRACKS_METRICS %>%
  mutate(across(all_of(col_names), as.factor))
str(SINGLETRACKS_METRICS)
levels(SINGLETRACKS_METRICS$Treatment)

#simple plot of mean turning angle by treatment
plot_tortuosity <- ggplot(SINGLETRACKS_METRICS, aes(x = Treatment, y = mean_angle_3D, fill = Treatment)) +
  geom_boxplot() +
  theme_classic() +
  labs(title = "", y = "Mean turning angle (degrees)", x = NULL) +
  theme(
    axis.text = element_text(size = 16),
    axis.title = element_text(size = 18),
    axis.text.x = element_text(angle = 45, hjust = 1),
    legend.position = "none",
    panel.grid.major = element_blank(),
    panel.grid.minor = element_blank(),
    panel.border = element_blank(),  # keep axis lines, so we don't add a full box
    axis.line = element_line(color = "black")  # this adds back the x and y axis lines
  )
plot_tortuosity

#and standard deviation of turning angle
plot_SD <- ggplot(SINGLETRACKS_METRICS, aes(x = Treatment, y = sd_angle_3D, fill = Treatment)) +
  geom_boxplot() +
  theme_classic() +  # Clean theme
  labs(title = "", y = "SD turning angle") +  # Remove x axis label
  theme(
    axis.text = element_text(size = 16),
    axis.title = element_text(size = 18),
    axis.text.x = element_text(angle = 45, hjust = 1),
    legend.position = "none",
    panel.grid.major = element_blank(),
    panel.grid.minor = element_blank(),
    panel.border = element_blank(),  # keep axis lines, so we don't add a full box
    axis.line = element_line(color = "black"))
plot_SD

#statistics on overall metrics----

#dataset for comparison between years:
YEAR_SINGLETRACKS_METRICS<-SINGLETRACKS_METRICS%>%
  filter(Treatment=="Control"|Treatment=="BW_30lx"|Treatment=="BA_30lx"|Treatment=="NA_30lx")%>%
  droplevels()
#dataset for comparison between effects of spectrum and intensity:
SINGLETRACKS_METRICS_EXCLCONTROL<-SINGLETRACKS_METRICS%>%
  filter(Treatment!="Control")%>%
  droplevels()

#1) mean3D turning angle----

#check if year is important
year_tort1<-lmer(mean_angle_3D ~ Treatment*Year+CaptureMethod+(1|Family) + (1|LatinName), data=YEAR_SINGLETRACKS_METRICS)
year_tort1_resid <- simulateResiduals(year_tort1)
plot(year_tort1_resid)#good
summary(year_tort1)

year_tort1b<-lmer(mean_angle_3D ~ Treatment+Year+CaptureMethod+(1|Family) + (1|LatinName), data=YEAR_SINGLETRACKS_METRICS)
anova(year_tort1, year_tort1b)#can remove interaction
year_tort1c<-lmer(mean_angle_3D ~ Treatment+CaptureMethod+(1|Family) + (1|LatinName), data=YEAR_SINGLETRACKS_METRICS)
anova(year_tort1c, year_tort1b)#can remove year and move on to analyse both years together

#across both years
tort_mod1<-lmer(mean_angle_3D ~ Treatment*CaptureMethod+(1|Family) + (1|LatinName), data=SINGLETRACKS_METRICS)
tort1_resid <- simulateResiduals(tort_mod1)
plot(tort1_resid)#poor
hist(resid(tort_mod1))
tort_mod1<-lmer(sqrt(mean_angle_3D) ~ Treatment*CaptureMethod+(1|Family) + (1|LatinName), data=SINGLETRACKS_METRICS)
tort1_resid <- simulateResiduals(tort_mod1)
plot(tort1_resid)#better
summary(tort_mod1)

tort_mod1b<-lmer(sqrt(mean_angle_3D) ~ Treatment+CaptureMethod+(1|Family) + (1|LatinName), data=SINGLETRACKS_METRICS)
anova(tort_mod1, tort_mod1b)
summary(tort_mod1b)
tort_mod1c<-lmer(sqrt(mean_angle_3D) ~ Treatment+(1|Family) + (1|LatinName), data=SINGLETRACKS_METRICS)
anova(tort_mod1c, tort_mod1b)
summary(tort_mod1c)
tort_mod1d<-lmer(sqrt(mean_angle_3D) ~ 1+(1|Family) + (1|LatinName), data=SINGLETRACKS_METRICS)
anova(tort_mod1c, tort_mod1d)
summary(tort_mod1c)

summary(glht(tort_mod1c, linfct = mcp(Treatment = "Tukey"))) #all sig differences involve 30lx treatments

#spectrum and intensity
tort_mod2<-lmer(mean_angle_3D ~ Spectrum*Intensity+CaptureMethod+(1|Family) + (1|LatinName), data=SINGLETRACKS_METRICS_EXCLCONTROL)
tort2_resid <- simulateResiduals(tort_mod2)
plot(tort2_resid)#poor
hist(resid(tort_mod2))
tort_mod2<-lmer(sqrt(mean_angle_3D) ~ Spectrum*Intensity+CaptureMethod+(1|Family) + (1|LatinName), data=SINGLETRACKS_METRICS_EXCLCONTROL)
tort2_resid <- simulateResiduals(tort_mod2)
plot(tort2_resid)#better
hist(resid(tort_mod2))

summary(tort_mod2)
tort_mod2b<-lmer(sqrt(mean_angle_3D) ~ Spectrum+Intensity+CaptureMethod+(1|Family) + (1|LatinName), data=SINGLETRACKS_METRICS_EXCLCONTROL)
anova(tort_mod2, tort_mod2b)
summary(tort_mod2b)
tort_mod2c<-lmer(sqrt(mean_angle_3D) ~ Intensity+CaptureMethod+(1|Family) + (1|LatinName), data=SINGLETRACKS_METRICS_EXCLCONTROL)
anova(tort_mod2c, tort_mod2b)
summary(tort_mod2c)
tort_mod2d<-lmer(sqrt(mean_angle_3D) ~ Intensity+(1|Family) + (1|LatinName), data=SINGLETRACKS_METRICS_EXCLCONTROL)
anova(tort_mod2c, tort_mod2d)
summary(tort_mod2d)
tort_mod2e<-lmer(sqrt(mean_angle_3D) ~ 1+(1|Family) + (1|LatinName), data=SINGLETRACKS_METRICS_EXCLCONTROL)
anova(tort_mod2d, tort_mod2e)#significant effect of intensity

summary(glht(tort_mod2d, linfct = mcp(Intensity = "Tukey")))#30lx different from others

#2) sd of turning angle----

#check if year is important
year_tort2<-lmer(sd_angle_3D ~ Treatment*Year+CaptureMethod+(1|Family) + (1|LatinName), data=YEAR_SINGLETRACKS_METRICS)
year_tort2_resid <- simulateResiduals(year_tort2)
plot(year_tort2_resid)#ok
summary(year_tort2)

year_tort2b<-lmer(sd_angle_3D ~ Treatment+Year+CaptureMethod+(1|Family) + (1|LatinName), data=YEAR_SINGLETRACKS_METRICS)
anova(year_tort2, year_tort2b)#can remove interaction
year_tort2c<-lmer(sd_angle_3D ~ Treatment+CaptureMethod+(1|Family) + (1|LatinName), data=YEAR_SINGLETRACKS_METRICS)
anova(year_tort2c, year_tort2b)#can remove year and move on to analyse both years together

#test effect of treatment across both years
tort_mod3<-lmer(sd_angle_3D ~ Treatment*CaptureMethod+(1|Family) + (1|LatinName), data=SINGLETRACKS_METRICS)
tort3_resid <- simulateResiduals(tort_mod3)
plot(tort3_resid)#ok

tort_mod3b<-lmer(sd_angle_3D ~ Treatment+CaptureMethod+(1|Family)+ (1|LatinName) , data=SINGLETRACKS_METRICS)
anova(tort_mod3, tort_mod3b)
summary(tort_mod3b)
tort_mod3c<-lmer(sd_angle_3D ~ Treatment+(1|Family) + (1|LatinName), data=SINGLETRACKS_METRICS)
anova(tort_mod3c, tort_mod3b)
summary(tort_mod3c)#can remove capture method
tort_mod3d<-lmer(sd_angle_3D ~ 1+(1|Family)+ (1|LatinName) , data=SINGLETRACKS_METRICS)
anova(tort_mod3c, tort_mod3d) #effect of treatment

summary(glht(tort_mod3c, linfct = mcp(Treatment = "Tukey")))
  #all significant contrasts involve 30 lx lights vs control or 0.3 lx

#spectrum and intensity
tort_mod4<-lmer(sd_angle_3D ~ Spectrum*Intensity+CaptureMethod+(1|Family)+ (1|LatinName), data=SINGLETRACKS_METRICS_EXCLCONTROL)
tort4_resid <- simulateResiduals(tort_mod4)
plot(tort4_resid)#ok
summary(tort_mod4)

tort_mod4b<-lmer(sd_angle_3D ~ Spectrum+Intensity+CaptureMethod+(1|Family)+ (1|LatinName), data=SINGLETRACKS_METRICS_EXCLCONTROL)
anova(tort_mod4, tort_mod4b)#can just remove interaction
summary(tort_mod4b)
tort_mod4c<-lmer(sd_angle_3D ~ Intensity+CaptureMethod+(1|Family)+ (1|LatinName) , data=SINGLETRACKS_METRICS_EXCLCONTROL)
anova(tort_mod4c, tort_mod4b)#remove spectrum
summary(tort_mod4c)
tort_mod4d<-lmer(sd_angle_3D ~ Intensity+(1|Family)+ (1|LatinName), data=SINGLETRACKS_METRICS_EXCLCONTROL)
anova(tort_mod4c, tort_mod4d)#remove capture method
summary(tort_mod4d)
tort_mod4e<-lmer(sd_angle_3D ~ 1+(1|Family) + (1|LatinName), data=SINGLETRACKS_METRICS_EXCLCONTROL)
anova(tort_mod4d, tort_mod4e)#significant effect of intensity

summary(glht(tort_mod4d, linfct = mcp(Intensity = "Tukey")))#30lx different from others

#flight metrics before and after exposure to light----
  #exposure to light means that the moth has passed into the area lit by the focal light beam (or the equivalent area, in the control)
  #and was able to perceive it (ie. not heading straight away from it)

#create variable to signify whether or not the moth is able to see the light, based on bearing angle (0 no and 1 yes) 
SINGLETRACKS_ANGLES$Exposed <- ifelse(SINGLETRACKS_ANGLES$bearing_to_light_deg <= 135, 1, 0)

#combine with position after light cone to create variable indicating when moth has both passed the light and  "seen" it
SINGLETRACKS_ANGLES$After_Light_Both_Exposed <- ifelse(
  !is.na(SINGLETRACKS_ANGLES$Exposed) & 
    !is.na(SINGLETRACKS_ANGLES$After_Light_Both) & 
    SINGLETRACKS_ANGLES$Exposed == 1 & SINGLETRACKS_ANGLES$After_Light_Both == 1,
  1,
  0
)

#create variable to separate flights before and after exposure
SINGLETRACKS_ANGLES$UniqueSegment_AfterLightBoth<-paste(SINGLETRACKS_ANGLES$Unique_Moth_ID,SINGLETRACKS_ANGLES$After_Light_Both, sep="_")
SINGLETRACKS_ANGLES$UniqueSegment<-paste(SINGLETRACKS_ANGLES$Unique_Moth_ID,SINGLETRACKS_ANGLES$After_Light_Both_Exposed, sep="_")
str(SINGLETRACKS_ANGLES)

#identify flights with at least 5 points before and after cone (in both cameras)
WITHBOTH<-SINGLETRACKS_ANGLES%>%
  group_by(Unique_Moth_ID,After_Light_Both_Exposed)%>%
  summarise(n=n())%>%
  filter(n>=5)%>%
  group_by(Unique_Moth_ID)%>%
  summarise(n=n())%>%
  filter(n==2)
unique(WITHBOTH$Unique_Moth_ID)#255 flights out of 270

#restrict track data to only those flights
TRACKS_BEFOREAFTER<-SINGLETRACKS_ANGLES%>%
  filter(Unique_Moth_ID %in% WITHBOTH$Unique_Moth_ID)

#apply function to calculate tortuosity metrics for these segments

#split by unique segment IDs - turning angles etc already calculated
segmentIDs<-unique(TRACKS_BEFOREAFTER$UniqueSegment)
SEGMENTS<-list()

for (i in 1:length(segmentIDs)){
  SEGMENTS[[i]]<-TRACKS_BEFOREAFTER%>%
    filter(UniqueSegment == segmentIDs[i])
}

# Apply tortuosity function to each dataframe in the results_list
SINGLE_TORTUOSITY_LIST_SEGMENTS <- lapply(SEGMENTS, calculate_tortuosity_metrics)
length(SINGLE_TORTUOSITY_LIST_SEGMENTS)#510 - 2 segments each for 255 flights
head(SINGLE_TORTUOSITY_LIST_SEGMENTS[[1]])

# Combine the results into a single dataframe
SINGLETRACKS_SEGMENTS <- do.call(rbind, SINGLE_TORTUOSITY_LIST_SEGMENTS)
dim(SINGLETRACKS_SEGMENTS)

#1) mean turning angle----

col_names2 <- c("Unique_Moth_ID","UniqueSegment","After_Light_Both_Exposed","Treatment","LatinName","Family","CaptureMethod","Location")
SINGLETRACKS_SEGMENTS<- SINGLETRACKS_SEGMENTS %>%
  mutate(across(all_of(col_names2), as.factor))
str(SINGLETRACKS_SEGMENTS)
dim(SINGLETRACKS_SEGMENTS)

#number of flight paths per treatment:
with(SINGLETRACKS_SEGMENTS,table(Treatment,After_Light_Both_Exposed))

#model impact of exposure to light and treatment on mean turning angles
ba_tort_mod1 <- lmer(mean_angle_3D ~ Treatment*After_Light_Both_Exposed +CaptureMethod+ (1|Family) + (1|LatinName) + (1|Unique_Moth_ID), data=SINGLETRACKS_SEGMENTS)
resid_ba_tort_mod1<-simulateResiduals(ba_tort_mod1)
plot(resid_ba_tort_mod1)#poor
plot(ba_tort_mod1)
hist(resid(ba_tort_mod1))
ba_tort_mod1 <- lmer(sqrt(mean_angle_3D) ~ Treatment*After_Light_Both_Exposed +CaptureMethod+ (1|Family) + (1|LatinName)+ (1|Unique_Moth_ID), data=SINGLETRACKS_SEGMENTS)
resid_ba_tort_mod1<-simulateResiduals(ba_tort_mod1)
plot(resid_ba_tort_mod1)
plot(ba_tort_mod1)#looks ok
hist(resid(ba_tort_mod1))#better
summary(ba_tort_mod1)
anova(ba_tort_mod1, type = 3)  # type-III ANOVA tests interaction and main effects - found significant interaction
ba_tort_mod1b <- lmer(sqrt(mean_angle_3D) ~ Treatment+After_Light_Both_Exposed +CaptureMethod+ (1|Family) + (1|LatinName) + (1|Unique_Moth_ID), data=SINGLETRACKS_SEGMENTS)
anova(ba_tort_mod1, ba_tort_mod1b)#significant effect of interaction
ba_tort_mod1c <- lmer(sqrt(mean_angle_3D) ~ Treatment*After_Light_Both_Exposed+ (1|Family) + (1|LatinName) + (1|Unique_Moth_ID), data=SINGLETRACKS_SEGMENTS)
anova(ba_tort_mod1, ba_tort_mod1c)#no significant effect of capture method
ba_tort_mod1d <- lmer(sqrt(mean_angle_3D) ~ Treatment+After_Light_Both_Exposed+ (1|Family) + (1|LatinName) + (1|Unique_Moth_ID), data=SINGLETRACKS_SEGMENTS)
anova(ba_tort_mod1d, ba_tort_mod1c)#still a significant effect of interaction
summary(ba_tort_mod1c)

#effect of exposure for every treatment level - see: https://cran.r-project.org/web/packages/emmeans/vignettes/comparisons.html
emmres<-emmeans(ba_tort_mod1c, ~ After_Light_Both_Exposed | Treatment)
pairs(emmres)
#additional figure with pairwise comparisons: (plotted 700x622h)
plot(emmres, comparisons=TRUE)+
  xlab("Estimated marginal means")+ylab("Exposure to light (0 = before exposure, 1 = after exposure)")+theme_bw()

#plot tortuosity before and after exposure - figure 4D

#create new variable for plotting
SINGLETRACKS_SEGMENTS$TreatmentSeg<-paste(SINGLETRACKS_SEGMENTS$Treatment,SINGLETRACKS_SEGMENTS$After_Light_Both_Exposed)
SINGLETRACKS_SEGMENTS$TreatmentSeg<-factor(SINGLETRACKS_SEGMENTS$TreatmentSeg,
                                           levels = c("Control 0",  "Control 1",
                                                      "BW_0.3lx 0", "BW_0.3lx 1",
                                                      "BW_3lx 0" ,  "BW_3lx 1",
                                                      "BW_30lx 0",  "BW_30lx 1",
                                                      "BA_0.3lx 0", "BA_0.3lx 1",
                                                      "BA_3lx 0","BA_3lx 1",
                                                      "BA_30lx 0" , "BA_30lx 1",
                                                      "NA_0.3lx 0","NA_0.3lx 1",
                                                      "NA_3lx 0" ,  "NA_3lx 1",
                                                      "NA_30lx 0" , "NA_30lx 1"))
levels(SINGLETRACKS_SEGMENTS$TreatmentSeg)
expt1colours<-c("#ab7fff","#afddfa","#64c1fa","#19a4fa","#ffe4b2",
                "#ffc966","#ffae1a","#ffc7b2","#ff8f66","#ff571a")
expt1colours2<-c("white","#ab7fff","white","#afddfa","white","#64c1fa","white","#19a4fa","white","#ffe4b2",
                 "white","#ffc966","white","#ffae1a","white","#ffc7b2","white","#ff8f66","white","#ff571a")
expt1colours3<-c("#ab7fff","grey40","#afddfa","grey40","#64c1fa","grey40","#19a4fa","grey40","#ffe4b2",
                 "grey40","#ffc966","grey40","#ffae1a","grey40","#ffc7b2","grey40","#ff8f66","grey40","#ff571a","grey40")


plot_tortuosity_ba <- ggplot(SINGLETRACKS_SEGMENTS, aes(x = Treatment, 
                                                               y = mean_angle_3D, col = TreatmentSeg, fill = TreatmentSeg)) +
  geom_boxplot() +
  theme_classic() +
  scale_fill_manual(values=expt1colours2)+
  scale_color_manual(values=expt1colours3)+
  guides(col="none")+guides(fill="none")+
  labs(title = "D", y = "Mean turning angle (degrees)", x = NULL, fill="") +
  annotate(
    "text",
    x = c(1, 2, 3, 4, 5, 6,7,8,9,10),     # numeric positions treatments on the x-axis
    y = c(75, 75, 75, 75, 75,75,75,75,75,75),  # y positions for asterisks
    label = c("", "*", "***","***", "", "*","***", "", "","***"),  # asterisks corresponding to pvalues from emmeans
    size = 4 ) +
  labs(fill = "Flight \nsegment")+  scale_x_discrete(labels = c("Control","White 0.3 lx", "White 3 lx", "White 30 lx",
                                                                "BA 0.3 lx", "BA 3 lx", "BA 30 lx",
                                                                "NA 0.3 lx", "NA 3 lx", "NA 30 lx"))+
  
  theme(
    axis.text = element_text(size = 12),
    axis.title = element_text(size = 16),
    axis.text.x = element_text(angle = 45, hjust = 1),
    panel.grid.major = element_blank(),
    panel.grid.minor = element_blank(),
    panel.border = element_blank(),  
    axis.line = element_line(color = "black") 
  )

plot_tortuosity_ba

#2) sd turning angle ----
ba_sd_mod1 <- lmer(sd_angle_3D ~ Treatment*After_Light_Both_Exposed +CaptureMethod+ (1|Family) +
                     (1|Unique_Moth_ID)+(1|LatinName) , data=SINGLETRACKS_SEGMENTS)
  #singular fit, try removing a random effect
resid_ba_sd_mod1<-simulateResiduals(ba_sd_mod1)
plot(resid_ba_sd_mod1)
plot(ba_sd_mod1)
hist(resid(ba_sd_mod1))
ba_sd_mod2 <- lmer(log(sd_angle_3D) ~ Treatment*After_Light_Both_Exposed +CaptureMethod+ (1|LatinName) +
                     (1|Unique_Moth_ID), data=SINGLETRACKS_SEGMENTS)
ba_sd_mod3 <- lmer(log(sd_angle_3D) ~ Treatment*After_Light_Both_Exposed +CaptureMethod+ (1|Family) +
                     (1|Unique_Moth_ID), data=SINGLETRACKS_SEGMENTS)
AIC(ba_sd_mod1, ba_sd_mod2, ba_sd_mod3)#best model includes family and unique moth id
resid_ba_sd_mod3<-simulateResiduals(ba_sd_mod3)
plot(resid_ba_sd_mod3)#poor

#try transforms
ba_sd_mod1 <- lmer(sqrt(sd_angle_3D) ~ Treatment*After_Light_Both_Exposed +CaptureMethod+ (1|Family) +
(1|Unique_Moth_ID)+(1|LatinName) , data=SINGLETRACKS_SEGMENTS)

ba_sd_mod2 <- lmer(sqrt(sd_angle_3D) ~ Treatment*After_Light_Both_Exposed +CaptureMethod+ (1|LatinName) +
                     (1|Unique_Moth_ID), data=SINGLETRACKS_SEGMENTS)
ba_sd_mod3 <- lmer(sqrt(sd_angle_3D) ~ Treatment*After_Light_Both_Exposed +CaptureMethod+ (1|Family) +
                     (1|Unique_Moth_ID), data=SINGLETRACKS_SEGMENTS)
AIC(ba_sd_mod1, ba_sd_mod2, ba_sd_mod3)#best model still includes family and unique moth id
resid_ba_sd_mod3<-simulateResiduals(ba_sd_mod3)
plot(resid_ba_sd_mod3)#better

ba_sd_mod3b <- lmer(sqrt(sd_angle_3D) ~ Treatment+After_Light_Both_Exposed +CaptureMethod+ (1|Unique_Moth_ID)+(1|Family), data=SINGLETRACKS_SEGMENTS)
anova(ba_sd_mod3, ba_sd_mod3b)#significant effect of interaction
summary(ba_sd_mod3b)
ba_sd_mod3c <- lmer(sqrt(sd_angle_3D) ~ Treatment*After_Light_Both_Exposed+ (1|Family) + (1|Unique_Moth_ID) , data=SINGLETRACKS_SEGMENTS)
anova(ba_sd_mod3, ba_sd_mod3c)#no significant effect of capture method
summary(ba_sd_mod3c)
ba_sd_mod3d <- lmer(sqrt(sd_angle_3D) ~ Treatment+After_Light_Both_Exposed+ (1|Family) + (1|Unique_Moth_ID) , data=SINGLETRACKS_SEGMENTS)
anova(ba_sd_mod3d, ba_sd_mod3c)#still significant effect of interaction

emmeans(ba_sd_mod3c,  pairwise ~ After_Light_Both_Exposed | Treatment) #p values per treatment group

plot_sd_ba <- ggplot(SINGLETRACKS_SEGMENTS, aes(x = Treatment, 
                                                               y = log(sd_angle_3D), col = TreatmentSeg, fill = TreatmentSeg)) +
  geom_boxplot() +
  theme_classic() +
  scale_fill_manual(values=expt1colours2)+
  scale_color_manual(values=expt1colours3)+
  guides(col="none")+guides(fill="none")+
  labs(title = "A", y = "sqrt (SD turning angle (degrees))", x = "Focal light treatment", fill="") +
  annotate(
    "text",
    x = c(1, 2, 3, 4, 5, 6,7,8,9,10),     
    y = c(5, 5, 5, 5, 5,5,5,5,5,5),  
    label = c("", "*", "***","*", "", "","**", "", "",""),
    size = 4 ) +
  labs(fill = "Flight \nsegment")+  scale_x_discrete(labels = c("Control","White 0.3 lx", "White 3 lx", "White 30 lx",
                                                                "BA 0.3 lx", "BA 3 lx", "BA 30 lx",
                                                                "NA 0.3 lx", "NA 3 lx", "NA 30 lx"))+
  
  theme(
    axis.text = element_text(size = 12),
    axis.title = element_text(size = 16),
    axis.text.x = element_text(angle = 45, hjust = 1),
    panel.grid.major = element_blank(),
    panel.grid.minor = element_blank(),
    panel.border = element_blank(),  
    axis.line = element_line(color = "black") 
  )
plot_sd_ba


#EXPERIMENT 2 - MULTIPLE LIGHTS----

#Experiment 2 - trial information----

INFO_MULTI<-read.csv("Experiment2_MothInfo.csv", header=TRUE)
str(INFO_MULTI) #900 flights

#identify which moths only have a single flight
EXCEPTIONS1<-INFO_MULTI%>%
  group_by(Unique_Moth_ID)%>%
  summarise(n=n())%>%
  filter(n==1)
dim(EXCEPTIONS1)
  #24 moths have only one flight - 
  #they are all moths exposed to a central light in 3lx background as trial 2
  #their trial 1 was the 3lx background control (no focal light)
  #providing supplementary data for experiment 1
  #of these 24 moths, only 2 took off

#this leaves 876 paired flights (438 moths)

#break down treatment into test light structure and background light levels
for (i in 1:length(INFO_MULTI$Treatment)){
  INFO_MULTI$LightStructure[i]<-str_split(INFO_MULTI$Treatment, "_")[[i]][1]
  INFO_MULTI$BgdLightLevel[i]<-str_split(INFO_MULTI$Treatment, "_")[[i]][2]
}
#recode take-off yes/no and light treatment variables, and set variables to factors where needed
factors <- c("Unique_Flight_ID","Unique_Moth_ID","MothID", "Treatment","LatinName","Family",
             "CaptureMethod","Location","TrialNumber","Year","LightStructure", "BgdLightLevel")
INFO_MULTI<-INFO_MULTI%>%
  mutate(LightStructure=dplyr::recode(LightStructure,
                               "DL"="Multiple","CL"="Single","SingleC" ="Single 10lx"))%>%
  mutate(BgdLightLevel=dplyr::recode(BgdLightLevel,
                              "0.03lx"="Low","0.3lx"="Medium","3lx"="High",))%>%
  mutate(across(all_of(factors), as.factor))%>%
  mutate(BgdLightLevel = factor(BgdLightLevel,
                            levels = c("Low","Medium","High")))%>%
  mutate(LightStructure = factor(LightStructure,
                                levels = c("Single","Multiple","Single 10lx")))%>%
  mutate(Treatment = factor(Treatment,
                                 levels = c("CL_0.03lx","DL_0.03lx","CL_0.3lx","DL_0.3lx","CL_3lx","DL_3lx","SingleC_0.3lx")))
str(INFO_MULTI)

#remove additional "single light 10 lx" treatment from main dataset
#and create supplementary dataset with only medium background light level in 2024
SUPP_MULTI<-INFO_MULTI%>%
  filter(BgdLightLevel=="Medium")%>%
  filter(Year=="2024")%>%
  droplevels()
str(SUPP_MULTI)#260 flights
#View((SUPP_MULTI))
INFO_MULTI<-INFO_MULTI%>%
  filter(LightStructure!="Single 10lx")%>%
  droplevels()
dim(INFO_MULTI)#now 853 flights - 47 flights in single 10 lx group

#identify which moths now only have a single flight due to removal of single 10lx flights
INFO_SINGLE10LX<-SUPP_MULTI%>%
  filter(LightStructure=="Single 10lx")%>%
  droplevels()
dim(INFO_SINGLE10LX)
EXCEPTIONS2<-INFO_MULTI%>%
  filter(Unique_Moth_ID %in% INFO_SINGLE10LX$Unique_Moth_ID)
#View(EXCEPTIONS2)
  #all have been paired with the multiple light treatment, background light level of 0.3 lx
  #of the 47 trials, moths took off in 18
  #806 paired flights remain

#count number of species & families in experiment 2
Expt2Individuals<-length(levels(INFO_MULTI$Unique_Moth_ID))
Expt2Species<-unique(INFO_MULTI$LatinName)
Expt2Family<-unique(INFO_MULTI$Family)
Expt2Individuals#462 individuals
length(Expt2Species)#32 species
length(Expt2Family) #5 families tested

#sample sizes per treatment and year:
INFO_MULTI%>%
  group_by(Treatment, Year)%>%
  summarise(n=n())#samples range from 46 to 164 trials per group
MULTISPEC<-INFO_MULTI%>%
  group_by(Unique_Moth_ID, LatinName)%>%
  summarise(n=n())%>%
  group_by(LatinName)%>%
  summarise(n=n())
#number of species tested across both experiments:
length(unique(c(Expt1Species,Expt2Species)))#62 species 
length(unique(c(Expt1Family,Expt2Family)))#from 6 families

#Experiment 2 - track data ----

MULTITRACKS<-read.csv("Experiment2_AnalysedTracks.csv", header=TRUE)
head(MULTITRACKS)
str(MULTITRACKS)
unique(MULTITRACKS$Unique_Flight_ID)#302 tracks
unique(MULTITRACKS$Unique_Moth_ID)#from 219 individual moths

#Clean data

#1) remove tracks that do not cross light path

#identify flights that do not contain points in light cone for both cameras
NOLIGHT_EXPT2<-MULTITRACKS%>%
  group_by(Unique_Flight_ID)%>%
  summarise(PointsInLight=sum(After_Light_Both))%>%
  filter(PointsInLight==0)
unique(NOLIGHT_EXPT2$Unique_Flight_ID)#23 flights to remove

#remove those unique flights from dataset
MULTITRACKS_LIGHT<-MULTITRACKS%>%
  filter(!(Unique_Flight_ID %in% NOLIGHT_EXPT2$Unique_Flight_ID))

#2)check that tracks start and end according to rules

MULTITRACKS_LIGHT_CHECK_A_B <- MULTITRACKS_LIGHT %>%
  group_by(Unique_Flight_ID) %>% #group by track
  arrange(Frame, by_group="Unique_Flight_ID") %>% #arrange frames in order (this should already be the case)
  mutate(Euclidean_distance_A = sqrt((A_X-lag(A_X))^2+ (A_Y-lag(A_Y))^2)) %>% #calculate euclidean distance between consecutive points in the track from camera A
  mutate(Stop_A = ifelse(Euclidean_distance_A==0, 1, 0)) %>% #identify points where moth does not move
  mutate(Counter_A = Stop_A + lag(Stop_A,1)) %>% #look for two instances of 0 movement (this implies 3 consecutive points in same position)
  mutate(Euclidean_distance_B = sqrt((B_X-lag(B_X))^2+ (B_Y-lag(B_Y))^2)) %>% #repeat for track from camera B
  mutate(Stop_B = ifelse(Euclidean_distance_B==0, 1, 0)) %>%
  mutate(Counter_B = Stop_B + lag(Stop_B,1)) %>%
  filter(any(Counter_A>=2|Counter_B>=2)) %>% #restrict to cases where conditions are met in tracks from A or B
  ungroup()
unique(MULTITRACKS_LIGHT_CHECK_A_B$Unique_Flight_ID) #list moths with potential issues
#13 flights to check
#of these, 5 were deemed to not need trimming (moths did not land or exit through curtain)
#for the others, apply trimming based on frame numbers for stop counts, and manual checks of the footage
#also checked that scored outcomes match new end points of tracks

MULTITRACKS_LIGHT<-MULTITRACKS_LIGHT%>%
  mutate(trim = case_when( #create new variable, trim, with trim = 1 if frame should be removed, else trim=0
    Unique_Flight_ID == "M417_2025_CL_0.03lx" & Frame>1007 ~ 1,
    Unique_Flight_ID == "M470_2024_DL_0.03lx" & Frame>162 ~ 1,
    Unique_Flight_ID == "M499_2024_CL_0.03lx" & Frame>56 ~ 1,
    Unique_Flight_ID == "M465_2025_DL_0.03lx" & Frame<44 ~ 1,
    Unique_Flight_ID == "M463_2025_DL_0.03lx" & Frame>130 ~ 1,
    Unique_Flight_ID == "M490_2024_CL_0.03lx" & Frame>94 ~ 1,
    Unique_Flight_ID == "M459_2025_CL_0.03lx" & Frame>72 ~ 1,
    Unique_Flight_ID == "M502_2024_CL_0.03lx" & Frame>147 ~ 1,
    .default = 0
  ))%>%
  filter(trim==0)#remove frames to be trimmed

#3) identify any tracks with overly large displacements, indicating likely tracking errors

#calculate displacement (in 3D) between points, per moth track
TEMPTRACKS2<-list()

for (i in 1: length(unique(MULTITRACKS_LIGHT$Unique_Flight_ID))){
  
  flightid<-unique(MULTITRACKS_LIGHT$Unique_Flight_ID)[i]
  
  TEMPTRACKS2[[i]]<-MULTITRACKS_LIGHT%>%
    filter(Unique_Flight_ID ==flightid)%>%
    arrange(Frame, by = "Unique_Flight_ID")
  TEMPTRACKS2[[i]]$Displacement<-rep(0, length(TEMPTRACKS2[[i]]$X))
  
  for (j in 2:length(TEMPTRACKS2[[i]]$X)){
    TEMPTRACKS2[[i]]$Displacement[j]<-sqrt((TEMPTRACKS2[[i]]$X_smooth[j]-TEMPTRACKS2[[i]]$X_smooth[j-1])^2+
                                            (TEMPTRACKS2[[i]]$Y_smooth[j]-TEMPTRACKS2[[i]]$Y_smooth[j-1])^2+
                                            (TEMPTRACKS2[[i]]$Z_smooth[j]-TEMPTRACKS2[[i]]$Z_smooth[j-1])^2)
  }
}
#recombine into single dataframe
MULTITRACKS_LIGHT<-do.call("rbind", TEMPTRACKS2)

#identify tracks with odd displacements
hist(MULTITRACKS_LIGHT$Displacement, breaks=100)
summary(MULTITRACKS_LIGHT$Displacement)
LARGE2<-MULTITRACKS_LIGHT%>%
  filter(Displacement> 0.5)
unique(LARGE2$Unique_Flight_ID)#11 - fits with observations from track plots
  #keep these in dataset for the moment, for analysis of flight outcomes etc.
  #but the error-prone segments should be removed for tortuosity analysis
length(unique(MULTITRACKS_LIGHT$Unique_Flight_ID))#279 flights remain for analysis

#data cleaning done

#Recode variables as for info dataset above
for (i in 1:length(MULTITRACKS_LIGHT$Treatment)){
  MULTITRACKS_LIGHT$LightStructure[i]<-str_split(MULTITRACKS_LIGHT$Treatment, "_")[[i]][1]
  MULTITRACKS_LIGHT$BgdLightLevel[i]<-str_split(MULTITRACKS_LIGHT$Treatment, "_")[[i]][2]
}
factors <- c("Unique_Flight_ID","Unique_Moth_ID","Moth_ID", "Treatment","LatinName","Family",
             "CaptureMethod","Location","TrialNumber","Year","LightStructure", "BgdLightLevel")
MULTITRACKS_LIGHT<-MULTITRACKS_LIGHT%>%
  mutate(LightStructure=recode(LightStructure,
                               "DL"="Multiple","CL"="Single","SingleC" ="Single 10lx"))%>%
  mutate(BgdLightLevel=recode(BgdLightLevel,
                              "0.03lx"="Low","0.3lx"="Medium","3lx"="High",))%>%
  mutate(across(all_of(factors), as.factor))%>%
  mutate(BgdLightLevel = factor(BgdLightLevel,
                                levels = c("Low","Medium","High")))%>%
  mutate(LightStructure = factor(LightStructure,
                                 levels = c("Single","Multiple","Single 10lx")))%>%
  mutate(Treatment = factor(Treatment,
                            levels = c("CL_0.03lx","DL_0.03lx","CL_0.3lx","DL_0.3lx","CL_3lx","DL_3lx","SingleC_0.3lx")))
str(MULTITRACKS_LIGHT)

#remove Single 10lx light treatment, and save for supplementary analyses
SUPP_MULTITRACKS<-MULTITRACKS_LIGHT%>%
  filter(BgdLightLevel=="Medium")%>%
  filter(Year=="2024")%>%
  droplevels()
str(SUPP_MULTITRACKS)
length(unique((SUPP_MULTITRACKS$Unique_Flight_ID)))#93 flights here

MULTITRACKS_LIGHT<-MULTITRACKS_LIGHT%>%
  filter(LightStructure!="Single 10lx")%>%
  droplevels()
str(MULTITRACKS_LIGHT)

#Key numbers

#number of flights to analyse:
length(unique(MULTITRACKS_LIGHT$Unique_Flight_ID))#264 flights to analyse
MULTITRACKS_LIGHT%>%
  group_by(Treatment, Unique_Flight_ID)%>%
  summarise(n=n())%>%
  group_by(Treatment)%>%
  summarise(n=n())#32 to 58 flights per treatment

#break down by species and family
MULTITRACKS_LIGHT%>%
       group_by(LatinName, Unique_Flight_ID)%>%
       summarise(n=n())%>%
       group_by(LatinName)%>%
       summarise(n=n())#from 27 different species
FAMILIES_MULTI<-MULTITRACKS_LIGHT%>%
  group_by(Family,Treatment, Unique_Moth_ID)%>%
  summarise(n=n())%>%
  group_by(Family, Treatment)%>%
  summarise(n=n())
ggplot(FAMILIES_MULTI, aes(fill=Family, y=n, x=Treatment, label=n)) + 
  geom_bar(position="fill", stat="identity")+
  theme_bw()+ylab("Proportion of flights")+
  geom_text(size = 3, position = position_fill(vjust = 0.5))
#family composition is similar across treatment groups, except for lack of Crambidae in DL_3lx (high background light, multiple focals)

#number of species
MULTITRACKSPECIES<-unique(MULTITRACKS_LIGHT$LatinName)

#total number of species across experiments 1 and 2
unique(c(MULTITRACKSPECIES, SINGLETRACKSPECIES))#39 different species in total across experiments
setdiff(MULTITRACKSPECIES, SINGLETRACKSPECIES)#5 species in experiment 2 but not in experiment 1
setdiff(SINGLETRACKSPECIES,MULTITRACKSPECIES)#12 species in experiment 1 but not experiment 2

#1) Take-off probability----

#Key numbers
INFO_MULTI%>%
  group_by(TakeOff)%>%
  summarise(n=n()) #853 flight attempts; of these, 19 excluded, 539 did not fly within 1 minute, 295 took off

#Plot proportion of successful take-offs

#summarise by treatment
SUMMARY_INFO_MULTI<-INFO_MULTI%>%
  filter(!is.na(TakeOff))%>%#exclude NAs
  group_by(Treatment,TakeOff)%>%
  summarise(n=n())%>%
  ungroup()%>%
  mutate(TakeOff = factor(TakeOff,
                          levels = c("0","1")))
  
#alternative with barplot
plot_TO_expt2<-ggplot(SUMMARY_INFO_MULTI, aes(fill=TakeOff, col=TakeOff,y=n, x=Treatment, label=n)) + 
  geom_bar(position="fill", stat="identity")+
  ylab("Flight success (% trials)")+
  labs(fill="Take-off")+
  scale_y_continuous(expand=c(0,0),breaks=c(0, 0.25, 0.5, 0.75, 1), labels= c(0,25,50,75,100))+
  
  scale_x_discrete(labels = c("Single\nLow", "Multiple\nLow",
                              "Single\nMedium", "Multiple\nMedium",
                              "Single\nHigh", "Multiple\nHigh"))+
  scale_fill_manual(values = c("white","#19A4FA"),labels=c('no flight', 'flight'))+
  scale_colour_manual(values=c("#19A4FA","#19A4FA"))+
  geom_text(size = 3, position = position_fill(vjust = 0.5), col="grey40")+
  theme(legend.text = element_text(size=10))+
  theme_classic()+
  ggtitle("B")+
  guides(color="none")+guides(fill="none")+
  theme(axis.title.x = element_text(size = 12),
        axis.text.x = element_text(size = 10),
        axis.title.y = element_text(size = 12),
        axis.title=element_text(size=14),
        axis.text.y=element_text(size=10))
plot_TO_expt2

#model probability of take-off

#a) Is there a difference between years (for repeated treatments)?

YEARTEST_MULTI<-INFO_MULTI%>%
  filter(BgdLightLevel=="Low")%>%
  droplevels()
str(YEARTEST_MULTI)

#plot
SUMMARY_YEARTEST_MULTI<-YEARTEST_MULTI%>%
  filter(!is.na(TakeOff))%>%
  group_by(Treatment,Year, TakeOff)%>%
  summarise(n=n())%>%
  ungroup()
focallights<-c("single focal light","multiple focal lights")
names(focallights)<-c("CL_0.03lx","DL_0.03lx")
SUMMARY_YEARTEST_MULTI$TakeOff<-as.factor(SUMMARY_YEARTEST_MULTI$TakeOff)
ggplot(SUMMARY_YEARTEST_MULTI, aes(x=Year,y=n,fill=TakeOff,col= TakeOff,label=n)) + 
  geom_bar(position="fill", stat="identity")+
  facet_wrap(.~Treatment,labeller = labeller(Treatment = focallights))+
  scale_fill_manual(values = c("white","#19A4FA"),labels=c('no flight', 'flight'))+
  scale_colour_manual(values=c("#19A4FA","#19A4FA"))+
  theme_classic()+ylab("Proportion of flights")+
  geom_text(size = 3, position = position_fill(vjust = 0.5), col="grey40")+
  guides(col="none", fill="none")

#model
year_mod_multi1<-glmmTMB(TakeOff ~ Treatment*Year+TrialNumber+CaptureMethod+
                     (1|Unique_Moth_ID)+(1|LatinName)+(1|Family),
                   family=binomial, data=YEARTEST_MULTI)
summary(year_mod_multi1)
year_mod_multi1_resid <- simulateResiduals(year_mod_multi1)
plot(year_mod_multi1_resid)#good

year_mod_multi1b<-glmmTMB(TakeOff ~ Treatment+TrialNumber+Year+CaptureMethod+
                      (1|Unique_Moth_ID)+(1|LatinName)+(1|Family),
                    family=binomial, data=YEARTEST_MULTI)
anova(year_mod_multi1, year_mod_multi1b)#can remove interaction
summary(year_mod_multi1b)

year_mod_multi1c<-glmmTMB(TakeOff ~ Treatment+TrialNumber+CaptureMethod+
                      (1|Unique_Moth_ID)+(1|LatinName)+(1|Family),
                    family=binomial, data=YEARTEST_MULTI)
anova(year_mod_multi1c, year_mod_multi1b)#can remove year

#So can move on to analyse data from 2024 & 2025 together

#b) probability of take-off between treatments

#model with treatment only
INFO_MULTI$BgdLightLevel<-factor(INFO_MULTI$BgdLightLevel, levels =c("Low", "Medium","High"))
prob_mod_multi1<-glmmTMB(TakeOff ~ Treatment*CaptureMethod+TrialNumber+
                     (1|Unique_Moth_ID)+(1|LatinName)+(1|Family),
                   family=binomial, data=INFO_MULTI)
summary(prob_mod_multi1)
prob_mod_multi1_resid <- simulateResiduals(prob_mod_multi1)
plot(prob_mod_multi1_resid)#good

prob_mod_multi1b<-glmmTMB(TakeOff ~ Treatment+TrialNumber+CaptureMethod+
                      (1|Unique_Moth_ID)+(1|LatinName)+(1|Family),
                    family=binomial, data=INFO_MULTI)
anova(prob_mod_multi1, prob_mod_multi1b)#can remove interaction
summary(prob_mod_multi1b)

prob_mod_multi1c<-glmmTMB(TakeOff ~ Treatment+CaptureMethod+
                      (1|Unique_Moth_ID)+(1|LatinName)+(1|Family),
                    family=binomial, data=INFO_MULTI)
anova(prob_mod_multi1c, prob_mod_multi1b)#significant effect of trial number

prob_mod_multi1d<-glmmTMB(TakeOff ~ Treatment+TrialNumber+
                      (1|Unique_Moth_ID)+(1|LatinName)+(1|Family),
                    family=binomial, data=INFO_MULTI)
anova(prob_mod_multi1b, prob_mod_multi1d)#significant effect of capture method

prob_mod_multi1e<-glmmTMB(TakeOff ~ CaptureMethod+TrialNumber+
                      (1|Unique_Moth_ID)+(1|LatinName)+(1|Family),
                    family=binomial, data=INFO_MULTI)
anova(prob_mod_multi1b, prob_mod_multi1e)#significant effect of treatment

#significant effects of treatment number, capture method & treatment
summary(glht(prob_mod_multi1b, linfct = mcp(Treatment = "Tukey"))) #all significant pairwise comparisons involve the highest bgd light level

#move on to more useful model with light structure and background light level
prob_mod_multi2<-glmmTMB(TakeOff ~ LightStructure*BgdLightLevel+TrialNumber+CaptureMethod+
                     (1|Unique_Moth_ID)+(1|LatinName)+(1|Family),
                   family=binomial, data=INFO_MULTI)
summary(prob_mod_multi2)
prob_mod_multi2_resid <- simulateResiduals(prob_mod_multi2)
plot(prob_mod_multi2_resid)#good

prob_mod_multi2b<-glmmTMB(TakeOff ~ LightStructure+BgdLightLevel+TrialNumber+CaptureMethod+
                      (1|Unique_Moth_ID)+(1|LatinName)+(1|Family),
                    family=binomial, data=INFO_MULTI)
anova(prob_mod_multi2b, prob_mod_multi2)#interaction just not significant
summary(prob_mod_multi2b)
prob_mod_multi2c<-glmmTMB(TakeOff ~ TrialNumber+BgdLightLevel+CaptureMethod+
                      (1|Unique_Moth_ID)+(1|LatinName)+(1|Family),
                    family=binomial, data=INFO_MULTI)
anova(prob_mod_multi2b, prob_mod_multi2c)#no effect of structure
summary(prob_mod_multi2c)
prob_mod_multi2d<-glmmTMB(TakeOff ~ BgdLightLevel+CaptureMethod+
                      (1|Unique_Moth_ID)+(1|LatinName)+(1|Family),
                    family=binomial, data=INFO_MULTI)
anova(prob_mod_multi2d, prob_mod_multi2c)#significant effect of trial number

prob_mod_multi2e<-glmmTMB(TakeOff ~ BgdLightLevel+TrialNumber+
                      (1|Unique_Moth_ID)+(1|LatinName)+(1|Family),
                    family=binomial, data=INFO_MULTI)
anova(prob_mod_multi2c, prob_mod_multi2e)#significant effect of capture method

prob_mod_multi2f<-glmmTMB(TakeOff ~ TrialNumber+CaptureMethod+
                      (1|Unique_Moth_ID)+(1|LatinName)+(1|Family),
                    family=binomial, data=INFO_MULTI)
anova(prob_mod_multi2c, prob_mod_multi2f)#significant effect of background lighting

summary(glht(prob_mod_multi2c, linfct = mcp(BgdLightLevel = "Tukey"))) 
  #take-offs significantly lower in high lighting treatment than others

#odds ratios
summary(prob_mod_multi2c)
plogis(0.240)#56% probability of taking off under low light treatment, for moths caught with nets, in first trial
plogis(0.240-1.251)#27% probability of taking off under high light
exp(-1.251)#odds ratio 0.286, odds of taking off are reduced by over two thirds for high bgd light level compared to low

#odds for capture method and trial number 
exp(-0.699)
exp(-0.550)

#odds ratios-relevel factors for easier interpretation
INFO_MULTI_relevel<-INFO_MULTI
INFO_MULTI_relevel$BgdLightLevel<-factor(INFO_MULTI_relevel$BgdLightLevel, levels=c("Medium","Low","High"))
prob_mod2c<-glmmTMB(TakeOff ~ BgdLightLevel+TrialNumber+CaptureMethod+
                      (1|Unique_Moth_ID)+(1|LatinName)+(1|Family),
                    family=binomial, data=INFO_MULTI_relevel)
summary(prob_mod2c)
exp(-1.233)#odds ratio 0.291, odds of taking off are reduced by over two thirds from medium to high light treatment

#plot binomial model results
plot_TO_expt2_capture<-plot_model(prob_mod2c, type = "pred", 
                                  terms = c("CaptureMethod"),
                                  colors = "darkgreen",
                                  title = "F")+
  theme_classic()+
  xlab ("Capture method")+ylab("Predicted flight probability (%)")+
  scale_y_continuous(limits=c(0.1,0.8),breaks = c(0.2, 0.4, 0.6,0.8), labels = c(20,40,60,80))+
  theme(axis.title.x = element_text(size = 12),
        axis.title = element_text(size = 14),
        axis.text.x = element_text(size = 10),
        axis.title.y = element_text(size = 12),
        axis.text.y=element_text(size=10))
plot_TO_expt2_capture
plot_TO_expt2_trial<-plot_model(prob_mod2c, type = "pred", 
                                terms = c("TrialNumber"),
                                colors = "chocolate4",
                                title = "D")+
  theme_classic()+
  xlab ("Trial number")+ylab("Predicted flight probability (%)")+
  scale_y_continuous(limits=c(0.1,0.8),breaks = c(0.2, 0.4, 0.6,0.8), labels = c(20,40,60,80))+
  theme(axis.title.x = element_text(size = 12),
        axis.title = element_text(size = 14),
        axis.text.x = element_text(size = 10),
        axis.title.y = element_text(size = 12),
        axis.text.y=element_text(size=10))
plot_TO_expt2_trial
plot_TO_expt2_bgd<-plot_model(prob_mod2c, type = "pred", 
                              terms = c("BgdLightLevel"),
                              colors = "bw",
                              title = "C")+
  theme_classic()+
  scale_y_continuous(limits=c(0.1,0.8),breaks = c(0.2, 0.4, 0.6,0.8), labels = c(20,40,60,80))+
  xlab ("Background light level")+ylab("Predicted flight probability (%)")+
  theme(axis.title.x = element_text(size = 12),
        axis.title = element_text(size = 14),
        axis.text.x = element_text(size = 10),
        axis.title.y = element_text(size = 12),
        axis.text.y=element_text(size=10))
plot_TO_expt2_bgd

#plot both experiments together - Figure 2 
plot_TO_expt1/((plot_TO_expt2 | ((plot_TO_expt2_bgd+plot_TO_expt2_trial) / 
                                   (plot_TO_expt1_capture+plot_TO_expt2_capture))))+
  plot_layout(axis_titles = "collect")

#2) Outcomes - flight to light----

#keep only outcome information for flights that meet analysis criteria
OUTCOMES_MULTI<-INFO_MULTI%>%
  filter(Unique_Flight_ID %in% MULTITRACKS_LIGHT$Unique_Flight_ID)
dim(OUTCOMES_MULTI)#264 flights for analysis
str(OUTCOMES_MULTI)

OUTCOMES_MULTI%>%
  group_by(Treatment)%>%
  summarise(n=n())#32 to 58 per treatment combination in experiment 2

OUTCOMES_MULTI$Outcome_simpler<-factor(OUTCOMES_MULTI$Outcome_simpler,
                                       levels=c("at light","up","down","other"))
#overall proportion of flights ending at the light:
OUTCOMES_MULTI%>%
  group_by(FlighttoLight_tracked)%>%
  summarise(n=n())%>%
  ungroup()
82/(182+82)#31% flights ended at the light

OUTCOMES_MULTI%>%
  group_by(Down_tracked)%>%
  summarise(n=n())%>%
  ungroup()

OUTCOME_SUMMARY_MULTI<-OUTCOMES_MULTI%>%
  group_by(Treatment, Outcome_simpler, Year)%>%
  summarise(n=n())%>%
  ungroup()
OUTCOME_SUMMARY_MULTI$Outcome_simpler<-factor(OUTCOME_SUMMARY_MULTI$Outcome_simpler,
                                       levels=c("other","down","up","at light"))

outcomes_expt2_byyear<-ggplot(OUTCOME_SUMMARY_MULTI, aes(fill=Outcome_simpler, y=n, x=Treatment, label=n)) + 
  geom_bar(position="fill", stat="identity")+
  theme_bw()+
  geom_text(size = 3, position = position_fill(vjust = 0.5))+
  labs(fill="Flight outcome")+
  facet_grid(. ~ Year, scales = "free", space = "free")+ 
  ylab("Flight outcomes (% trials)")+
  scale_y_continuous(breaks=c(0,0.25, 0.5, 0.75, 1), labels = c(0,25,50,75,100))+
  ggtitle("E")+
  scale_fill_manual(values = c("#B3B3B3","#66C2A5","#E78AC3","#FFD92F"))+
  geom_text(size = 3, position = position_fill(vjust = 0.5), col="grey50")+  theme(legend.text = element_text(size=10))+
  theme_classic()+
  scale_x_discrete(drop = TRUE)+
  theme(axis.title.x = element_text(size = 12),
        axis.title = element_text(size = 14),
        axis.text.x = element_text(size = 10),
        axis.title.y = element_text(size = 12),
        axis.text.y=element_text(size=10))

outcomes_expt2_byyear #needs x axis labels updating manually

#a) test for difference between years

#model
OUTCOMES_YEAR_MULTI<-OUTCOMES_MULTI%>%
  filter(Treatment=="CL_0.03lx"|Treatment=="DL_0.03lx")%>%
  droplevels()
year_mod_multi2<-glmmTMB(FlighttoLight_tracked ~ Treatment*Year+TrialNumber+CaptureMethod+
                     (1|Unique_Moth_ID)+(1|LatinName)+(1|Family),
                   family=binomial, data=OUTCOMES_YEAR_MULTI)
summary(year_mod_multi2)
year_mod_multi2_resid <- simulateResiduals(year_mod_multi2)
plot(year_mod_multi2_resid)#good

year_mod_multi2b<-glmmTMB(FlighttoLight_tracked ~ Treatment+Year+TrialNumber+CaptureMethod+
                      (1|Unique_Moth_ID)+(1|LatinName)+(1|Family),
                    family=binomial, data=OUTCOMES_YEAR_MULTI)

anova(year_mod_multi2, year_mod_multi2b)#remove interaction
summary(year_mod_multi2b)

year_mod_multi2c<-glmmTMB(FlighttoLight_tracked ~ Treatment+Year+TrialNumber+
                      (1|Unique_Moth_ID)+(1|LatinName)+(1|Family),
                    family=binomial, data=OUTCOMES_YEAR_MULTI)

anova(year_mod_multi2c, year_mod_multi2b)
summary(year_mod_multi2c)

year_mod_multi2d<-glmmTMB(FlighttoLight_tracked ~ Treatment+Year+
                      (1|Unique_Moth_ID)+(1|LatinName)+(1|Family),
                    family=binomial, data=OUTCOMES_YEAR_MULTI)

anova(year_mod_multi2c, year_mod_multi2d)
summary(year_mod_multi2d)

year_mod_multi2e<-glmmTMB(FlighttoLight_tracked ~ Year+
                      (1|Unique_Moth_ID)+(1|LatinName)+(1|Family),
                    family=binomial, data=OUTCOMES_YEAR_MULTI)

anova(year_mod_multi2d, year_mod_multi2e)
summary(year_mod_multi2e)

year_mod_multi2f<-glmmTMB(FlighttoLight_tracked ~ 1+
                      (1|Unique_Moth_ID)+(1|LatinName)+(1|Family),
                    family=binomial, data=OUTCOMES_YEAR_MULTI)
anova(year_mod_multi2f, year_mod_multi2e) #here, significant effect of year

#b) model for each year - 2024 (low and medium bgd light levels)

OUTCOMES_MULTI_24<-OUTCOMES_MULTI%>%
  filter(Year=="2024")%>%
  droplevels()

ftl_mod_multi1<-glmmTMB(FlighttoLight_tracked ~ LightStructure*BgdLightLevel+TrialNumber+CaptureMethod+
                    (1|Unique_Moth_ID)+(1|LatinName)+(1|Family),
                  family=binomial, data=OUTCOMES_MULTI_24)
ftl_mod_multi1 <- update(ftl_mod_multi1,
                             control=glmmTMBControl(optimizer=optim,
                                                    optArgs=list(method="BFGS")))

summary(ftl_mod_multi1)
ftl_mod_multi1_resid <- simulateResiduals(ftl_mod_multi1)
plot(ftl_mod_multi1_resid)#good

ftl_mod_multi1b<-glmmTMB(FlighttoLight_tracked ~ LightStructure+BgdLightLevel+TrialNumber+CaptureMethod+
                     (1|Unique_Moth_ID)+(1|LatinName)+(1|Family),
                   family=binomial, data=OUTCOMES_MULTI_24)
anova(ftl_mod_multi1, ftl_mod_multi1b)#remove interaction
summary(ftl_mod_multi1b)

ftl_mod_multi1c<-glmmTMB(FlighttoLight_tracked ~ LightStructure+BgdLightLevel+CaptureMethod+
                     (1|Unique_Moth_ID)+(1|LatinName)+(1|Family),
                   family=binomial, data=OUTCOMES_MULTI_24)
anova(ftl_mod_multi1c, ftl_mod_multi1b)#remove trial number
summary(ftl_mod_multi1c)

ftl_mod_multi1d<-glmmTMB(FlighttoLight_tracked ~ BgdLightLevel+CaptureMethod+
                     (1|Unique_Moth_ID)+(1|LatinName)+(1|Family),
                   family=binomial, data=OUTCOMES_MULTI_24)
anova(ftl_mod_multi1c, ftl_mod_multi1d)#remove light structure
summary(ftl_mod_multi1d)

ftl_mod_multi1e<-glmmTMB(FlighttoLight_tracked ~ CaptureMethod+
                     (1|Unique_Moth_ID)+(1|LatinName)+(1|Family),
                   family=binomial, data=OUTCOMES_MULTI_24)
anova(ftl_mod_multi1e, ftl_mod_multi1d)#significant effect of bgd light level

ftl_mod_multi1f<-glmmTMB(FlighttoLight_tracked ~ BgdLightLevel+
                     (1|Unique_Moth_ID)+(1|LatinName)+(1|Family),
                   family=binomial, data=OUTCOMES_MULTI_24)
anova(ftl_mod_multi1d, ftl_mod_multi1f)#significant effect of capture method 

summary(ftl_mod_multi1d)
exp(1.073)#OR=2.924 for light trapped moths compared to netted
exp(0.999)#OR=2.716 for medium bgd compared to low

plot_ftl_expt2_bgd_2024<-plot_model(ftl_mod_multi1d, type = "pred", terms = c("BgdLightLevel"),
                                    colors="bw", title ="H")+
  theme_classic()+
  scale_y_continuous(limits=c(0,0.9),breaks = c(0,0.2, 0.4, 0.6,0.8), 
                     labels = c(0,20,40,60,80), expand =c(0,0))+
  xlab("Background lighting level")+ylab("Predicted probability of flight-to-light (%)")+
  theme(axis.title.x = element_text(size = 12),
        axis.title = element_text(size = 16),
        axis.text.x = element_text(size = 10),
        axis.title.y = element_text(size = 12),
        axis.text.y=element_text(size=10))
plot_ftl_expt2_bgd_2024

plot_ftl_expt2_capture_2024<-plot_model(ftl_mod_multi1d, type = "pred", 
                                        terms = c("CaptureMethod"),
                                    colors="darkgreen", title ="I")+
  theme_classic()+
  scale_y_continuous(limits=c(0,0.9),breaks = c(0,0.2, 0.4, 0.6,0.8), 
                     labels = c(0,20,40,60,80), expand =c(0,0))+
  xlab("Capture method")+ylab("Predicted probability of flight-to-light (%)")+
  theme(axis.title.x = element_text(size = 12),
        axis.title = element_text(size = 14),
        axis.text.x = element_text(size = 10),
        axis.title.y = element_text(size = 12),
        axis.text.y=element_text(size=10))
plot_ftl_expt2_capture_2024

#c) model for 2025
OUTCOMES_MULTI_25<-OUTCOMES_MULTI%>%
  filter(Year=="2025")

ftl_mod_multi2<-glmmTMB(FlighttoLight_tracked ~ LightStructure*BgdLightLevel+TrialNumber+CaptureMethod+
                    (1|Unique_Moth_ID)+(1|LatinName)+(1|Family),
                  family=binomial, data=OUTCOMES_MULTI_25)
summary(ftl_mod_multi2)
ftl_mod_multi2_resid <- simulateResiduals(ftl_mod_multi2)
plot(ftl_mod_multi2_resid)#good

ftl_mod_multi2b<-glmmTMB(FlighttoLight_tracked ~ LightStructure+BgdLightLevel+TrialNumber+CaptureMethod+
                     (1|Unique_Moth_ID)+(1|LatinName)+(1|Family),
                   family=binomial, data=OUTCOMES_MULTI_25)

anova(ftl_mod_multi2, ftl_mod_multi2b)#significant interaction

ftl_mod_multi2c<-glmmTMB(FlighttoLight_tracked ~ LightStructure*BgdLightLevel+CaptureMethod+
                     (1|Unique_Moth_ID)+(1|LatinName)+(1|Family),
                   family=binomial, data=OUTCOMES_MULTI_25)
anova(ftl_mod_multi2c, ftl_mod_multi2)#no effect of trial number
summary(ftl_mod_multi2c)

ftl_mod_multi2d<-glmmTMB(FlighttoLight_tracked ~ LightStructure*BgdLightLevel+
                     (1|Unique_Moth_ID)+(1|LatinName)+(1|Family),
                   family=binomial, data=OUTCOMES_MULTI_25)
anova(ftl_mod_multi2c, ftl_mod_multi2d)#no effect of capture method
summary(ftl_mod_multi2d)
ftl_mod_multi2e<-glmmTMB(FlighttoLight_tracked ~ BgdLightLevel+LightStructure+
                     (1|Unique_Moth_ID)+(1|LatinName)+(1|Family),
                   family=binomial, data=OUTCOMES_MULTI_25)
anova(ftl_mod_multi2e, ftl_mod_multi2d)#cannot remove interaction

#pairwise comparisons within focal light structure, and within background light level
emmres_ftl<-emmeans(ftl_mod_multi2d,  ~ BgdLightLevel | LightStructure)
pwpm(emmres_ftl)
pairs(emmres_ftl)
plot(emmres_ftl, comparisons=TRUE)
emmres_ftl2<-emmeans(ftl_mod_multi2d,  ~ LightStructure|  BgdLightLevel)
pwpm(emmres_ftl2)
pairs(emmres_ftl2)

#plot model results
plot_ftl_expt2_bgd_2025<-plot_model(ftl_mod_multi2d, type = "pred", terms = c("BgdLightLevel","LightStructure"),
                                    colors = c("grey10","grey70"),
                                    title ="J")+
  theme_classic()+
  scale_y_continuous(limits=c(0,0.9),breaks = c(0,0.2, 0.4, 0.6,0.8), 
                     labels = c(0,20,40,60,80), expand =c(0,0))+
  labs(col="Focal light \nstructure")+
  xlab("Background light level")+ylab("Predicted probability of flight-to-light (%)")+
  theme(axis.title.x = element_text(size = 12),
        axis.title = element_text(size = 14),
        axis.text.x = element_text(size = 10),
        axis.title.y = element_text(size = 12),
        axis.text.y=element_text(size=10))
plot_ftl_expt2_bgd_2025

#3) Outcomes - up----

SUMMARY_YEARTEST_MULTI<-OUTCOMES_MULTI%>%
  filter(Treatment=="CL_0.03lx"|Treatment=="DL_0.03lx")%>%
  group_by(Treatment,Year, Up_tracked)%>%
  summarise(n=n())%>%
  ungroup()
#overall proportion of flights ending up:
OUTCOMES_MULTI%>%
  group_by(Up_tracked)%>%
  summarise(n=n())%>%
  ungroup()
137/(127+137)#52% flights went up

#a) test for difference between years
SUMMARY_YEARTEST_MULTI$Up_tracked<-as.factor(SUMMARY_YEARTEST_MULTI$Up_tracked)
ggplot(SUMMARY_YEARTEST_MULTI, aes(fill=Up_tracked, y=n, x=Year, label=n)) + 
  geom_bar(position="fill", stat="identity")+
  facet_wrap(.~Treatment)+
  theme_classic()+ylab("Proportion of flights")+
  geom_text(size = 3, position = position_fill(vjust = 0.5))+
  guides(fill="none")#more flight to light in 2025

#model
year_mod_multi3<-glmmTMB(Up_tracked ~ Treatment*Year+TrialNumber+CaptureMethod+
                     (1|Unique_Moth_ID)+(1|LatinName)+(1|Family),
                   family=binomial, data=OUTCOMES_YEAR_MULTI)
summary(year_mod_multi3)
year_mod_multi3_resid <- simulateResiduals(year_mod_multi3)
plot(year_mod_multi3_resid)#good

year_mod_multi3b<-glmmTMB(Up_tracked ~ Treatment+Year+TrialNumber+CaptureMethod+
                      (1|Unique_Moth_ID)+(1|LatinName)+(1|Family),
                    family=binomial, data=OUTCOMES_YEAR_MULTI)
anova(year_mod_multi3, year_mod_multi3b)#remove interaction
summary(year_mod_multi3b)

year_mod_multi3c<-glmmTMB(Up_tracked ~ Treatment+CaptureMethod+TrialNumber+
                      (1|Unique_Moth_ID)+(1|LatinName)+(1|Family),
                    family=binomial, data=OUTCOMES_YEAR_MULTI)
anova(year_mod_multi3c, year_mod_multi3b)#can remove year
summary(year_mod_multi3c)
#so analyse all together

#model across both years
up_mod_multi1<-glmmTMB(Up_tracked ~ LightStructure*BgdLightLevel+TrialNumber+CaptureMethod+
                     (1|Unique_Moth_ID)+(1|LatinName)+(1|Family),
                   family=binomial, data=OUTCOMES_MULTI)
summary(up_mod_multi1)
up_mod_multi1_resid <- simulateResiduals(up_mod_multi1)
plot(up_mod_multi1_resid)#good

up_mod_multi1b<-glmmTMB(Up_tracked ~ LightStructure+BgdLightLevel+TrialNumber+CaptureMethod+
                      (1|Unique_Moth_ID)+(1|LatinName)+(1|Family),
                    family=binomial, data=OUTCOMES_MULTI)
anova(up_mod_multi1b, up_mod_multi1)#can remove interaction
summary(up_mod_multi1b)

up_mod_multi1c<-glmmTMB(Up_tracked ~ LightStructure+BgdLightLevel+CaptureMethod+
                    (1|Unique_Moth_ID)+(1|LatinName)+(1|Family),
                  family=binomial, data=OUTCOMES_MULTI)
anova(up_mod_multi1b, up_mod_multi1c)#can remove trial number
summary(up_mod_multi1c)

up_mod_multi1d<-glmmTMB(Up_tracked ~ BgdLightLevel+CaptureMethod+
                    (1|Unique_Moth_ID)+(1|LatinName)+(1|Family),
                  family=binomial, data=OUTCOMES_MULTI)
anova(up_mod_multi1d, up_mod_multi1c)#can remove light structure
summary(up_mod_multi1d)

up_mod_multi1e<-glmmTMB(Up_tracked ~ CaptureMethod+
                    (1|Unique_Moth_ID)+(1|LatinName)+(1|Family),
                  family=binomial, data=OUTCOMES_MULTI)
anova(up_mod_multi1d, up_mod_multi1e)#significant effect of bgd light level

up_mod_multi1f<-glmmTMB(Up_tracked ~BgdLightLevel+
                    (1|Unique_Moth_ID)+(1|LatinName)+(1|Family),
                  family=binomial, data=OUTCOMES_MULTI)
anova(up_mod_multi1f, up_mod_multi1d)#significant effect of capture method

summary(glht(up_mod_multi1d, linfct = mcp(BgdLightLevel = "Tukey"))) 
#high light condition differs from two others - more likely to fly up in high light conditions

#odds ratios for effect of low- high, low - medium, medium - high bgd light level, and capture method for trap
exp(1.803)
exp(0.443)
exp(1.359)
exp(-0.711)

#plot model results
plot_model(up_mod_multi1d, type = "pred", 
           terms = c("BgdLightLevel"),
           colors = "bw",
           title = "")+
  theme_classic()+  xlab ("Background light level")+ylab("Predicted upwards probability (%)")+
  theme(axis.title.x = element_text(size = 12),
        axis.title = element_text(size = 14),
        axis.text.x = element_text(size = 10),
        axis.title.y = element_text(size = 12),
        axis.text.y=element_text(size=10))

#4) Erratic spiraling----

OUTCOMES_MULTI%>%
  group_by(Erratic_spiralling)%>%
  summarise(n=n())%>%
  ungroup()
57/264#erratic spiraling in 22 % flights overall

#plot 
SPIRAL_SUMMARY<-OUTCOMES_MULTI%>%
  group_by(Erratic_spiralling, Treatment)%>%
  summarise(n=n())%>%
  ungroup()

SPIRAL_SUMMARY$Erratic_spiralling<-factor(SPIRAL_SUMMARY$Erratic_spiralling, levels=c("0","1"))
ggplot(SPIRAL_SUMMARY, aes(fill=Erratic_spiralling, y=n, x=Treatment, label=n)) + 
  geom_bar(position="fill", stat="identity")+
  ylab("Proportion of trials")+
  labs(fill="Erratic spiraling")+
  scale_fill_manual(values = c("grey70","violetred3"))+
  geom_text(size = 3, position = position_fill(vjust = 0.5), col="grey30")+
  theme(legend.text = element_text(size=10))+
  theme_classic()+
  theme(axis.title.x = element_text(size = 12),
        axis.text.x = element_text(size = 10),
        axis.title.y = element_text(size = 12),
        axis.text.y=element_text(size=10))+
  scale_y_continuous(expand=c(0,0))#+guides(fill="none")


#a) test for difference between years

year_mod_multi4<-glmmTMB(Erratic_spiralling ~ Treatment*Year+TrialNumber+CaptureMethod+
                     (1|Unique_Moth_ID)+(1|LatinName)+(1|Family),
                   family=binomial, data=OUTCOMES_YEAR_MULTI)
summary(year_mod_multi4)
year_mod_multi4_resid <- simulateResiduals(year_mod_multi4)
plot(year_mod_multi4_resid)#ok

year_mod_multi4b<-glmmTMB(Erratic_spiralling ~ Treatment+Year+TrialNumber+CaptureMethod+
                      (1|Unique_Moth_ID)+(1|LatinName)+(1|Family),
                    family=binomial, data=OUTCOMES_YEAR_MULTI)

anova(year_mod_multi4, year_mod_multi4b)#remove interaction
summary(year_mod_multi4b)

year_mod_multi4c<-glmmTMB(Erratic_spiralling ~ Treatment+TrialNumber+CaptureMethod+
                      (1|Unique_Moth_ID)+(1|LatinName)+(1|Family),
                    family=binomial, data=OUTCOMES_YEAR_MULTI)
anova(year_mod_multi4c, year_mod_multi4b)#remove year

#b) model across both years
str(OUTCOMES_MULTI)
OUTCOMES_MULTI<-OUTCOMES_MULTI%>%
  mutate(AnySpiral = case_when(
    Spiral_up ==1|Spiral_down==1|Spiral_other==1 ~1,
    .default=0))
spiral_mod_multi1<-glmmTMB(Erratic_spiralling ~ LightStructure*BgdLightLevel+TrialNumber+CaptureMethod+
                   (1|Unique_Moth_ID)+(1|LatinName)+(1|Family),
                 family=binomial, data=OUTCOMES_MULTI)
summary(spiral_mod_multi1)
spiral_mod_multi1_resid <- simulateResiduals(spiral_mod_multi1)
plot(spiral_mod_multi1_resid)#good

spiral_mod_multi1b<-glmmTMB(Erratic_spiralling ~ LightStructure+BgdLightLevel+CaptureMethod+TrialNumber+
                    (1|Unique_Moth_ID)+(1|LatinName)+(1|Family),
                  family=binomial, data=OUTCOMES_MULTI)
anova(spiral_mod_multi1b, spiral_mod_multi1)#can remove interaction
summary(spiral_mod_multi1b)

spiral_mod_multi1c<-glmmTMB(Erratic_spiralling ~ LightStructure+BgdLightLevel+CaptureMethod+
                    (1|Unique_Moth_ID)+(1|LatinName)+(1|Family),
                  family=binomial, data=OUTCOMES_MULTI)#remove trial number
anova(spiral_mod_multi1b, spiral_mod_multi1c)
summary(spiral_mod_multi1c)

spiral_mod_multi1d<-glmmTMB(Erratic_spiralling ~ LightStructure+BgdLightLevel+
                    (1|Unique_Moth_ID)+(1|LatinName)+(1|Family),
                  family=binomial, data=OUTCOMES_MULTI)
anova(spiral_mod_multi1d, spiral_mod_multi1c)#remove capture method
summary(spiral_mod_multi1d)

spiral_mod_multi1e<-glmmTMB(Erratic_spiralling ~ LightStructure+
                    (1|Unique_Moth_ID)+(1|LatinName)+(1|Family),
                  family=binomial, data=OUTCOMES_MULTI)
anova(spiral_mod_multi1d, spiral_mod_multi1e)#cannot remove bgd light level
summary(spiral_mod_multi1e)

spiral_mod_multi1f<-glmmTMB(Erratic_spiralling ~BgdLightLevel+
                        (1|Unique_Moth_ID)+(1|LatinName)+(1|Family),
                      family=binomial, data=OUTCOMES_MULTI)
anova(spiral_mod_multi1f, spiral_mod_multi1d)#significant effect of light structure
summary(spiral_mod_multi1d)
summary(glht(spiral_mod_multi1d, linfct = mcp(BgdLightLevel = "Tukey"))) 
#OR for multiple lights: 
exp(-0.779)#half as likely to see erratic spiraling under multiple lights 

#plot model results
plot_sp_expt2b<-plot_model(spiral_mod_multi1d, type = "pred", 
                          terms = c("LightStructure","BgdLightLevel"),
                          colors = c("grey30","grey50","grey90"),
                          title = "E")+
  theme_classic()+
  scale_y_continuous(limits=c(0,0.5),breaks = c(0,0.2, 0.4), 
                     labels = c(0,20,40), expand =c(0,0))+
  xlab("Focal light structure")+ylab("Predicted probability of erratic spiralling (%)")+
  theme(axis.title.x = element_text(size = 12),
        axis.title = element_text(size = 14),
        axis.text.x = element_text(size = 10),
        axis.title.y = element_text(size = 12),
        axis.text.y=element_text(size=10))
plot_sp_expt2b

#c) what about any form of spiraling?

#make variable for record of any spiral observed, and update datasets
OUTCOMES_MULTI<-OUTCOMES_MULTI%>%
  mutate(AnySpiral = case_when(
    Spiral_up ==1|Spiral_down==1|Spiral_other==1 ~1,
    .default=0))
OUTCOMES_YEAR_MULTI<-OUTCOMES_MULTI%>%
  filter(Treatment=="CL_0.03lx"|Treatment=="DL_0.03lx")%>%
  droplevels()
ANYSPIRAL_SUMMARY_MULTI<-OUTCOMES_MULTI%>%
  group_by(Treatment, AnySpiral,Year)%>%
  summarise(n=n())%>%
  ungroup()

OUTCOMES_MULTI%>%
  group_by(AnySpiral)%>%
  summarise(n=n())
135/264#spiral in 51% of trials

ANYSPIRAL_SUMMARY_MULTI$AnySpiral<-as.factor(ANYSPIRAL_SUMMARY_MULTI$AnySpiral)
ggplot(ANYSPIRAL_SUMMARY_MULTI, aes(fill=AnySpiral, y=n, x=Treatment, label=n)) + 
  geom_bar(position="fill", stat="identity")+
  theme_bw()+
  facet_grid(.~Year, scales = "free", space = "free")+
  geom_text(size = 3, position = position_fill(vjust = 0.5))+
  labs(fill="Spiralling")+
  ylab("Spiral observed (% trials)")+
  scale_y_continuous(breaks=c(0,0.25, 0.5, 0.75, 1), labels = c(0,25,50,75,100))+
  geom_text(size = 3, position = position_fill(vjust = 0.5), col="grey50")+  theme(legend.text = element_text(size=10))+
  theme_classic()+
  scale_x_discrete(drop = TRUE)+
  theme(axis.title.x = element_text(size = 12),
        axis.title = element_text(size=16),
        axis.text.x = element_text(angle = 45, hjust = 1, size=10),
        axis.title.y = element_text(size = 12),
        axis.text.y=element_text(size=10))

#any spiral - test for difference between years

year_mod_multi5<-glmmTMB(AnySpiral ~ Treatment*Year+TrialNumber+CaptureMethod+
                     (1|Unique_Moth_ID)+(1|LatinName)+(1|Family),
                   family=binomial, data=OUTCOMES_YEAR_MULTI)
summary(year_mod_multi5)
year_mod_multi5_resid <- simulateResiduals(year_mod_multi5)
plot(year_mod_multi5_resid)#ok

year_mod_multi5b<-glmmTMB(AnySpiral ~ Treatment+Year+TrialNumber+CaptureMethod+
                      (1|Unique_Moth_ID)+(1|LatinName)+(1|Family),
                    family=binomial, data=OUTCOMES_YEAR_MULTI)

anova(year_mod_multi5, year_mod_multi5b)#remove interaction
summary(year_mod_multi5b)

year_mod_multi5c<-glmmTMB(AnySpiral ~ Treatment+TrialNumber+CaptureMethod+
                      (1|Unique_Moth_ID)+(1|LatinName)+(1|Family),
                    family=binomial, data=OUTCOMES_YEAR_MULTI)

anova(year_mod_multi5c, year_mod_multi5b)#can't remove year - more spiraling in 2025

#model for each year separately - 2024
OUTCOMES_MULTI_24<-OUTCOMES_MULTI%>%
  filter(Year=="2024")%>%
  droplevels()
OUTCOMES_MULTI_25<-OUTCOMES_MULTI%>%
  filter(Year=="2025")%>%
  droplevels()

spiral_mod_multi2<-glmmTMB(AnySpiral ~ LightStructure*BgdLightLevel+TrialNumber+CaptureMethod+
                       (1|Unique_Moth_ID)+(1|LatinName)+(1|Family),
                     family=binomial, data=OUTCOMES_MULTI_24)
summary(spiral_mod_multi2)
spiral_mod_multi2_resid <- simulateResiduals(spiral_mod_multi2)
plot(spiral_mod_multi2_resid)#ok

spiral_mod_multi2b<-glmmTMB(AnySpiral ~ LightStructure+BgdLightLevel+TrialNumber+CaptureMethod+
                        (1|Unique_Moth_ID)+(1|LatinName)+(1|Family),
                      family=binomial, data=OUTCOMES_MULTI_24)
anova(spiral_mod_multi2b, spiral_mod_multi2)
summary(spiral_mod_multi2b)

spiral_mod_multi2c<-glmmTMB(AnySpiral ~ LightStructure+BgdLightLevel+TrialNumber+
                        (1|Unique_Moth_ID)+(1|LatinName)+(1|Family),
                      family=binomial, data=OUTCOMES_MULTI_24)
anova(spiral_mod_multi2b, spiral_mod_multi2c)#remove capture method
summary(spiral_mod_multi2c)

spiral_mod_multi2d<-glmmTMB(AnySpiral ~ LightStructure+BgdLightLevel+
                        (1|Unique_Moth_ID)+(1|LatinName)+(1|Family),
                      family=binomial, data=OUTCOMES_MULTI_24)
anova(spiral_mod_multi2d, spiral_mod_multi2c)#can remove trial number
summary(spiral_mod_multi2d)

spiral_mod_multi2e<-glmmTMB(AnySpiral ~ LightStructure+
                        (1|Unique_Moth_ID)+(1|LatinName)+(1|Family),
                      family=binomial, data=OUTCOMES_MULTI_24)
anova(spiral_mod_multi2d, spiral_mod_multi2e)#can remove bgd light level
summary(spiral_mod_multi2e)

spiral_mod_multi2f<-glmmTMB(AnySpiral ~1+
                        (1|Unique_Moth_ID)+(1|LatinName)+(1|Family),
                      family=binomial, data=OUTCOMES_MULTI_24)
anova(spiral_mod_multi2f, spiral_mod_multi2e)#significant effect of structure, more spiraling with single light

#OR for multiple lights: 
exp(-0.904)#OR=0.405, less than half the probability of seeing spirals in flights under multiple compared to single lights

plot_model(spiral_mod_multi2e, type = "pred", 
                          terms = c("LightStructure"),
                          colors = "violetred3",
                          title = "")+
  theme_classic()+
  xlab("Focal light structure")+ylab("Predicted probability of any spirals")+
  theme(axis.title.x = element_text(size = 12),
        axis.title = element_text(size = 14),
        axis.text.x = element_text(size = 10),
        axis.title.y = element_text(size = 12),
        axis.text.y=element_text(size=10))


#2025
spiral_mod_multi3<-glmmTMB(AnySpiral ~ LightStructure*BgdLightLevel+TrialNumber+CaptureMethod+
                       (1|Unique_Moth_ID)+(1|LatinName)+(1|Family),
                     family=binomial, data=OUTCOMES_MULTI_25)
summary(spiral_mod_multi3)
spiral_mod_multi3_resid <- simulateResiduals(spiral_mod_multi3)
plot(spiral_mod_multi3_resid)#ok

spiral_mod_multi3b<-glmmTMB(AnySpiral ~ LightStructure+BgdLightLevel+TrialNumber+CaptureMethod+
                        (1|Unique_Moth_ID)+(1|LatinName)+(1|Family),
                      family=binomial, data=OUTCOMES_MULTI_25)
anova(spiral_mod_multi3b, spiral_mod_multi3)#can remove interaction
summary(spiral_mod_multi3b)

spiral_mod_multi3c<-glmmTMB(AnySpiral ~ LightStructure+BgdLightLevel+TrialNumber+
                        (1|Unique_Moth_ID)+(1|LatinName)+(1|Family),
                      family=binomial, data=OUTCOMES_MULTI_25)
anova(spiral_mod_multi3b, spiral_mod_multi3c)#remove capture method
summary(spiral_mod_multi3c)

spiral_mod_multi3d<-glmmTMB(AnySpiral ~ LightStructure+BgdLightLevel+
                        (1|Unique_Moth_ID)+(1|LatinName)+(1|Family),
                      family=binomial, data=OUTCOMES_MULTI_25)
anova(spiral_mod_multi3d, spiral_mod_multi3c)#can remove trial number
summary(spiral_mod_multi3d)

spiral_mod_multi3e<-glmmTMB(AnySpiral ~ BgdLightLevel+
                        (1|Unique_Moth_ID)+(1|LatinName)+(1|Family),
                      family=binomial, data=OUTCOMES_MULTI_25)
anova(spiral_mod_multi3d, spiral_mod_multi3e)#can remove structure
summary(spiral_mod_multi3e)

spiral_mod_multi3f<-glmmTMB(AnySpiral ~1+
                        (1|Unique_Moth_ID)+(1|LatinName)+(1|Family),
                      family=binomial, data=OUTCOMES_MULTI_25)
anova(spiral_mod_multi3f, spiral_mod_multi3e)#significant effect of bgd level, more spiraling with low bgd light level

#OR for high light level: 
exp(-1.502)#OR=0.223, far smaller chance of seeing a spiral under high background light

plot_model(spiral_mod_multi3e, type = "pred", 
                           terms = c("BgdLightLevel"),
                           colors = "violetred3",
                           title = "")+
  theme_classic()+
  xlab("Focal light structure")+ylab("Predicted probability of any spirals")+
  theme(axis.title.x = element_text(size = 12),
        axis.title = element_text(size = 14),
        axis.text.x = element_text(size = 10),
        axis.title.y = element_text(size = 12),
        axis.text.y=element_text(size=10))


#Plot all results for flight outcomes, experiments 1 and 2 (Figure 3)----

outcomes_single+outcomes_expt2_byyear+plot_layout(guides="collect")

plot_ftl_expt1_intensity+ plot_ftl_expt1_capture+ plot_ftl_expt2_bgd_2024+
  plot_ftl_expt2_capture_2024+plot_ftl_expt2_bgd_2025+plot_layout(ncol=5, axis_titles = "collect")


#5)Trajectory analyses----

#calculate tortuosity and other metrics----

#apply turning angle function to each unique flight id
FlightIDs<-unique(MULTITRACKS_LIGHT$Unique_Flight_ID)
FLIGHTS<-list()

for (i in 1:length(FlightIDs)){
  TEMPDF<-MULTITRACKS_LIGHT%>%
    filter(Unique_Flight_ID ==FlightIDs[i])
  str(TEMPDF)
  FLIGHTS[[i]]<-calculate_turning_angles_and_lengths(TEMPDF)#create new list with data with turning angles for each flight
}

#before calculating summary metrics for each track, remove points identified as having large displacements in 3D,
#indicating error in 3D tracking
FLIGHTS<-lapply(FLIGHTS, function(x) filter(x, Displacement < 0.5))

#combine into larger dataframe
MULTITRACKS_ANGLES <- bind_rows(FLIGHTS)

#apply tortuosity function to each flight
MULTI_TORTUOSITY_LIST <- lapply(FLIGHTS, calculate_tortuosity_metrics)
length(MULTI_TORTUOSITY_LIST)
head(MULTI_TORTUOSITY_LIST[[1]])

#combine tortuosity results
MULTITRACKS_METRICS <- do.call(rbind,MULTI_TORTUOSITY_LIST)
str(MULTITRACKS_METRICS)

#plot results
col_names <- c("Unique_Moth_ID","Treatment","LatinName","Family","CaptureMethod","Location")
MULTITRACKS_METRICS <- MULTITRACKS_METRICS %>%
  mutate(across(all_of(col_names), as.factor))
str(MULTITRACKS_METRICS)
levels(MULTITRACKS_METRICS$Treatment)

plot_tortuosity_multi <- ggplot(MULTITRACKS_METRICS, aes(x = Treatment, y = mean_angle_3D, fill = Treatment)) +
  geom_boxplot() +
  theme_classic() +
  labs(title = "", y = "Mean turning angle (degrees)", x = NULL) +
  theme(
    axis.text = element_text(size = 16),
    axis.title = element_text(size = 18),
    axis.text.x = element_text(angle = 45, hjust = 1),
    legend.position = "none",
    panel.grid.major = element_blank(),
    panel.grid.minor = element_blank(),
    panel.border = element_blank(),  # keep axis lines, so we don't add a full box
    axis.line = element_line(color = "black")  # this adds back the x and y axis lines
  )
plot_tortuosity_multi

plot_SD_multi <- ggplot(MULTITRACKS_METRICS, aes(x = Treatment, y = sd_angle_3D, fill = Treatment)) +
  geom_boxplot() +
  theme_classic() +  # Clean theme
  labs(title = "", y = "SD turning angle") +  # Remove x axis label
  theme(
    axis.text = element_text(size = 16),
    axis.title = element_text(size = 18),
    axis.text.x = element_text(angle = 45, hjust = 1),
    legend.position = "none",
    panel.grid.major = element_blank(),
    panel.grid.minor = element_blank(),
    panel.border = element_blank(),  # keep axis lines, so we don't add a full box
    axis.line = element_line(color = "black"))
plot_SD_multi

#statistics on overall metrics - mean turning angle ----

#a) check if there is a difference between years
MULTITRACKS_METRICS_YEAR<-MULTITRACKS_METRICS%>%
  filter(BgdLightLevel=="Low")%>%
  droplevels()
year_tort_multi1<-lmer(mean_angle_3D ~ Treatment*Year+TrialNumber+CaptureMethod+ (1|Unique_Moth_ID)+(1|Family) + (1|LatinName), data=MULTITRACKS_METRICS_YEAR)
  #singular fit
length(MULTITRACKS_METRICS_YEAR$Unique_Moth_ID)
dim(MULTITRACKS_METRICS_YEAR)
length(unique(MULTITRACKS_METRICS_YEAR$Unique_Moth_ID))#only 115-85=30 moths have replicate flights, so estimating effect of moth ID will be tricky 

#test which random effects are most useful
year_tort_multi1<-lmer(mean_angle_3D ~ Treatment*Year+CaptureMethod+TrialNumber+ (1|Unique_Moth_ID) + (1|Family), data=MULTITRACKS_METRICS_YEAR)
year_tort_multi2<-lmer(mean_angle_3D ~ Treatment*Year+CaptureMethod+TrialNumber+ (1|Unique_Moth_ID) + (1|LatinName), data=MULTITRACKS_METRICS_YEAR)
year_tort_multi3<-lmer(mean_angle_3D ~ Treatment*Year+CaptureMethod+ (1|Family) + (1|LatinName), data=MULTITRACKS_METRICS_YEAR)#singular fit
AIC(year_tort_multi1, year_tort_multi2, year_tort_multi3)#model w/o family is best
summary(year_tort_multi2)
year_tort_multi2_resid <- simulateResiduals(year_tort_multi2)
plot(year_tort_multi2_resid)#ok

year_tort_multi2b<-lmer(mean_angle_3D ~ Treatment+Year+TrialNumber+CaptureMethod+(1|Unique_Moth_ID)+(1|LatinName), data=MULTITRACKS_METRICS_YEAR)
anova(year_tort_multi2, year_tort_multi2b)
summary(year_tort_multi2b)#remove interaction
year_tort_multi2c<-lmer(mean_angle_3D ~ Treatment+TrialNumber+CaptureMethod+(1|Unique_Moth_ID)+(1|LatinName), data=MULTITRACKS_METRICS_YEAR)
anova(year_tort_multi2c, year_tort_multi2b)#cannot remove year - more tortuosity in 2025

#b) model for each year - 2024
MULTITRACKS_METRICS_2024<-MULTITRACKS_METRICS%>%
  filter(Year=="2024")%>%
  droplevels()
tort_model1_24 <- lmer(mean_angle_3D ~ LightStructure*BgdLightLevel+TrialNumber+CaptureMethod+(1|Family)+ (1|LatinName)+ (1|Unique_Moth_ID), data=MULTITRACKS_METRICS_2024)
  #singular fit with all random effects, try removing them one by one
tort_model1_24 <- lmer(mean_angle_3D ~ LightStructure*BgdLightLevel+TrialNumber+CaptureMethod+(1|Family)+ (1|LatinName), data=MULTITRACKS_METRICS_2024)
tort_model2_24 <- lmer(mean_angle_3D ~ LightStructure*BgdLightLevel+TrialNumber+CaptureMethod+(1|Unique_Moth_ID)+ (1|LatinName), data=MULTITRACKS_METRICS_2024)
tort_model3_24 <- lmer(mean_angle_3D ~ LightStructure*BgdLightLevel+TrialNumber+CaptureMethod+(1|Family)+ (1|Unique_Moth_ID), data=MULTITRACKS_METRICS_2024)
AIC(tort_model1_24,tort_model2_24,tort_model3_24)  #best model is with family and species

summary(tort_model1_24)
tort_model1_resi <- simulateResiduals(tort_model1_24)
plot(tort_model1_resi)#good
tort_model1_24b <- lmer(mean_angle_3D ~ LightStructure+BgdLightLevel+TrialNumber+CaptureMethod+(1|Family)+ (1|LatinName), data=MULTITRACKS_METRICS_2024)
anova(tort_model1_24b,tort_model1_24) #remove interaction
summary(tort_model1_24b)
tort_model1_24c <- lmer(mean_angle_3D ~ LightStructure+BgdLightLevel+CaptureMethod+(1|Family)+ (1|LatinName), data=MULTITRACKS_METRICS_2024)
anova(tort_model1_24b,tort_model1_24c) #remove trial number
summary(tort_model1_24c)
tort_model1_24d <- lmer(mean_angle_3D ~ LightStructure+CaptureMethod+(1|Family)+ (1|LatinName), data=MULTITRACKS_METRICS_2024)
anova(tort_model1_24d,tort_model1_24c) #remove bgd light level
summary(tort_model1_24d)
tort_model1_24e <- lmer(mean_angle_3D ~ LightStructure+(1|Family)+ (1|LatinName), data=MULTITRACKS_METRICS_2024)
anova(tort_model1_24d,tort_model1_24e) #remove capture method
summary(tort_model1_24e)
tort_model1_24f <- lmer(mean_angle_3D ~ 1+(1|Family)+(1|LatinName), data=MULTITRACKS_METRICS_2024)
anova(tort_model1_24f,tort_model1_24e) #remove light structure - no fixed effects in 2024

#c) model for each year  - 2025
MULTITRACKS_METRICS_2025<-MULTITRACKS_METRICS%>%
  filter(Year=="2025")%>%
  droplevels()
tort_model1_25 <- lmer(mean_angle_3D ~ LightStructure*BgdLightLevel+TrialNumber+CaptureMethod+(1|Unique_Moth_ID)+(1|Family)+ (1|LatinName), data=MULTITRACKS_METRICS_2025)
summary(tort_model1_25)
tort_model1_25_resi <- simulateResiduals(tort_model1_25)
plot(tort_model1_25_resi)#poor
tort_model2_25 <- lmer(log(mean_angle_3D) ~ LightStructure*BgdLightLevel+TrialNumber+CaptureMethod+(1|Unique_Moth_ID)+(1|Family)+ (1|LatinName), data=MULTITRACKS_METRICS_2025)
tort_model2_25_resi <- simulateResiduals(tort_model2_25)
plot(tort_model2_25_resi)#better with log transform
summary(tort_model2_25)

tort_model2_25b <- lmer(log(mean_angle_3D) ~ LightStructure+BgdLightLevel+TrialNumber+CaptureMethod+(1|Family)+ (1|LatinName)+(1|Unique_Moth_ID), data=MULTITRACKS_METRICS_2025)
anova(tort_model2_25b,tort_model2_25) #remove interaction
summary(tort_model2_25b)
tort_model2_25c <- lmer(log(mean_angle_3D) ~ LightStructure+BgdLightLevel+CaptureMethod+(1|Family)+ (1|LatinName)+ (1|Unique_Moth_ID), data=MULTITRACKS_METRICS_2025)
anova(tort_model2_25b,tort_model2_25c) #remove trial number
summary(tort_model2_25c)
tort_model2_25d <- lmer(log(mean_angle_3D) ~ LightStructure+BgdLightLevel+(1|Family)+ (1|LatinName)+ (1|Unique_Moth_ID), data=MULTITRACKS_METRICS_2025)
anova(tort_model2_25d,tort_model2_25c) #remove capture method
summary(tort_model2_25d)
tort_model2_25e <- lmer(log(mean_angle_3D) ~ BgdLightLevel+(1|Family)+ (1|LatinName)+ (1|Unique_Moth_ID), data=MULTITRACKS_METRICS_2025)
anova(tort_model2_25d,tort_model2_25e)#remove structure, just
summary(tort_model2_25e)
tort_model2_25f <- lmer(log(mean_angle_3D) ~1+(1|Family)+ (1|LatinName)+ (1|Unique_Moth_ID), data=MULTITRACKS_METRICS_2025)
anova(tort_model2_25f,tort_model2_25e)
#significant effect of background light level only - tortuosity lower under high light, in 2025

#plot results for 2025
plot_tort_level <- ggplot(MULTITRACKS_METRICS_2025, aes(x = BgdLightLevel, y = log(mean_angle_3D), fill = LightStructure)) +
  geom_boxplot() +
  theme_classic() +
  labs(title = "", y = "Mean turning angle (degrees)", x = NULL) +
  theme(
    axis.text = element_text(size = 16),
    axis.title = element_text(size = 18),
    axis.text.x = element_text(angle = 45, hjust = 1),
    legend.position = "none",
    panel.grid.major = element_blank(),
    panel.grid.minor = element_blank(),
    panel.border = element_blank(),  
    axis.line = element_line(color = "black") 
  )
plot_tort_level

#statistics on overall metrics - SD of turning angle----

#a) test for a difference between years
year_sd_multi1<-lmer(sd_angle_3D ~ Treatment*Year+TrialNumber+CaptureMethod+ (1|Unique_Moth_ID)+ (1|LatinName), data=MULTITRACKS_METRICS_YEAR)
resid_sd_multi1<-simulateResiduals(year_sd_multi1)#good
plot(resid_sd_multi1)
summary(year_sd_multi1)

#proceed without family for the moment:

#try removing interaction
year_sd_multi1b<-lmer(sd_angle_3D ~ Treatment+Year+TrialNumber+CaptureMethod+ (1|Unique_Moth_ID) + (1|LatinName), data=MULTITRACKS_METRICS_YEAR)
anova(year_sd_multi1, year_sd1b)
summary(year_sd_multi1b)#remove interaction
year_sd_multi1c<-lmer(sd_angle_3D ~ Treatment+TrialNumber+CaptureMethod+(1|Unique_Moth_ID)+(1|LatinName), data=MULTITRACKS_METRICS_YEAR)
anova(year_sd_multi1c, year_sd_multi1b)#can remove year

#so model across both years

sd_model1 <- lmer(sd_angle_3D ~ LightStructure*BgdLightLevel+TrialNumber+CaptureMethod+ (1|Unique_Moth_ID)+(1|Family)+ (1|LatinName), data=MULTITRACKS_METRICS)
#singular fit
sd_model2 <- lmer(sd_angle_3D ~ LightStructure*BgdLightLevel+TrialNumber+CaptureMethod+ (1|Unique_Moth_ID)+(1|Family), data=MULTITRACKS_METRICS)
sd_model3 <- lmer(sd_angle_3D ~ LightStructure*BgdLightLevel+TrialNumber+CaptureMethod+ (1|Unique_Moth_ID)+ (1|LatinName), data=MULTITRACKS_METRICS)
sd_model4 <- lmer(sd_angle_3D ~ LightStructure*BgdLightLevel+TrialNumber+CaptureMethod+ (1|Family)+ (1|LatinName), data=MULTITRACKS_METRICS)
AIC(sd_model1, sd_model2, sd_model3, sd_model4)#best model is with family and species
summary(sd_model4)
sd_model4_resi <- simulateResiduals(sd_model4)
plot(sd_model4_resi)#good
sd_model4b <- lmer(sd_angle_3D ~ LightStructure+BgdLightLevel+TrialNumber+CaptureMethod+(1|Family)+ (1|LatinName), data=MULTITRACKS_METRICS)
anova(sd_model4, sd_model4b) #remove interaction
summary(sd_model4b)
sd_model4c <- lmer(sd_angle_3D ~ LightStructure+BgdLightLevel+CaptureMethod+(1|Family)+ (1|LatinName), data=MULTITRACKS_METRICS)
anova(sd_model4b,sd_model4c) #remove trial number
summary(sd_model4c)
sd_model4d <- lmer(sd_angle_3D ~ LightStructure+BgdLightLevel+(1|Family)+ (1|LatinName), data=MULTITRACKS_METRICS)
anova(sd_model4d,sd_model4c) #remove capture method
summary(sd_model4d)
sd_model4e <- lmer(sd_angle_3D ~ BgdLightLevel+(1|Family)+ (1|LatinName), data=MULTITRACKS_METRICS)
anova(sd_model4d,sd_model4e) #cannot remove light structure
summary(sd_model4e)
sd_model4f <- lmer(sd_angle_3D ~ LightStructure+(1|Family)+(1|LatinName), data=MULTITRACKS_METRICS)
anova(sd_model4f,sd_model4d) #significant effect of light level

summary(glht(sd_model4d, linfct = mcp(BgdLightLevel = "Tukey"))) 
#the only significant comparison is high-low bgd lighting

plot_sd_level <- ggplot(MULTITRACKS_METRICS, aes(x = BgdLightLevel, y = sd_angle_3D)) +
  geom_boxplot() +
  theme_classic() +
  labs(title = "", y = "SD turning angle (degrees)", x = NULL) +
  theme(
    axis.text = element_text(size = 16),
    axis.title = element_text(size = 18),
    axis.text.x = element_text(angle = 45, hjust = 1),
    legend.position = "none",
    panel.grid.major = element_blank(),
    panel.grid.minor = element_blank(),
    panel.border = element_blank(),  
    axis.line = element_line(color = "black") 
  )
plot_sd_level

#flight metrics before and after exposure to light----

#create variable to signify whether or not (0 no and 1 yes) the moth is exposed by facing the light 
MULTITRACKS_ANGLES$Exposed <- ifelse(MULTITRACKS_ANGLES$bearing_to_light_deg <= 135, 1, 0)

#combine with position after light cone to create variable for when moth has both passed the light and has been exposed to it
MULTITRACKS_ANGLES$After_Light_Both_Exposed <- ifelse(
  !is.na(MULTITRACKS_ANGLES$Exposed) & 
    !is.na(MULTITRACKS_ANGLES$After_Light_Both) & 
    MULTITRACKS_ANGLES$Exposed == 1 & MULTITRACKS_ANGLES$After_Light_Both == 1,
  1,
  0
)

MULTITRACKS_ANGLES <- MULTITRACKS_ANGLES %>%
  arrange(Unique_Flight_ID, Frame) %>%# sort chronologically within each track
  group_by(Unique_Flight_ID) %>%
  mutate(
    After_Light_Both_Exposed = if_else(
      cumsum(After_Light_Both_Exposed) > 0,  # once a 1 appears, stays 1 afterward
      1,
      0
    )
  ) %>%
  ungroup()

#create variable to separate flights before and after exposure to light
MULTITRACKS_ANGLES$UniqueSegment<-paste(MULTITRACKS_ANGLES$Unique_Flight_ID,MULTITRACKS_ANGLES$After_Light_Both_Exposed, sep="_")
str(MULTITRACKS_ANGLES)

#identify flights with at least 5 points before and after light exposure 
WITHBOTH<-MULTITRACKS_ANGLES%>%
  group_by(Unique_Flight_ID,After_Light_Both_Exposed)%>%
  summarise(n=n())%>%
  filter(n>=5)%>%
  group_by(Unique_Flight_ID)%>%
  summarise(n=n())%>%
  filter(n==2)
unique(WITHBOTH$Unique_Flight_ID)#244 flights out of 264 initially - very little loss

#restrict track data to only those flights
TRACKS_BEFOREAFTER<-MULTITRACKS_ANGLES%>%
  filter(Unique_Flight_ID %in% WITHBOTH$Unique_Flight_ID)

#apply function to calculate tortuosity for these segments

#split by unique segment IDs - turning angles etc already calculated
segmentIDs<-unique(TRACKS_BEFOREAFTER$UniqueSegment)
SEGMENTS<-list()

for (i in 1:length(segmentIDs)){
  SEGMENTS[[i]]<-TRACKS_BEFOREAFTER%>%
    filter(UniqueSegment == segmentIDs[i])
  }

# Apply new tortuosity function to each dataframe in the results_list
MULTI_TORTUOSITY_LIST_SEGMENTS <- lapply(SEGMENTS, calculate_tortuosity_metrics)
length(MULTI_TORTUOSITY_LIST_SEGMENTS)#488 - 2 segments each for 244 flights
head(MULTI_TORTUOSITY_LIST_SEGMENTS[[1]])

# Combine the results into a single dataframe
MULTITRACKS_SEGMENTS <- do.call(rbind, MULTI_TORTUOSITY_LIST_SEGMENTS)
dim(MULTITRACKS_SEGMENTS)
#add a column for straightness index
MULTITRACKS_SEGMENTS$straighness_index <- MULTITRACKS_SEGMENTS$distance_first_last / MULTITRACKS_SEGMENTS$total_path_length

col_names2 <- c("Unique_Moth_ID","UniqueSegment","After_Light_Both_Exposed","Treatment","LatinName","Family","CaptureMethod","Location")
MULTITRACKS_SEGMENTS<- MULTITRACKS_SEGMENTS %>%
  mutate(across(all_of(col_names2), as.factor))
str(MULTITRACKS_SEGMENTS)

#number of flight paths per treatment
with(MULTITRACKS_SEGMENTS,table(Treatment, After_Light_Both_Exposed))

#statistics on flight segments - mean turning angle ----

#a) for 2024 
MULTITRACKS_METRICS_2024<-MULTITRACKS_SEGMENTS%>%
  filter(Year=="2024")%>%
  droplevels()
ba_tort_model1_24 <- lmer(mean_angle_3D ~ Treatment*After_Light_Both_Exposed+TrialNumber+CaptureMethod+(1|Family) + (1|LatinName)+(1|Unique_Moth_ID)+(1|Unique_Flight_ID), data=MULTITRACKS_METRICS_2024)
  #ideally models should include moth id (for repeated trials) and flight id (each track now has two segments)
  #this leads to singular fit, and unique moth id may be challenging to fit, due to small number of moths with repeated successful trials
  #focus on flight id
ba_tort_model2_24 <- lmer(mean_angle_3D ~ Treatment*After_Light_Both_Exposed+TrialNumber+CaptureMethod+(1|Family) + (1|LatinName)+(1|Unique_Flight_ID), data=MULTITRACKS_METRICS_2024)
  #still  singular fit, reduce number of random effects
ba_tort_model3_24 <- lmer(mean_angle_3D ~ Treatment*After_Light_Both_Exposed+TrialNumber+CaptureMethod+(1|Unique_Flight_ID), data=MULTITRACKS_METRICS_2024)
ba_tort_model4_24 <- lmer(mean_angle_3D ~ Treatment*After_Light_Both_Exposed+TrialNumber+CaptureMethod+(1|Family) + (1|Unique_Flight_ID), data=MULTITRACKS_METRICS_2024)
ba_tort_model5_24 <- lmer(mean_angle_3D ~ Treatment*After_Light_Both_Exposed+TrialNumber+CaptureMethod+ (1|LatinName)+(1|Unique_Flight_ID), data=MULTITRACKS_METRICS_2024)
ba_tort_model6_24 <- lmer(mean_angle_3D ~ Treatment*After_Light_Both_Exposed+TrialNumber+CaptureMethod+ (1|LatinName)+(1|Family), data=MULTITRACKS_METRICS_2024)
AIC(ba_tort_model2_24,ba_tort_model3_24,ba_tort_model4_24,ba_tort_model5_24,ba_tort_model6_24)
  #best model includes only family and species, but conceptually more important to account for repeated measures (flight id)
summary(ba_tort_model3_24)#this is the only model with flight ID that is not a singular fit

ba_tort_model3_resi <- simulateResiduals(ba_tort_model3_24)
plot(ba_tort_model3_resi)#ok
hist(resid(ba_tort_model1_24))
ba_tort_model3_24b <- lmer(mean_angle_3D ~ Treatment+After_Light_Both_Exposed+TrialNumber+CaptureMethod+(1|Unique_Flight_ID), data=MULTITRACKS_METRICS_2024)
anova(ba_tort_model3_24b,ba_tort_model3_24) #remove interaction
summary(ba_tort_model3_24b)
ba_tort_model3_24c <- lmer(mean_angle_3D ~ Treatment+After_Light_Both_Exposed+CaptureMethod+(1|Unique_Flight_ID), data=MULTITRACKS_METRICS_2024)
anova(ba_tort_model3_24b,ba_tort_model3_24c) #remove trial number
summary(ba_tort_model3_24c)
ba_tort_model3_24d <- lmer(mean_angle_3D ~ Treatment+After_Light_Both_Exposed+(1|Unique_Flight_ID), data=MULTITRACKS_METRICS_2024)
anova(ba_tort_model3_24d,ba_tort_model3_24c) #remove capture method
summary(ba_tort_model3_24d)
ba_tort_model3_24e <- lmer(mean_angle_3D ~ After_Light_Both_Exposed+(1|Unique_Flight_ID), data=MULTITRACKS_METRICS_2024)
anova(ba_tort_model3_24d,ba_tort_model3_24e) #remove treatment
summary(ba_tort_model3_24e)
ba_tort_model3_24f <- lmer(mean_angle_3D ~ 1+(1|Unique_Flight_ID), data=MULTITRACKS_METRICS_2024)
anova(ba_tort_model3_24f,ba_tort_model3_24e) #effect of crossing light, but otherwise no difference between treatments

#repeat analyses with light structure and background light level, rather than just treatment
ba_tort_model7_24 <- lmer(mean_angle_3D ~ BgdLightLevel*LightStructure*After_Light_Both_Exposed+TrialNumber+CaptureMethod+ (1|Family)+(1|LatinName), data=MULTITRACKS_METRICS_2024)
ba_tort_model8_24 <- lmer(mean_angle_3D ~ BgdLightLevel*LightStructure*After_Light_Both_Exposed+TrialNumber+CaptureMethod+ (1|Unique_Flight_ID)+(1|LatinName), data=MULTITRACKS_METRICS_2024)
ba_tort_model9_24 <- lmer(mean_angle_3D ~ BgdLightLevel*LightStructure*After_Light_Both_Exposed+TrialNumber+CaptureMethod+ (1|Family)+ (1|Unique_Flight_ID), data=MULTITRACKS_METRICS_2024)
ba_tort_model10_24 <- lmer(mean_angle_3D ~ BgdLightLevel*LightStructure*After_Light_Both_Exposed+TrialNumber+CaptureMethod+ (1|Unique_Flight_ID), data=MULTITRACKS_METRICS_2024)
AIC(ba_tort_model7_24,ba_tort_model8_24,ba_tort_model9_24,ba_tort_model10_24)#again, model with family and species is best, but conceptually important to use flight ID
summary(ba_tort_model10_24)
model1_resi <- simulateResiduals(ba_tort_model10_24)
plot(model1_resi)#ok

ba_tort_model10_24b <- lmer(mean_angle_3D ~ BgdLightLevel*LightStructure+BgdLightLevel*After_Light_Both_Exposed+LightStructure*After_Light_Both_Exposed+TrialNumber+CaptureMethod+ (1|Unique_Flight_ID), data=MULTITRACKS_METRICS_2024)
anova(ba_tort_model10_24,ba_tort_model10_24b)
summary(ba_tort_model10_24b)
ba_tort_model10_24c <- lmer(mean_angle_3D ~ BgdLightLevel*LightStructure+BgdLightLevel*After_Light_Both_Exposed+TrialNumber+CaptureMethod+ (1|Unique_Flight_ID), data=MULTITRACKS_METRICS_2024)
anova(ba_tort_model10_24c,ba_tort_model10_24b)
summary(ba_tort_model10_24c)
ba_tort_model10_24d <- lmer(mean_angle_3D ~ LightStructure+BgdLightLevel*After_Light_Both_Exposed+TrialNumber+CaptureMethod+ (1|Unique_Flight_ID), data=MULTITRACKS_METRICS_2024)
anova(ba_tort_model10_24c,ba_tort_model10_24d)
summary(ba_tort_model10_24d)
ba_tort_model10_24e <- lmer(mean_angle_3D ~ LightStructure+BgdLightLevel+After_Light_Both_Exposed+TrialNumber+CaptureMethod+ (1|Unique_Flight_ID), data=MULTITRACKS_METRICS_2024)
anova(ba_tort_model10_24e,ba_tort_model10_24d)
summary(ba_tort_model10_24e)
ba_tort_model10_24f<-lmer(mean_angle_3D ~ LightStructure+BgdLightLevel+After_Light_Both_Exposed+CaptureMethod+ (1|Unique_Flight_ID), data=MULTITRACKS_METRICS_2024)
anova(ba_tort_model10_24e,ba_tort_model10_24f)
summary(ba_tort_model10_24f)
ba_tort_model10_24g<-lmer(mean_angle_3D ~ BgdLightLevel+After_Light_Both_Exposed+CaptureMethod+ (1|Unique_Flight_ID), data=MULTITRACKS_METRICS_2024)
anova(ba_tort_model10_24g,ba_tort_model10_24f)
summary(ba_tort_model10_24g)
ba_tort_model10_24h<-lmer(mean_angle_3D ~ BgdLightLevel+After_Light_Both_Exposed+ (1|Unique_Flight_ID), data=MULTITRACKS_METRICS_2024)
anova(ba_tort_model10_24g,ba_tort_model10_24h)
summary(ba_tort_model10_24h)
ba_tort_model10_24i<-lmer(mean_angle_3D ~After_Light_Both_Exposed+ (1|Unique_Flight_ID), data=MULTITRACKS_METRICS_2024)
anova(ba_tort_model10_24i,ba_tort_model10_24h)
summary(ba_tort_model10_24i)
ba_tort_model10_24j<-lmer(mean_angle_3D ~1+ (1|Unique_Flight_ID), data=MULTITRACKS_METRICS_2024)
anova(ba_tort_model10_24i,ba_tort_model10_24j) #only exposure is significant

#check this would not change if best model (with family and species) was used
ba_tort_model7_24 <- lmer(mean_angle_3D ~ BgdLightLevel*LightStructure*After_Light_Both_Exposed+TrialNumber+CaptureMethod+ (1|Family)+(1|LatinName), data=MULTITRACKS_METRICS_2024)
summary(ba_tort_model7_24)
ba_tort_model7_24_resi <- simulateResiduals(ba_tort_model7_24)
plot(ba_tort_model7_24_resi)#ok

ba_tort_model7_24b <- lmer(mean_angle_3D ~ BgdLightLevel*LightStructure+BgdLightLevel*After_Light_Both_Exposed+LightStructure*After_Light_Both_Exposed+TrialNumber+CaptureMethod+ (1|Family)+(1|LatinName), data=MULTITRACKS_METRICS_2024)
anova(ba_tort_model7_24,ba_tort_model7_24b)
summary(ba_tort_model7_24b)
ba_tort_model7_24c <- lmer(mean_angle_3D ~ BgdLightLevel*LightStructure+BgdLightLevel*After_Light_Both_Exposed+TrialNumber+CaptureMethod+ (1|Family)+(1|LatinName), data=MULTITRACKS_METRICS_2024)
anova(ba_tort_model7_24c,ba_tort_model7_24b)
summary(ba_tort_model7_24c)
ba_tort_model7_24d <- lmer(mean_angle_3D ~ LightStructure+BgdLightLevel*After_Light_Both_Exposed+TrialNumber+CaptureMethod+ (1|Family)+(1|LatinName), data=MULTITRACKS_METRICS_2024)
anova(ba_tort_model7_24c,ba_tort_model7_24d)
summary(ba_tort_model7_24d)
ba_tort_model7_24e <- lmer(mean_angle_3D ~ LightStructure+BgdLightLevel+After_Light_Both_Exposed+TrialNumber+CaptureMethod+ (1|Family)+(1|LatinName), data=MULTITRACKS_METRICS_2024)
anova(ba_tort_model7_24e,ba_tort_model7_24d)
summary(ba_tort_model7_24e)
ba_tort_model7_24f <- lmer(mean_angle_3D ~ LightStructure+BgdLightLevel+After_Light_Both_Exposed+CaptureMethod+ (1|Family)+(1|LatinName), data=MULTITRACKS_METRICS_2024)
anova(ba_tort_model7_24e,ba_tort_model7_24f)
summary(ba_tort_model7_24f)
ba_tort_model7_24g <- lmer(mean_angle_3D ~ LightStructure+BgdLightLevel+After_Light_Both_Exposed+ (1|Family)+(1|LatinName), data=MULTITRACKS_METRICS_2024)
anova(ba_tort_model7_24g,ba_tort_model7_24f)
summary(ba_tort_model7_24g)
ba_tort_model7_24h <- lmer(mean_angle_3D ~ BgdLightLevel+After_Light_Both_Exposed+ (1|Family)+(1|LatinName), data=MULTITRACKS_METRICS_2024)
anova(ba_tort_model7_24g,ba_tort_model7_24h)
summary(ba_tort_model7_24h)
ba_tort_model7_24i <- lmer(mean_angle_3D ~ After_Light_Both_Exposed+ (1|Family)+(1|LatinName), data=MULTITRACKS_METRICS_2024)
anova(ba_tort_model7_24i,ba_tort_model7_24h)
summary(ba_tort_model7_24i)
ba_tort_model7_24j <- lmer(mean_angle_3D ~ 1+ (1|Family)+(1|LatinName), data=MULTITRACKS_METRICS_2024)
anova(ba_tort_model7_24i,ba_tort_model7_24j)# same result
AIC(ba_tort_model7_24i ,ba_tort_model10_24i )# model fit better with family and species

#model for 2025
MULTITRACKS_METRICS_2025<-MULTITRACKS_SEGMENTS%>%
  filter(Year=="2025")%>%
  droplevels()
#go straight to testing impacts of light structure and background light level
ba_tort_model1_25 <- lmer(mean_angle_3D ~ BgdLightLevel*LightStructure*After_Light_Both_Exposed+TrialNumber+CaptureMethod+ (1|Unique_Flight_ID)+(1|Family)+(1|LatinName), data=MULTITRACKS_METRICS_2025)
  #here, can fit all random effects
summary(ba_tort_model1_25)
ba_tort_model1_25_resi <- simulateResiduals(ba_tort_model1_25)
plot(ba_tort_model1_25_resi)#ok
ba_tort_model1_25b <- lmer(mean_angle_3D  ~ BgdLightLevel*LightStructure +LightStructure*After_Light_Both_Exposed+BgdLightLevel*After_Light_Both_Exposed+TrialNumber+CaptureMethod+ (1|Unique_Flight_ID)+(1|Family)+(1|LatinName), data=MULTITRACKS_METRICS_2025)
anova(ba_tort_model1_25b,ba_tort_model1_25) #remove three-way interaction
summary(ba_tort_model1_25b)
ba_tort_model1_25c <- lmer(mean_angle_3D  ~ BgdLightLevel*LightStructure +LightStructure*After_Light_Both_Exposed+BgdLightLevel*After_Light_Both_Exposed+CaptureMethod+(1|Unique_Flight_ID)+(1|Family)+(1|LatinName), data=MULTITRACKS_METRICS_2025)
anova(ba_tort_model1_25b,ba_tort_model1_25c) #remove trial number
summary(ba_tort_model1_25c)
ba_tort_model1_25d <- lmer(mean_angle_3D  ~ LightStructure*After_Light_Both_Exposed+BgdLightLevel*After_Light_Both_Exposed+CaptureMethod+(1|Unique_Flight_ID)+(1|Family)+(1|LatinName), data=MULTITRACKS_METRICS_2025)
anova(ba_tort_model1_25d,ba_tort_model1_25c) #remove bgd:structure interaction
summary(ba_tort_model1_25d)
ba_tort_model1_25e <- lmer(mean_angle_3D  ~ LightStructure*After_Light_Both_Exposed+BgdLightLevel*After_Light_Both_Exposed+(1|Unique_Flight_ID)+(1|Family)+(1|LatinName), data=MULTITRACKS_METRICS_2025)
anova(ba_tort_model1_25d,ba_tort_model1_25e) #remove capture method
summary(ba_tort_model1_25e)
ba_tort_model1_25f <- lmer(mean_angle_3D  ~ LightStructure+After_Light_Both_Exposed+BgdLightLevel*After_Light_Both_Exposed+(1|Unique_Flight_ID)+(1|Family)+(1|LatinName), data=MULTITRACKS_METRICS_2025)
anova(ba_tort_model1_25f,ba_tort_model1_25e) #remove structure:exposure interaction
summary(ba_tort_model1_25f)
ba_tort_model1_25g <- lmer(mean_angle_3D ~ LightStructure+After_Light_Both_Exposed+BgdLightLevel+(1|Unique_Flight_ID)+(1|Family)+(1|LatinName), data=MULTITRACKS_METRICS_2025)
anova(ba_tort_model1_25f,ba_tort_model1_25g) #cannot remove bgd:exposure interaction
ba_tort_model1_25h <- lmer(mean_angle_3D ~ After_Light_Both_Exposed*BgdLightLevel+(1|Unique_Flight_ID)+(1|Family)+(1|LatinName), data=MULTITRACKS_METRICS_2025)
anova(ba_tort_model1_25f,ba_tort_model1_25h) #cannot remove structure
summary(ba_tort_model1_25f)
emmeans(ba_tort_model1_25f, pairwise ~ After_Light_Both_Exposed | BgdLightLevel) #p values per treatment group


#plot results for 2025
plot_tortuosity_25_level <- ggplot(MULTITRACKS_METRICS_2025, aes(x = BgdLightLevel, y = mean_angle_3D, fill = After_Light_Both_Exposed,col = After_Light_Both_Exposed)) +
  geom_boxplot() +
  scale_color_manual(values=c("#19a4fa","grey40"))+scale_fill_manual(values=c("white","#19a4fa"))+
  theme_classic() +
  #guides(fill="none", col="none")+
  labs(title = "F", y = "Mean turning angle (degrees)", x = "Background light level", fill="Light exposure", col="Light exposure") +
  annotate(
    "text",
    x = c(1, 2),    
    y = c(60,60), 
    label = c("***","**"),  
    size = 4 ) +
  theme(
    axis.text = element_text(size = 10),
    axis.title = element_text(size = 12),
    plot.title = element_text(size = 16),
    panel.grid.major = element_blank(),
    panel.grid.minor = element_blank(),
    panel.border = element_blank(),
    axis.line = element_line(color = "black")
  )
plot_tortuosity_25_level 

plot_tortuosity_25_structure <- ggplot(MULTITRACKS_METRICS_2025, aes(x = LightStructure, y = mean_angle_3D)) +
  geom_boxplot_pattern(aes(fill=LightStructure,pattern_spacing=LightStructure), 
                       pattern="circle", pattern_fill="#19a4fa",colour="#19a4fa", fill="white") +
  theme_classic() +
  scale_pattern_spacing_manual(values = c(0.1, 0.05))+
  annotate(
    "text",
    x = c(1.5), 
    y = c(60),  
    label = c("*"),  
    size = 6 ) +
  labs(title = "G", x = "Focal light structure", fill="", y="Mean turning angle (degrees)") +
  theme(
    axis.text = element_text(size = 10),
    axis.title = element_text(size = 12),
    plot.title = element_text(size = 16),
    panel.grid.major = element_blank(),
    panel.grid.minor = element_blank(),
    panel.border = element_blank(), 
    axis.line = element_line(color = "black")  
  )+guides(fill="none", pattern_spacing="none")
plot_tortuosity_25_structure

#statistics on flight segments - sd----

ba_sd_multi_mod1 <- lmer(sd_angle_3D ~ BgdLightLevel*LightStructure*After_Light_Both_Exposed+TrialNumber+CaptureMethod+ (1|Unique_Flight_ID)+(1|Family)+(1|LatinName), data=MULTITRACKS_SEGMENTS)
summary(ba_sd_multi_mod1 )
ba_sd_multi_mod1_resi <- simulateResiduals(ba_sd_multi_mod1 )
plot(ba_sd_multi_mod1_resi)#poor
hist(resid(ba_sd_multi_mod1 ))
ba_sd_multi_mod2 <- lmer(sqrt(sd_angle_3D) ~ BgdLightLevel*LightStructure*After_Light_Both_Exposed+TrialNumber+CaptureMethod+ (1|Unique_Flight_ID)+ (1|Family)+(1|LatinName), data=MULTITRACKS_SEGMENTS)
summary(ba_sd_multi_mod2)
ba_sd_multi_mod2_resi <- simulateResiduals(ba_sd_multi_mod2)
plot(ba_sd_multi_mod2_resi)#not perfect but better, a log transform is worse
hist(resid(ba_sd_multi_mod2))
ba_sd_multi_mod2b <- lmer(sqrt(sd_angle_3D) ~ BgdLightLevel*LightStructure +LightStructure*After_Light_Both_Exposed+BgdLightLevel*After_Light_Both_Exposed+TrialNumber+CaptureMethod+ (1|Unique_Flight_ID)+(1|Family)+(1|LatinName), data=MULTITRACKS_SEGMENTS)
anova(ba_sd_multi_mod2b,ba_sd_multi_mod2) #remove three-way interaction
summary(ba_sd_multi_mod2b)
ba_sd_multi_mod2c <- lmer(sqrt(sd_angle_3D) ~ BgdLightLevel*LightStructure +LightStructure*After_Light_Both_Exposed+BgdLightLevel*After_Light_Both_Exposed+CaptureMethod+ (1|Unique_Flight_ID)+(1|Family)+(1|LatinName), data=MULTITRACKS_SEGMENTS)
anova(ba_sd_multi_mod2b,ba_sd_multi_mod2c) #remove trial number
summary(ba_sd_multi_mod2c)
ba_sd_multi_mod2d <- lmer(sqrt(sd_angle_3D) ~ LightStructure*After_Light_Both_Exposed+BgdLightLevel*After_Light_Both_Exposed+CaptureMethod+ (1|Unique_Flight_ID)+(1|Family)+(1|LatinName), data=MULTITRACKS_SEGMENTS)
anova(ba_sd_multi_mod2d,ba_sd_multi_mod2c) #remove bgd:structure interaction
summary(ba_sd_multi_mod2d)
ba_sd_multi_mod2e <- lmer(sqrt(sd_angle_3D) ~ LightStructure*After_Light_Both_Exposed+BgdLightLevel*After_Light_Both_Exposed+ (1|Unique_Flight_ID)+(1|Family)+(1|LatinName), data=MULTITRACKS_SEGMENTS)
anova(ba_sd_multi_mod2d,ba_sd_multi_mod2e) #remove capture method
summary(ba_sd_multi_mod2e)
ba_sd_multi_mod2f <- lmer(sqrt(sd_angle_3D) ~ LightStructure+After_Light_Both_Exposed+BgdLightLevel*After_Light_Both_Exposed+ (1|Unique_Flight_ID)+(1|Family)+(1|LatinName), data=MULTITRACKS_SEGMENTS)
anova(ba_sd_multi_mod2f,ba_sd_multi_mod2e) #remove structure:exposure interaction
summary(ba_sd_multi_mod2f)

ba_sd_multi_mod2g <- lmer(sqrt(sd_angle_3D) ~ LightStructure+After_Light_Both_Exposed+BgdLightLevel+ (1|Unique_Flight_ID)+(1|Family)+(1|LatinName), data=MULTITRACKS_SEGMENTS)
anova(ba_sd_multi_mod2f,ba_sd_multi_mod2g) #cannot remove bgd:exposure interaction
ba_sd_multi_mod2h <- lmer(sqrt(sd_angle_3D) ~ After_Light_Both_Exposed*BgdLightLevel+ (1|Unique_Flight_ID)+(1|Family)+(1|LatinName), data=MULTITRACKS_SEGMENTS)
anova(ba_sd_multi_mod2f,ba_sd_multi_mod2h) #cannot remove structure

emmeans(ba_sd_multi_mod2f, pairwise ~ After_Light_Both_Exposed | BgdLightLevel) #p values per treatment group

#plot for sd
plot_sd_level <- ggplot(MULTITRACKS_SEGMENTS, aes(x = BgdLightLevel, y = sqrt(sd_angle_3D), fill = After_Light_Both_Exposed,col = After_Light_Both_Exposed)) +
  geom_boxplot() +
  scale_color_manual(values=c("#19a4fa","grey40"))+scale_fill_manual(values=c("white","#19a4fa"))+
  theme_classic() +
  #guides(fill="none", col="none")+
  labs(title = "F", y = "sqrt (SD turning angle (degrees))", x = "Background light level", fill="Light exposure", col="Light exposure") +
  annotate(
    "text",
    x = c(1, 2,3),    
    y = c(9,9,9), 
    label = c("***","***","**"),  
    size = 4 ) +
  theme(
    axis.text = element_text(size = 10),
    axis.title = element_text(size = 12),
    plot.title = element_text(size = 16),
    panel.grid.major = element_blank(),
    panel.grid.minor = element_blank(),
    panel.border = element_blank(),
    axis.line = element_line(color = "black")
  )
plot_sd_level 

plot_sd_structure <- ggplot(MULTITRACKS_METRICS, aes(x = LightStructure, y = sqrt(sd_angle_3D))) +
  geom_boxplot_pattern(aes(fill=LightStructure,pattern_spacing=LightStructure), 
                       pattern="circle", pattern_fill="#19a4fa",colour="#19a4fa", fill="white") +
  theme_classic() +
  scale_pattern_spacing_manual(values = c(0.1, 0.05))+
  annotate(
    "text",
    x = c(1.5), 
    y = c(9),  
    label = c("***"),  
    size = 6 ) +
  labs(title = "G", x = "Focal light structure", fill="", y="sqrt (SD turning angle (degrees))") +
  theme(
    axis.text = element_text(size = 10),
    axis.title = element_text(size = 12),
    plot.title = element_text(size = 16),
    panel.grid.major = element_blank(),
    panel.grid.minor = element_blank(),
    panel.border = element_blank(), 
    axis.line = element_line(color = "black")  
  )+guides(fill="none", pattern_spacing="none")
plot_sd_structure

#Plot all results for flight tortuosity (Figure 4)----
layout <- "
####B
CCCC#
EEFFG
"
plot_sp_expt1_intensity+plot_tortuosity_ba+plot_sp_expt2b+
  plot_tortuosity_25_level+plot_tortuosity_25_structure+plot_layout(design=layout)


#SUPPLEMENTARY ANALYSES -----

#Expt 1 : dark control vs control with 3 lx background light----
str(SUPP_SINGLE)
  #can only look at take-off probability - only 2 flights in control 3 lx group meet criteria for outcome/tortuosity analysis
  #supplementary dataset includes control flights from 2025 with low background light level, and 3lx (high) background light level

#create new variable to represent background light levels:
SUPP_SINGLE<-SUPP_SINGLE%>%
  dplyr::mutate(Level = case_when(
    is.na(Intensity) ~ "Low",
    Intensity =="3lx" ~"High"))%>%
  filter(LatinName!="Unknown")#remove trials with unknown species
    
SUPP_SINGLE%>%
  group_by(TakeOff, Level)%>%
  summarise(n=n()) 
  #109 flight attempts in regular controls in 2025, with 37 take-offs (34%), 
  #24 flights under 3lx bgd light, of which only 3 had moths taking off (13%) 

#Plot proportion of successful take-offs - figure S1

SUPP_SINGLE$Level<-factor(SUPP_SINGLE$Level, levels=c("Low","High"))
ggplot(data = SUPP_SINGLE) +
  geom_mosaic(aes(x = product(TakeOff, Level), fill = TakeOff), col="mediumpurple1")+
  labs(fill="Take-off")+
  scale_fill_manual(values = c("white","mediumpurple1"))+
  ylab("Proportion of trials")+
  xlab("Background light level")+
  geom_mosaic_text(aes(x = product(TakeOff, Level),label = after_stat(.wt)), col="grey30")+
  theme(legend.text = element_text(size=10))+
  theme_classic()+
  theme(axis.title.x = element_text(size = 12),
        axis.text.x = element_text(size = 10),
        axis.title.y = element_text(size = 12),
        axis.text.y=element_text(size=10))

#probability of take-off between treatments
str(SUPP_SINGLE)
supp_mod1<-glmmTMB(TakeOff ~ Level+CaptureMethod+
                     (1|LatinName)+(1|Family),
                   family=binomial, data=SUPP_SINGLE)
summary(supp_mod1)
supp_mod1_resid <- simulateResiduals(supp_mod1)
plot(supp_mod1_resid)#good

supp_mod1b<-glmmTMB(TakeOff ~Level+
                      (1|LatinName)+(1|Family),
                    family=binomial, data=SUPP_SINGLE)
anova(supp_mod1, supp_mod1b)#no effect of capture method

supp_mod1c<-glmmTMB(TakeOff ~ 1+(1|LatinName)+(1|Family),
                    family=binomial, data=SUPP_SINGLE)
anova(supp_mod1c, supp_mod1b)#significant effect of background light level
summary(supp_mod1b)

#odds ratios
exp(-1.354)#odds ratio 0.258, odds of taking off are reduced by three quarters under high background light levels

#Expt 1: understanding effects of capture method----
  #additional analyses to explore whether capture method effects are due to biased sampling

#1- Distribution of species and families in trap/net samples

#identify species caught with either method, or both

TRAP_ONLY<-INFO_SINGLE%>%
  filter(CaptureMethod =="Trap")%>%
  group_by(LatinName)%>%
  summarise(n=n())%>%
  droplevels()
NET_ONLY<-INFO_SINGLE%>%
  filter(CaptureMethod =="Net")%>%
  group_by(LatinName)%>%
  summarise(n=n())%>%
  droplevels()
trapspecies<-unique(TRAP_ONLY$LatinName)#48
netspecies<-unique(NET_ONLY$LatinName)#37

intersect(trapspecies, netspecies)#30 species collected using both methods

#families

FAMILIES_METHOD<-INFO_SINGLE%>%
  filter(!is.na(TakeOff))%>%
  group_by(Family, CaptureMethod, TakeOff)%>%
  summarise(n=n())
flightlabs<-c("no flight", "flight")
names(flightlabs)<-c(0,1)
species_a<-ggplot(FAMILIES_METHOD, aes(fill=Family, y=n, x=CaptureMethod, label=n)) + 
  geom_bar(position="fill", stat="identity")+
  facet_wrap(.~TakeOff, labeller = labeller(TakeOff = flightlabs))+
  theme_classic()+ylab("Proportion of trials")+
  scale_y_continuous(expand=c(0,0))+
  ylab("Capture method")+
  ggtitle("A")+
  geom_text(size = 3, position = position_fill(vjust = 0.5))+
  theme(axis.title.x = element_text(size = 14),
        axis.text.x = element_text(size = 12),
        axis.title.y = element_text(size = 14),
        axis.text.y=element_text(size=12))+
  theme(strip.text.x = element_text(size = 12))
  #composition is fairly similar between methods, and between successful and unsuccessful trials
species_a

#repeat analyses with species restricted to those caught in good numbers with both methods

SPECIESTEST<-INFO_SINGLE%>%
  group_by(LatinName, CaptureMethod)%>%
  summarise(n=n()) %>%
  filter(n>=10)%>%#at least 10 from a given method
  group_by(LatinName)%>%
  summarise(n2=n())%>%
  filter(n2==2)#keep only species with over 10 in both methods
#this yields 9 species - 2 erebids, 1 crambid, 3 noctuids, 3 geometrids

#run takeoff probability model for these only
  #treatment not included as it has no effect in main model
TEST<-INFO_SINGLE%>%
  filter(LatinName %in% SPECIESTEST$LatinName)
test_mod1<-glmmTMB(TakeOff ~ CaptureMethod+
                     (1|LatinName)+(1|Family),
                   family=binomial, data=TEST)
summary(test_mod1)
test_mod1_resid <- simulateResiduals(test_mod1)
plot(test_mod1_resid)#good
test_mod1b<-glmmTMB(TakeOff ~ 1+
                     (1|LatinName)+(1|Family),
                   family=binomial, data=TEST)
anova(test_mod1, test_mod1b) #still a significant effect of capture method
exp(-0.597)#moths half as likely to take off if caught with traps

#test for an interaction with species
test_mod2<-glmmTMB(TakeOff ~ CaptureMethod*LatinName+(1|Family),
                   family=binomial, data=TEST)
summary(test_mod2)
test_mod2_resid <- simulateResiduals(test_mod2)
plot(test_mod2_resid)#good
test_mod2b<-glmmTMB(TakeOff ~ CaptureMethod+LatinName+
                      (1|LatinName)+(1|Family),
                    family=binomial, data=TEST)
anova(test_mod2, test_mod2b) #interaction not significant
summary(test_mod2b)#species-level differences and overall effect of capture method


#plot
SUMMARY_TEST<-TEST%>%
  filter(!is.na(TakeOff))%>%
  group_by(CaptureMethod, TakeOff, LatinName)%>%
  summarise(n=n())%>%
  ungroup()
SUMMARY_TEST$TakeOff<-as.factor(SUMMARY_TEST$TakeOff)
SUMMARY_TEST$MethodTO<-paste(SUMMARY_TEST$CaptureMethod,SUMMARY_TEST$TakeOff)#plotting variable
testcolours<-c("darkgreen","white", "palegreen","white")

species_b<-ggplot(SUMMARY_TEST, aes(x=CaptureMethod, y=n,fill=MethodTO,col=CaptureMethod,label=n)) + 
  geom_bar(position="fill", stat="identity")+
  facet_wrap(.~LatinName)+
  ggtitle("B")+
  scale_fill_manual(values = c("white", "darkgreen","white","palegreen"))+
  scale_color_manual(values=c("darkgreen","palegreen"))+
  theme_classic()+xlab("Capture method")+ylab("Proportion of trials with flights")+
  geom_text(size = 3, position = position_fill(vjust = 0.5), col="grey70")+
  guides(fill="none", col="none")+
  theme(strip.text.x = element_text(size = 12,  face = "italic"))+
  theme(axis.title.x = element_text(size = 14),
        axis.title= element_text(size = 16),
      axis.text.x = element_text(size = 12),
      axis.title.y = element_text(size = 14),
      axis.text.y=element_text(size=12))
species_b

#plot model results
species_c<-plot_model(test_mod1, type = "pred", 
           terms = c("CaptureMethod"),
           title = "C",
           colors = "darkgreen")+
  theme_classic()+
  scale_y_continuous(limits=c(0.1,0.8),breaks = c(0.2, 0.4, 0.6,0.8), labels = c(20,40,60,80))+  
  ylab("Predicted flight probability (%)")+xlab("Capture method")+
  theme(axis.title.x = element_text(size = 14),
        axis.title = element_text(size=16),
        axis.text.x = element_text(size = 12),
        axis.title.y = element_text(size = 14),
        axis.text.y=element_text(size=12))

#re-analyse flight-to-light the same way

OUTCOMES_TEST<-OUTCOMES_EXPT1_FTL%>%
  filter(LatinName %in% SPECIESTEST$LatinName)

OUTCOMES_TEST%>%
  group_by(Treatment)%>%
  summarise(n=n())
ftl_testmod1<-glmmTMB(FlighttoLight_tracked ~ Spectrum+Intensity+CaptureMethod+
                    (1|LatinName),#best model here; family leads to non-convergence, using treatment as fixed effect to very high error
                  family=binomial, data=OUTCOMES_TEST)
summary(ftl_testmod1)
ftl_testmod1_resid <- simulateResiduals(ftl_testmod1)
plot(ftl_testmod1_resid)#ok
ftl_testmod1b<-glmmTMB(FlighttoLight_tracked ~ Intensity+CaptureMethod+
                     (1|LatinName),
                   family=binomial, data=OUTCOMES_TEST)
anova(ftl_testmod1, ftl_testmod1b)#no effect of spectrum
ftl_testmod1c<-glmmTMB(FlighttoLight_tracked ~ Intensity+
                     (1|LatinName),
                   family=binomial, data=OUTCOMES_TEST)
anova(ftl_testmod1c, ftl_testmod1b)#no effect of capture method (p = 0.0856)
summary(ftl_testmod1b)

#plot
SUMMARY_OUT<-OUTCOMES_TEST%>%
  filter(!is.na(TakeOff))%>%
  group_by(CaptureMethod, FlighttoLight_tracked, LatinName)%>%
  summarise(n=n())%>%
  ungroup()
SUMMARY_OUT$FlighttoLight_tracked<-as.factor(SUMMARY_OUT$FlighttoLight_tracked)
SUMMARY_OUT$MethodFTL<-paste(SUMMARY_OUT$CaptureMethod,SUMMARY_OUT$FlighttoLight_tracked)
species_d<-ggplot(SUMMARY_OUT, aes(x=CaptureMethod, y=n, fill=MethodFTL,col=CaptureMethod,label=n)) + 
  geom_bar(position="fill", stat="identity")+
  facet_wrap(.~LatinName)+
  ggtitle("D")+
  scale_fill_manual(values = c("white", "darkgreen","white","palegreen"))+
  scale_color_manual(values=c("darkgreen","palegreen"))+
  theme_classic()+xlab("Background light level")+ylab("Proportion of flights ending at light")+
  geom_text(size = 3, position = position_fill(vjust = 0.5), col="grey70")+
  guides(fill="none", col="none")+
  theme(strip.text.x = element_text(size = 12,  face = "italic"))+
  theme(axis.title.x = element_text(size = 14),
        axis.text.x = element_text(size = 12),
        axis.title.y = element_text(size = 14),
        axis.text.y=element_text(size=12))
species_d

#plot together for Figure S2
species_a+species_b+species_c+species_d+plot_layout(widths =c(1,2))

#Expt 2: effect of single light at 10lx----

str(SUPP_MULTI)
#restrict to all trials for the moths that experienced single 10lx central light
  #this is useful for testing effect of multiple lights compared to a single one
  #same year, same background light level, same individual moths
  #treatments are single 10lx versus 3 lights at 10 lx
SUPP_MULTI_PAIRS<-SUPP_MULTI%>%
  filter(Unique_Moth_ID %in% INFO_SINGLE10LX$Unique_Moth_ID)%>%
  droplevels()
str(SUPP_MULTI_PAIRS)#94 trials, 47 moths
levels(SUPP_MULTI$LatinName)#no unknown species here

#restrict to trials under single 10lx central light OR single 30lx central light
  #this is useful for testing the effect of focal light intensity alone 
  #should replicate findings of experiment 1 
  #same year and background light level, but different moths between groups (no need for ID random effect)
SUPP_MULTI_CENTRAL<-SUPP_MULTI%>%
  filter(LightStructure!="Multiple")%>%
  droplevels()
str(SUPP_MULTI_CENTRAL)#130 trials
length(unique(SUPP_MULTI_CENTRAL$Unique_Moth_ID))#130 unique moths

#Part 1: 1 light at 10lx vs 3----

#1)Take-off probability

#plots
SUMMARY_SUPP_PAIRS<-SUPP_MULTI_PAIRS%>%
  filter(!is.na(TakeOff))%>%#exclude NAs
  group_by(LightStructure,TakeOff)%>%
  summarise(n=n())%>%
  ungroup()%>%
  mutate(TakeOff = factor(TakeOff,
                          levels = c("0","1")))%>%
  mutate(LightStructure = factor(LightStructure,
                        levels = c("Single 10lx","Multiple")))
supp_to<-ggplot(SUMMARY_SUPP_PAIRS, aes(fill=TakeOff, y=n, x=LightStructure, label=n)) + 
  geom_bar(position="fill", stat="identity", col="lightskyblue1")+
  ylab("Proportion of trials")+
  labs(fill="Take-off")+
  ggtitle("A")+
  scale_x_discrete(name = "Number of 10 lx focal lights",labels = c("One", "Three"
  ))+
  scale_fill_manual(values = c("white","lightskyblue1"),labels=c('no flight', 'flight'))+
  geom_text(size = 3, position = position_fill(vjust = 0.5), col="grey40")+
  theme(legend.text = element_text(size=10),
        legend.title = element_blank())+
  theme_classic()+
  theme(axis.title.x = element_text(size = 12),
        axis.text.x = element_text(size=10),
        axis.title.y = element_text(size = 12),
        axis.text.y=element_text(size=10),
        axis.title=element_text(size=14))+
  scale_y_continuous(expand=c(0,0))
supp_to

#probability of take-off between treatments
SUPP_MULTI_PAIRS$LightStructure<-factor(SUPP_MULTI_PAIRS$LightStructure, levels=c("Single 10lx","Multiple"))
pairs_mod1<-glmmTMB(TakeOff ~ LightStructure+CaptureMethod+TrialNumber+(1|Unique_Moth_ID)+
                     (1|LatinName)+(1|Family),
                   family=binomial, data=SUPP_MULTI_PAIRS)
summary(pairs_mod1)
pairs_mod1_resid <- simulateResiduals(pairs_mod1)
plot(pairs_mod1_resid)#good

pairs_mod1b<-glmmTMB(TakeOff ~ LightStructure+TrialNumber+(1|Unique_Moth_ID)+
                      (1|LatinName)+(1|Family),
                    family=binomial, data=SUPP_MULTI_PAIRS)
anova(pairs_mod1, pairs_mod1b)#no effect of capture method

pairs_mod1c<-glmmTMB(TakeOff ~ LightStructure+(1|Unique_Moth_ID)+(1|LatinName)+(1|Family),
                    family=binomial, data=SUPP_MULTI_PAIRS)
anova(pairs_mod1c, pairs_mod1b)#no effect of trial number
summary(pairs_mod1c)
pairs_mod1d<-glmmTMB(TakeOff ~ 1+(1|Unique_Moth_ID)+(1|LatinName)+(1|Family),
                     family=binomial, data=SUPP_MULTI_PAIRS)
anova(pairs_mod1c, pairs_mod1d)#no effect of treatment

#2)Flight-to-light

#keep only outcome information for flights that meet analysis criteria, in the two paired groups
OUTCOMES_PAIRS_MULTI<-SUPP_MULTI_PAIRS%>%
  filter(Unique_Flight_ID %in% SUPP_MULTITRACKS$Unique_Flight_ID)%>%
  droplevels()
dim(OUTCOMES_PAIRS_MULTI)#30 flights, out of 48 that took off
str(OUTCOMES_PAIRS_MULTI)

OUTCOMES_PAIRS_MULTI$Outcome_simpler<-factor(OUTCOMES_PAIRS_MULTI$Outcome_simpler,
                                       levels=c("at light","up","down","other"))

#overall proportion of flights ending at the light:
OUTCOMES_PAIRS_MULTI%>%
  group_by(FlighttoLight_tracked)%>%
  summarise(n=n())%>%
  ungroup()
8/30#27% flights ended at the light

#plot
SUMMARY_OUTCOME_PAIRS_MULTI<-OUTCOMES_PAIRS_MULTI%>%
  group_by(LightStructure, Outcome_simpler, Year)%>%
  summarise(n=n())%>%
  ungroup()
SUMMARY_OUTCOME_PAIRS_MULTI$Outcome_simpler<-factor(SUMMARY_OUTCOME_PAIRS_MULTI$Outcome_simpler,
                                              levels=c("other","down","up","at light"))

supp_b<-ggplot(SUMMARY_OUTCOME_PAIRS_MULTI, aes(fill=Outcome_simpler, y=n, x=LightStructure, label=n)) + 
  geom_bar(position="fill", stat="identity")+
  theme_bw()+ylab("Proportion of flights")+
  geom_text(size = 3, position = position_fill(vjust = 0.5))+
  labs(fill="Flight outcome")+
  xlab("Number of 10lx focal lights")+
  scale_x_discrete(labels=c("One", "Three"))+
  scale_fill_manual(values = c("#B3B3B3","#66C2A5","#E78AC3","#FFD92F"))+
  geom_text(size = 3, position = position_fill(vjust = 0.5), col="grey30")+  theme(legend.text = element_text(size=10))+
  theme_classic()+
  ggtitle("B")+
  theme(axis.title.x = element_text(size = 12),
        axis.title = element_text(size = 14),
        axis.text.x = element_text(size = 10),
        axis.title.y = element_text(size = 12),
        axis.text.y=element_text(size=10))+
  #scale_x_discrete(drop = TRUE) +
  scale_y_continuous(expand=c(0,0))
supp_b
#analysis - for flight-to-light only

pairs_mod2<-glmmTMB(FlighttoLight_tracked ~ LightStructure+TrialNumber+CaptureMethod+
                    (1|Unique_Moth_ID)+(1|LatinName)+(1|Family),
                  family=binomial, data=OUTCOMES_PAIRS_MULTI)
summary(pairs_mod2)
pairs_mod2_resid <- simulateResiduals(pairs_mod2)
plot(pairs_mod2_resid)#ok

pairs_mod2b<-glmmTMB(FlighttoLight_tracked ~ LightStructure+TrialNumber+
                     (1|Unique_Moth_ID)+(1|LatinName)+(1|Family),
                   family=binomial, data=OUTCOMES_PAIRS_MULTI)

anova(pairs_mod2, pairs_mod2b)#remove capture method
summary(pairs_mod2b)
pairs_mod2c<-glmmTMB(FlighttoLight_tracked ~ LightStructure+
                       (1|Unique_Moth_ID)+(1|LatinName)+(1|Family),
                     family=binomial, data=OUTCOMES_PAIRS_MULTI)

anova(pairs_mod2c, pairs_mod2b)#remove trial number
pairs_mod2d<-glmmTMB(FlighttoLight_tracked ~ 1+
                       (1|Unique_Moth_ID)+(1|LatinName)+(1|Family),
                     family=binomial, data=OUTCOMES_PAIRS_MULTI)

anova(pairs_mod2c, pairs_mod2d)#no effect of light structure either

#3)Erratic spiraling

OUTCOMES_PAIRS_MULTI%>%
  group_by(Erratic_spiralling)%>%
  summarise(n=n())%>%
  ungroup()
5/30 #erratic spiraling in 16 % flights overall

SPIRAL_SUMMARY_PAIRS<-OUTCOMES_PAIRS_MULTI%>%
  group_by(Erratic_spiralling, LightStructure)%>%
  summarise(n=n())%>%
  ungroup()

#alternative as barplot
SPIRAL_SUMMARY_PAIRS$Erratic_spiralling<-factor(SPIRAL_SUMMARY_PAIRS$Erratic_spiralling, levels=c("0","1"))
supp_c<-ggplot(SPIRAL_SUMMARY_PAIRS, aes(fill=Erratic_spiralling, y=n, x=LightStructure, label=n)) + 
  geom_bar(position="fill", stat="identity", col="lightskyblue1")+
  ylab("Proportion of trials")+
  labs(fill="Erratic spiraling")+
  ggtitle("C")+
  xlab("Number of 10 lx focal lights")+
  scale_x_discrete(labels = c("One","Three"))+
  scale_fill_manual(values = c("white","lightskyblue1"), labels =c("no", "yes"))+
  geom_text(size = 3, position = position_fill(vjust = 0.5), col="grey30")+
  theme(legend.text = element_text(size=10))+
  theme_classic()+
  theme(axis.title.x = element_text(size = 12),
        axis.title = element_text(size = 14),
        axis.text.x = element_text(size = 10),
        axis.title.y = element_text(size = 12),
        axis.text.y=element_text(size=10))+
  scale_y_continuous(expand=c(0,0))
supp_c
#2 or 3 flights with erratic spiraling depending on treatment - not enough variation to analyse 

#4)Tortuosity

#calculate tortuosity pre and post exposure, for all supplementary tracks, then filter out only those relevant to this analysis

#apply turning angle function to each unique flight i
suppFlightIDs<-unique(SUPP_MULTITRACKS$Unique_Flight_ID)
suppFLIGHTS<-list()

for (i in 1:length(suppFlightIDs)){
  TEMPDF<-SUPP_MULTITRACKS%>%
    filter(Unique_Flight_ID ==suppFlightIDs[i])
  str(TEMPDF)
  suppFLIGHTS[[i]]<-calculate_turning_angles_and_lengths(TEMPDF)#create new list with data with turning angles for each flight
}

#remove any points with large displacements
suppFLIGHTS<-lapply(suppFLIGHTS, function(x) filter(x, Displacement < 0.5))


#combine into larger dataframe
SUPP_MULTITRACKS_ANGLES <- bind_rows(suppFLIGHTS)

#create variable to signify whether or not (0 no and 1 yes) the moth is exposed by facing the light 
SUPP_MULTITRACKS_ANGLES$Exposed <- ifelse(SUPP_MULTITRACKS_ANGLES$bearing_to_light_deg <= 135, 1, 0)

#combine with position after light cone to create variable for when moth has both passed the light and has been exposed to it
SUPP_MULTITRACKS_ANGLES$After_Light_Both_Exposed <- ifelse(
  !is.na(SUPP_MULTITRACKS_ANGLES$Exposed) & 
    !is.na(SUPP_MULTITRACKS_ANGLES$After_Light_Both) & 
    SUPP_MULTITRACKS_ANGLES$Exposed == 1 & SUPP_MULTITRACKS_ANGLES$After_Light_Both == 1,
  1,
  0
)

SUPP_MULTITRACKS_ANGLES <- SUPP_MULTITRACKS_ANGLES %>%
  arrange(Unique_Flight_ID, Frame) %>%# sort chronologically within each track
  group_by(Unique_Flight_ID) %>%
  mutate(
    After_Light_Both_Exposed = if_else(
      cumsum(After_Light_Both_Exposed) > 0, 
      1,
      0
    )
  ) %>%
  ungroup()

#create variable to separate flights before and after passing through light cone
SUPP_MULTITRACKS_ANGLES$UniqueSegment<-paste(SUPP_MULTITRACKS_ANGLES$Unique_Flight_ID,SUPP_MULTITRACKS_ANGLES$After_Light_Both_Exposed, sep="_")
str(SUPP_MULTITRACKS_ANGLES)

#identify flights with at least 5 points before and after cone (in both cameras)
SUPP_WITHBOTH<-SUPP_MULTITRACKS_ANGLES%>%
  group_by(Unique_Flight_ID,After_Light_Both_Exposed)%>%
  summarise(n=n())%>%
  filter(n>=5)%>%
  group_by(Unique_Flight_ID)%>%
  summarise(n=n())%>%
  filter(n==2)
unique(SUPP_WITHBOTH$Unique_Flight_ID)#82 flights out of 93 in supp_multitracks

#restrict track data to only those flights
SUPP_TRACKS_BEFOREAFTER<-SUPP_MULTITRACKS_ANGLES%>%
  filter(Unique_Flight_ID %in% SUPP_WITHBOTH$Unique_Flight_ID)

#apply function to calculate tortuosity for these segments

#split by unique segment IDs - turning angles etc already calculated
SUPP_segmentIDs<-unique(SUPP_TRACKS_BEFOREAFTER$UniqueSegment)
SUPP_SEGMENTS<-list()

for (i in 1:length(SUPP_segmentIDs)){
  SUPP_SEGMENTS[[i]]<-SUPP_TRACKS_BEFOREAFTER%>%
    filter(UniqueSegment == SUPP_segmentIDs[i])
}

# apply tortuosity function to each dataframe in the results_list
SUPP_MULTI_TORTUOSITY_LIST_SEGMENTS <- lapply(SUPP_SEGMENTS, calculate_tortuosity_metrics)
length(SUPP_MULTI_TORTUOSITY_LIST_SEGMENTS)#164 - 2 segments each for 82 flights
head(SUPP_MULTI_TORTUOSITY_LIST_SEGMENTS[[1]])

# Combine the results into a single dataframe
SUPP_MULTITRACKS_SEGMENTS <- do.call(rbind, SUPP_MULTI_TORTUOSITY_LIST_SEGMENTS)
dim(SUPP_MULTITRACKS_SEGMENTS)

#select only the ones for the pairs dataset
SUPP_PAIRS_SEGMENTS<-SUPP_MULTITRACKS_SEGMENTS%>%
  filter(Unique_Moth_ID %in% INFO_SINGLE10LX$Unique_Moth_ID)%>%
  droplevels()
str(SUPP_PAIRS_SEGMENTS)#27 flights, including 20 different moths

#Plot results
col_names2 <- c("Unique_Moth_ID","UniqueSegment","After_Light_Both_Exposed","Treatment","LatinName","Family","CaptureMethod","Location")
SUPP_PAIRS_SEGMENTS<- SUPP_PAIRS_SEGMENTS %>%
  mutate(across(all_of(col_names2), as.factor))
str(SUPP_PAIRS_SEGMENTS)

treatment_counts <- table(SUPP_PAIRS_SEGMENTS$Treatment)
treatment_counts_vector <- paste0("n=", as.numeric(treatment_counts))
treatment_counts

supp_plot_tortuosity_beforeafter <- ggplot(SUPP_PAIRS_SEGMENTS, aes(x = Treatment, y = mean_angle_3D, fill = After_Light_Both_Exposed)) +
  geom_boxplot() +
  theme_classic() +
  labs(title = "", y = "Mean turning angle (degrees)", x = "Number of 10 lx focal lights") +
  scale_x_discrete(labels = c("One", "Three"))+
  labs(fill="Flight segment")+
  scale_fill_discrete(labels = c("before \nexposure", "after \nexposure"))+
theme(
  
  axis.text = element_text(size = 12),
  axis.title = element_text(size = 14),
  axis.title.y = element_text(size = 12),
  axis.title.x = element_text(size = 12),
  axis.text.x = element_text(size = 10),
  panel.grid.major = element_blank(),
  panel.grid.minor = element_blank(),
  panel.border = element_blank(),  # keep axis lines, so we don't add a full box
  axis.line = element_line(color = "black")  # this adds back the x and y axis lines
) +
  ggtitle("D")
supp_plot_tortuosity_beforeafter #clear effect of exposure, but little difference between treatments otherwise

#statistics
pairs_mod3<- lmer(mean_angle_3D ~ Treatment*After_Light_Both_Exposed+(1|Unique_Flight_ID), data=SUPP_PAIRS_SEGMENTS)
summary(pairs_mod3)#prioritise random effect for flight ID, and remove variables that were not important in main models, to avoid singular fit
pairs_mod3_resi <- simulateResiduals(pairs_mod3)
plot(pairs_mod3_resi)#ok
hist(resid(pairs_mod3))
pairs_mod3b <- lmer(mean_angle_3D ~ Treatment+After_Light_Both_Exposed+(1|Unique_Flight_ID), data=SUPP_PAIRS_SEGMENTS)
anova(pairs_mod3b,pairs_mod3) #remove interaction
summary(pairs_mod3b)
pairs_mod3c <- lmer(mean_angle_3D ~ After_Light_Both_Exposed+(1|Unique_Flight_ID), data=SUPP_PAIRS_SEGMENTS)
anova(pairs_mod3b,pairs_mod3c) #remove treatment
summary(pairs_mod3c)
pairs_mod3d <- lmer(mean_angle_3D ~ 1+(1|Unique_Flight_ID), data=SUPP_PAIRS_SEGMENTS)
anova(pairs_mod3d,pairs_mod3c) #significant effect of exposure to light only

supp_to+supp_b+supp_c+supp_plot_tortuosity_beforeafter+plot_layout(ncol=4)
  #colours adjusted manually posthoc


#Part 2: 1 light at 10lx vs 30 lx----

#1)Take-off probability

#plots

ggplot(data = SUPP_MULTI_CENTRAL) +
  geom_mosaic(aes(x = product(TakeOff, LightStructure), fill = TakeOff), col="#17A3FA")+
  labs(fill="Take-off")+
  scale_fill_manual(values = c("white","#17A3FA"))+
  ylab("Proportion of trials")+
  xlab("Number of lights")+
  geom_mosaic_text(aes(x = product(TakeOff, LightStructure),label = after_stat(.wt)), col="grey30")+
  theme(legend.text = element_text(size=10))+
  theme_classic()+
  theme(axis.title.x = element_text(size = 12),
        axis.text.x = element_text(size = 10),
        axis.title.y = element_text(size = 12),
        axis.text.y=element_text(size=10))

SUMMARY_SUPP_CENTRAL<-SUPP_MULTI_CENTRAL%>%
  filter(!is.na(TakeOff))%>%#exclude NAs
  group_by(Treatment,TakeOff)%>%
  summarise(n=n())%>%
  ungroup()%>%
  mutate(TakeOff = factor(TakeOff,
                          levels = c("0","1")))%>%
  mutate(Treatment = factor(Treatment,
                                 levels = c("SingleC_0.3lx","CL_0.3lx")))
supp_to2<-ggplot(SUMMARY_SUPP_CENTRAL, aes(fill=TakeOff, y=n, x=Treatment, label=n)) + 
  geom_bar(position="fill", stat="identity", col="#17A3FA")+
  ylab("Proportion of trials")+
  labs(fill="Take-off")+
  ggtitle("E")+
  scale_x_discrete(name = "Focal light intensity (lx)",labels = c("10", "30"
  ))+
  scale_fill_manual(values = c("white","#17A3FA"),labels=c('no flight', 'flight'))+
  geom_text(size = 3, position = position_fill(vjust = 0.5), col="grey30")+
  theme(legend.text = element_text(size=10),
        legend.title = element_blank())+
  theme_classic()+
  theme(axis.title.x = element_text(size = 12),
        axis.text.x = element_text(size=10),
        axis.title.y = element_text(size = 12),
        axis.text.y=element_text(size=10),
        axis.title=element_text(size=14))+
  scale_y_continuous(expand=c(0,0))
supp_to2

#probability of take-off between treatments
SUPP_MULTI_CENTRAL$LightStructure<-factor(SUPP_MULTI_CENTRAL$LightStructure, levels=c("Single 10lx","Multiple"))
central_mod1<-glmmTMB(TakeOff ~ Treatment+CaptureMethod+TrialNumber+
                      (1|LatinName)+(1|Family),
                    family=binomial, data=SUPP_MULTI_CENTRAL)
summary(central_mod1)
central_mod1_resid <- simulateResiduals(central_mod1)
plot(central_mod1_resid)#good

central_mod1b<-glmmTMB(TakeOff ~ Treatment+TrialNumber+
                       (1|LatinName)+(1|Family),
                     family=binomial, data=SUPP_MULTI_CENTRAL)
anova(central_mod1, central_mod1b)#no effect of capture method

central_mod1c<-glmmTMB(TakeOff ~ Treatment+(1|LatinName)+(1|Family),
                     family=binomial, data=SUPP_MULTI_CENTRAL)
anova(central_mod1c, central_mod1b)#no effect of trial number
summary(central_mod1c)
central_mod1d<-glmmTMB(TakeOff ~ 1+(1|LatinName)+(1|Family),
                     family=binomial, data=SUPP_MULTI_CENTRAL)
anova(central_mod1c, central_mod1d)#no effect of treatment

#2)Flight-to-light

#keep only outcome information for flights that meet analysis criteria, in the two central groups
OUTCOMES_CENTRAL_MULTI<-SUPP_MULTI_CENTRAL%>%
  filter(Unique_Flight_ID %in% SUPP_MULTITRACKS$Unique_Flight_ID)%>%
  droplevels()
dim(OUTCOMES_CENTRAL_MULTI)#47 flights, out of 57 that took off. 15/47 in single 10lx treatment
str(OUTCOMES_CENTRAL_MULTI)

OUTCOMES_CENTRAL_MULTI$Outcome_simpler<-factor(OUTCOMES_CENTRAL_MULTI$Outcome_simpler,
                                             levels=c("at light","up","down","other"))

#overall proportion of flights ending at the light:
OUTCOMES_CENTRAL_MULTI%>%
  group_by(FlighttoLight_tracked)%>%
  summarise(n=n())%>%
  ungroup()
15/47#30% flights ended at the light

#plot
SUMMARY_OUTCOME_CENTRAL_MULTI<-OUTCOMES_CENTRAL_MULTI%>%
  group_by(Treatment, Outcome_simpler, Year)%>%
  summarise(n=n())%>%
  ungroup()
SUMMARY_OUTCOME_CENTRAL_MULTI$Outcome_simpler<-factor(SUMMARY_OUTCOME_CENTRAL_MULTI$Outcome_simpler,
                                                    levels=c("other","down","up","at light"))

SUMMARY_OUTCOME_CENTRAL_MULTI$Treatment <-factor(SUMMARY_OUTCOME_CENTRAL_MULTI$Treatment,
       levels = c("SingleC_0.3lx","CL_0.3lx"))

supp_b2<-ggplot(SUMMARY_OUTCOME_CENTRAL_MULTI, aes(fill=Outcome_simpler, y=n, x=Treatment, label=n)) + 
  geom_bar(position="fill", stat="identity")+
  theme_bw()+ylab("Proportion of flights")+
  geom_text(size = 3, position = position_fill(vjust = 0.5))+
  labs(fill="Flight outcome")+
  xlab("Focal light intensity (lx)")+
 scale_x_discrete(labels=c("10", "30"))+
  scale_fill_manual(values = c("#B3B3B3","#66C2A5","#E78AC3","#FFD92F"))+
  geom_text(size = 3, position = position_fill(vjust = 0.5), col="grey30")+  theme(legend.text = element_text(size=10))+
  theme_classic()+
  ggtitle("F")+
  theme(axis.title.x = element_text(size = 12),
        axis.title = element_text(size = 14),
        axis.text.x = element_text(size = 10),
        axis.title.y = element_text(size = 12),
        axis.text.y=element_text(size=10))+
  #scale_x_discrete(drop = TRUE) +
  scale_y_continuous(expand=c(0,0))
supp_b2

#analysis - for flight-to-light only

central_mod2<-glmmTMB(FlighttoLight_tracked ~ Treatment+TrialNumber+CaptureMethod+
                      (1|LatinName)+(1|Family),
                    family=binomial, data=OUTCOMES_CENTRAL_MULTI)
summary(central_mod2)
central_mod2_resid <- simulateResiduals(central_mod2)
plot(central_mod2_resid)#ok

central_mod2b<-glmmTMB(FlighttoLight_tracked ~ Treatment+CaptureMethod+
                       (1|LatinName)+(1|Family),
                     family=binomial, data=OUTCOMES_CENTRAL_MULTI)

anova(central_mod2, central_mod2b)#remove trial number
summary(central_mod2b)
central_mod2c<-glmmTMB(FlighttoLight_tracked ~ CaptureMethod+
                       (1|LatinName)+(1|Family),
                     family=binomial, data=OUTCOMES_CENTRAL_MULTI)
anova(central_mod2c, central_mod2b)#remove treatment

central_mod2d<-glmmTMB(FlighttoLight_tracked ~ 1+(1|LatinName)+(1|Family),
                     family=binomial, data=OUTCOMES_CENTRAL_MULTI)
anova(central_mod2c, central_mod2d)#effect of capture method
summary(central_mod2c)

exp(1.800)#OR = 6.050

supp_c2<-plot_model(central_mod2c, type = "pred", 
                                  terms = c("CaptureMethod"),
                                  title = "G",
                                  colors = "darkgreen")+
  theme_classic()+
  ylab("Predicted probability of flight-to-light (%)")+xlab("Capture method")+
  theme(axis.title.x = element_text(size = 12),
        axis.title = element_text(size=14),
        axis.text.x = element_text(size = 10),
        axis.title.y = element_text(size = 12),
        axis.text.y=element_text(size=10))

#3)Erratic spiralling

OUTCOMES_CENTRAL_MULTI%>%
  group_by(Erratic_spiralling)%>%
  summarise(n=n())%>%
  ungroup()
13/47#28% flights with erratic spiralling 

#plot
SUMMARY_SP_CENTRAL_MULTI<-OUTCOMES_CENTRAL_MULTI%>%
  group_by(Treatment, Erratic_spiralling)%>%
  summarise(n=n())%>%
  ungroup()#only 3 in 10 lx group, 9 in 30 lx

SUMMARY_SP_CENTRAL_MULTI$Erratic_spiralling<-as.factor(SUMMARY_SP_CENTRAL_MULTI$Erratic_spiralling)
SUMMARY_SP_CENTRAL_MULTI$Treatment<-factor(SUMMARY_SP_CENTRAL_MULTI$Treatment, levels = c("SingleC_0.3lx","CL_0.3lx"))
supp_d2<-ggplot(SUMMARY_SP_CENTRAL_MULTI, aes(fill=Erratic_spiralling, y=n, x=Treatment, label=n)) + 
  geom_bar(position="fill", stat="identity", col = "#17A3FA")+
  theme_bw()+ylab("Proportion of flights")+
  geom_text(size = 3, position = position_fill(vjust = 0.5))+
  labs(fill="Erratic spiraling")+
  xlab("Focal light intensity (lx)")+
  scale_x_discrete(labels=c("10", "30"))+
  scale_fill_manual(values = c("white","#17A3FA"), labels = c("no","yes"))+
  geom_text(size = 3, position = position_fill(vjust = 0.5), col="grey30")+  theme(legend.text = element_text(size=10))+
  theme_classic()+
  ggtitle("H")+
  theme(axis.title.x = element_text(size = 12),
        axis.title = element_text(size = 14),
        axis.text.x = element_text(size = 10),
        axis.title.y = element_text(size = 12),
        axis.text.y=element_text(size=10))+
  #scale_x_discrete(drop = TRUE) +
  scale_y_continuous(expand=c(0,0))

#analysis - for flight-to-light only

central_mod3<-glmmTMB(Erratic_spiralling ~ Treatment+TrialNumber+CaptureMethod+
                        (1|LatinName)+(1|Family),
                      family=binomial, data=OUTCOMES_CENTRAL_MULTI)
summary(central_mod3)
central_mod3_resid <- simulateResiduals(central_mod3)
plot(central_mod3_resid)#ok

central_mod3b<-glmmTMB(Erratic_spiralling ~ Treatment+TrialNumber+
                         (1|LatinName)+(1|Family),
                       family=binomial, data=OUTCOMES_CENTRAL_MULTI)

anova(central_mod3, central_mod3b)#remove capture method
summary(central_mod3b)
central_mod3c<-glmmTMB(Erratic_spiralling ~ TrialNumber+
                         (1|LatinName)+(1|Family),
                       family=binomial, data=OUTCOMES_CENTRAL_MULTI)

anova(central_mod3c, central_mod3b)#remove treatment
central_mod3d<-glmmTMB(Erratic_spiralling ~ 1+(1|LatinName)+(1|Family),
                       family=binomial, data=OUTCOMES_CENTRAL_MULTI)

anova(central_mod3c, central_mod3d)#no effect of trial

#4)Tortuosity

#select only the ones for the pairs dataset
SUPP_CENTRAL_SEGMENTS<-SUPP_MULTITRACKS_SEGMENTS%>%
  filter(Unique_Flight_ID %in% OUTCOMES_CENTRAL_MULTI$Unique_Flight_ID)%>%
  droplevels()
str(SUPP_CENTRAL_SEGMENTS)#42 flights, all different moths

#Plot results

col_names2 <- c("Unique_Moth_ID","UniqueSegment","After_Light_Both_Exposed","Treatment","LatinName","Family","CaptureMethod","Location")
SUPP_CENTRAL_SEGMENTS<- SUPP_CENTRAL_SEGMENTS %>%
  mutate(across(all_of(col_names2), as.factor))
str(SUPP_CENTRAL_SEGMENTS)

treatment_counts <- table(SUPP_CENTRAL_SEGMENTS$Treatment)
treatment_counts_vector <- paste0("n=", as.numeric(treatment_counts))
treatment_counts

SUPP_CENTRAL_SEGMENTS$Treatment<-factor(SUPP_CENTRAL_SEGMENTS$Treatment, levels = c("SingleC_0.3lx", "CL_0.3lx"))
supp2_plot_tortuosity_beforeafter <- ggplot(SUPP_CENTRAL_SEGMENTS, aes(x = Treatment, y = mean_angle_3D, fill = After_Light_Both_Exposed)) +
  geom_boxplot() +
  theme_classic() +
  labs(title = "", y = "Mean turning angle (degrees)", x = "Focal light intensity (lx)") +
  scale_x_discrete(labels = c("10", "30"))+
  labs(fill="Flight segment")+
  scale_fill_discrete(labels = c("before \nexposure", "after \nexposure"))+
  theme(
    
    axis.text = element_text(size = 12),
    axis.title = element_text(size = 14),
    axis.title.y = element_text(size = 12),
    axis.title.x = element_text(size = 12),
    axis.text.x = element_text(size = 10),
    panel.grid.major = element_blank(),
    panel.grid.minor = element_blank(),
    panel.border = element_blank(),  # keep axis lines, so we don't add a full box
    axis.line = element_line(color = "black")  # this adds back the x and y axis lines
  ) +
  ggtitle("I")
supp2_plot_tortuosity_beforeafter #clear effect of exposure, but little difference between treatments otherwise

#statistics
  #singular fit with all random effects, and best with sqrt transform
central_mod4<- lmer(sqrt(mean_angle_3D) ~ Treatment*After_Light_Both_Exposed+(1|Unique_Flight_ID), data=SUPP_CENTRAL_SEGMENTS)
central_mod4_resi <- simulateResiduals(central_mod4)
plot(central_mod4_resi)#better with transform
hist(resid(central_mod4))
central_mod4b <- lmer(sqrt(mean_angle_3D) ~ Treatment+After_Light_Both_Exposed+(1|Unique_Flight_ID), data=SUPP_CENTRAL_SEGMENTS)
anova(central_mod4b,central_mod4) #remove interaction
summary(central_mod4b)
central_mod4c <- lmer(sqrt(mean_angle_3D) ~ After_Light_Both_Exposed+(1|Unique_Flight_ID), data=SUPP_CENTRAL_SEGMENTS)
anova(central_mod4b,central_mod4c) #remove treatment
summary(central_mod4c)
central_mod4d <- lmer(sqrt(mean_angle_3D) ~1+(1|Unique_Flight_ID), data=SUPP_CENTRAL_SEGMENTS)
anova(central_mod4d,central_mod4c) #effect of exposure alone

#plot all (Figure S3)
layout<-"
AABBC
DDEEE"
supp_to2+supp_b2+supp_c2+supp_d2+supp2_plot_tortuosity_beforeafter+plot_layout(design = layout)
```
