## Supplementary data and code for "Effects of artificial light colour, intensity, structure and contrast on moth flight behaviour": README.txt

README

#### KEY INFORMATION ####

Title of the study: Effects of artificial light colour, intensity, structure and contrast on moth flight behaviour

Short summary of the study: This study investigates how moth flight behaviour (including probability of taking off, flight outcomes and flight tortuosity) is affected by different background light levels and mid-flight exposure to LED lights of varying characteristics.

Code and package versions: R version 4.3.2
Packages - tidyverse(2.0.0), ggplot2(3.5.1), stringr(1.5.1), lubridate (1.9.3), patchwork(1.2.0), ggmosaic (0.3.3),
ggpattern(1.1.3), sjPlot (2.9.0), lme4 (1.1.35.1), DHARMa (0.4.6), glmmTMB (1.1.10), emmeans (1.1.10), multcomp (1.4.25)
Last updated: 18th March 2026

#### DATA AND CODE PROVIDED ####

Datasets used for analyses presented in the paper:
1) Experiment1_MothInfo.csv: trial data for experiment 1 (testing the impact of exposure to single lights varying in intensity and spectrum)
2) Experiment2_MothInfo: as above, for experiment 2 (testing impacts of focal light structure [single vs multiple] and background light levels)
3) Experiment1_AnalysedTracks.csv: track data for analysed flight paths in experiment 1
4) Experiment2_AnalysedTracks.csv: track data for analysed flight paths in experiment 2

Code:
1) Analysis_Script.R: R script containing code for all analyses & plots presented in the paper

#### DESCRIPTION OF VARIABLES IN DATASETS #####

1) Experiment1_MothInfo.csv:
	Each row represents a single trial in the experiment
Date: Trial date
MothID: ID assigned to individual moth at time of trial
CaptureMethod: How each moth was collected for the experiment - Net or Trap (light trap)
Location: Location moth was collected - Campus or Penryn (private gardens)
Treatment: Focal light treatment moth was exposed to - Control (no light), BW 30 lx, BW 3 lx, BW 0.3 lx, BA 30 lx, BA 3 lx, BA 0.3 lx,NA 30 lx, NA 3 lx, NA 0.3 lx (BW = white LED, BA = broadband amber LED, NA = narrowband amber LED)	
TrialNumber: Trial number for individual moth - 1 or 2 (always 1 in experiment 1, as each moth was only used once)
TakeOff: did the moth fly off within one minute? - 0 (no), 1 (yes), NA (trial excluded)
Unique_Moth_ID: Moth ID combined with year of experiment - attributes a unique label to each moth	
LatinName: Moth species	
Family: Moth family
Type: How moth was filmed - 2D (with a single camera, showing flight path in 2D only), or 3D (with stereo camera system, allowing 3D reconstruction of flight path)
Year: Year in which trial was carried out - 2024 or 2025	
JulianDay: Julian day for trial	
TrackedFlight_Outcome: initial assessment of where tracked flight path ended. NA if no flight, or 2D flight that did not cross the light beam, or flight that was not tracked (e.g. because it was clearly too short or did not cross the light path)	
Spiral_up: Did moth perform an upwards spiral in flight (observer assessment over entire trial) - 0 (no), 1 (yes), NA (as above)
Spiral_down: Did moth perform a downwards spiral in flight (observer assessment over entire trial) - 0 (no), 1 (yes), NA (as above)
Spiral_other: Did moth perform any other kind of spiral in flight (observer assessment over entire trial) - 0 (no), 1 (yes), NA (as above)
Spiral: Did moth perform any kind of spirals (i.e. 1 in any of the three spiral variables above) - 0 (no), 1 (yes), NA (as above)
Erratic_spiralling: Did moth perform behaviour defined as "erratic spiralling" - 0 (no), 1 (yes), NA (as above)
Outcome_simpler: Simplified classification of tracked flight outcome - up, down, at light, other, NA(as above)	
FlighttoLight_tracked: Did tracked flight end at the light apparatus - 0 (no), 1 (yes), NA (as above)
Down_tracked: Did tracked flight end down (on the floor or exiting frame downwards) - 0 (no), 1 (yes), NA (as above)
Up_tracked: Did tracked flight end with moth flying upwards - 0 (no), 1 (yes), NA (as above)

2) Experiment2_MothInfo.csv:
	As for Experiment1_MothInfo.csv, with the following differences:
Unique_Flight_ID: Unique ID for each trial, combining Unique_Moth_ID with treatment  
Treatment: Variable representing the combination of focal light structure and background light levels moth was exposed to - DL_0.03lx, DL_0.3lx, DL_3lx, DL_0.03lx, DL_0.3lx, DL_3lx, SingleC_0.3lx (where DL = multiple lights, CL = single light, SingleC = single light at 10 lx)

3) Experiment1_AnalysedTracks.csv:
	Track data for all analysed flight paths - each row represents a video frame
Unique_Moth_ID: Moth ID combined with year of experiment - attributes a unique label to each moth
A_X, A_Y: 2D coordinates of moth position, from camera A  	
A_Centre_light_X, A_Centre_light_Y: 2D coordinates of focal light (central, if multiple) in footage from camera A
B_X, B_Y, B_Centre_light_X, B_Centre_light_Y: as above, for camera B
Frame: Frame number for each analysed video
X,Y,Z: Raw 3D coordinates of moth position	
X_smooth, Y_smooth,Z_smooth: Smoothed 3D coordinates	
MothID: ID assigned to individual moth at time of trial
Year: Year in which trial was carried out - 2024 or 2025	
In_Cone_Both: Indicates whether moth position was classed as being inside the focal light beam, in footage from both cameras - 0 (no), 1 (yes)
After_Light_Both: Indicates whether moth has been exposed to the focal light beam at this point, in footage from both cameras  (i.e. this track has a 1 in "In_Cone-Both" at this current frame or in a previous frame) - 0 (no), 1 (yes)	
Date: Trial date	
LatinName: Moth species	
Family: Moth family
Date: Trial date
MothID: ID assigned to individual moth at time of trial
CaptureMethod: How each moth was collected for the experiment - Net or Trap (light trap)
Location: Location moth was collected - Campus or Penryn (private gardens)
JulianDay: Julian day for trial	
Treatment: Focal light treatment moth was exposed to - Control (no light), BW 30 lx, BW 3 lx, BW 0.3 lx, BA 30 lx, BA 3 lx, BA 0.3 lx,NA 30 lx, NA 3 lx, NA 0.3 lx (BW = white LED, BA = broadband amber LED, NA = narrowband amber LED)	
LightX3D, LightY3D, LightZ3D: 3D coordinates of focal light (central, if multiple)

4) Experiment2_AnalysedTracks.csv:
	As for Experiment1_AnalysedTracks.csv, with the following additions:
Unique_Flight_ID: Unique ID for each trial, combining Unique_Moth_ID with treatment  
TrialNumber: Trial number for individual moth - 1 or 2 
